# A phylogenetic protein-coding genome-phenome map of complex traits across 224 primate species

**DOI:** 10.1101/2025.09.08.674744

**Authors:** Alejandro Valenzuela, Fabio Barteri, Claudia Vasallo, Joseph Orkin, Lukas Kuderna, Jean Boubli, Amanda Melin, Hafid Laayouni, Kyle Farh, Jeffrey Rogers, Tomàs Marquès-Bonet, Gerard Muntané, Arcadi Navarro, David Juan

## Abstract

Complex traits arise from networks of coding and regulatory loci, making it difficult to resolve their genetic basis. Macroevolutionary studies leverage tens of millions of years of divergence across species to uncover fixed genomic changes invisible to within-species approaches, such as GWAS, offering a complementary framework for generating hypotheses in biomedical research. Here, we present the first phylogenetic protein-coding primate-wide genome–phenome map (P3GMap), spanning 200 curated traits across 224 primate species, which we release through the Primate Genome-Phenome Archive (PGA, https://pgarchive.github.io). Using two complementary approaches, convergent amino acid substitutions and relative evolutionary rates, we linked protein-coding variation to complex phenotypes and identified thousands of candidate gene-trait associations, including lineage-specific signals related to diet, immunity, and lifespan.

**One sentence summary:** Cross-species genome-phenome mapping in primates reveals thousands of protein-coding variants linked to the evolution of complex traits.

## 1. Introduction

Complex traits arise from the interplay between genetic and non-genetic factors. Genetic contributions are typically distributed across interacting coding and regulatory elements. This multilayered organization makes it difficult to resolve the genetic basis of phenotypic variation. Much of our knowledge on the genetic architecture of complex traits originates from studies of model organisms (1–3) and human genome-wide association studies (GWAS) (4–7), which link genomic and phenotypic variation within populations. Although powerful, GWAS typically tag genetic markers rather than the causal variants themselves, capture only segregating variation within contemporary populations, and miss fixed substitutions along evolutionary lineages, changes that often underlie major phenotypic shifts and whose study can reveal loci of biological interest (8–9).

Macroevolutionary analyses address this gap by testing whether genetic elements are associated with traits across different species. Phylogenetic Comparative Methods (PCMs) (9–12) exploit millions of years of divergence to reveal associations with a wide variety of genomic variation, including gene gain and loss, rates of protein evolution, presence or absence of structural variants, and point mutations (13–16). PCMs have revealed the genetic architecture of key traits in mammalian evolution, such as body size (17–21), brain size (22–23), and maximum lifespan (24–27).

The growing availability of high-quality genomes enables scalable cross-species genome–phenome mapping. Leveraging new whole-genome sequencing data from 233 primate species (28), we created the first phylogenetic protein-coding genome–phenome map (P3GMap), which is publicly available through the Primate Genome–Phenome Archive (PGA, https://pgarchive.github.io). This resource integrates 200 traits across 224 primates with phenotypic data and reveals thousands of coding variants associated with phenotypic evolution, a number that exceeds by at least one order of magnitude the candidate associations typically reported in individual GWAS studies (29).

To illustrate the scope of the resource, we examined three case studies: insectivorous diet (30–33), white blood cell count (34–35), and maximum lifespan (24–27). For each of these traits, we defined phenotypic *Contrasts* as pairs of species from close lineages that show marked differences in trait values, and we used these to identify genes and individual substitutions consistently associated with the trait. These case studies exemplify how researchers can use the PGA to move from a trait of interest to candidate coding changes and pathways that may underlie the evolution of traits.

## 2. Results

### 2.1. The Primate Phenomic Space

We compiled phenotypic data for 224 of the 233 primate species with available whole-genome sequencing data (28), integrating the species mean values for 263 quantitative and qualitative traits derived from an initial set of 835 trait variables obtained from public sources (30,36–38). These traits were classified into five primary domains (behavior, 34 traits; ecology, 8; life history, 40; morphology, 114; and physiology, 67) and 15 secondary domains (Fig. 1, Methods, Table S1), thereby defining the primate phenomic space for comparative analyses. As data density varies across clades, comparisons should be constrained to sets of species with similar levels of trait information to avoid biases driven by uneven sampling. To do so, we used Multiple Correspondence Analysis (MCA) on a binary presence/absence matrix of trait data, followed by clustering in the main components to identify traits with similar patterns of species representation. This procedure resulted in 13 clusters of traits that shared comparable species coverage, allowing analyses to be conducted within subsets of traits measured in largely overlapping sets of species (Fig. 1, SM 1.3). Body mass-related traits were among the most consistently represented, explaining between 15 and 23% of the variance in the first component for clusters VIII–X (Fig. 1A). To control for allometry (39–43), we corrected 74 morphological, 14 life history, and two physiology traits for *Adult Body Mass* or for the more relevant body part size trait (SM 1.3, Data S1). After filtering out lower-quality and semantically redundant traits, we defined a final set of 200 traits for the phylogenetic genome-phenome association analyses. The curated dataset is publicly available from the Primate Genome–Phenome Archive (PGA, https://pgarchive.github.io).

**Figure 1.**
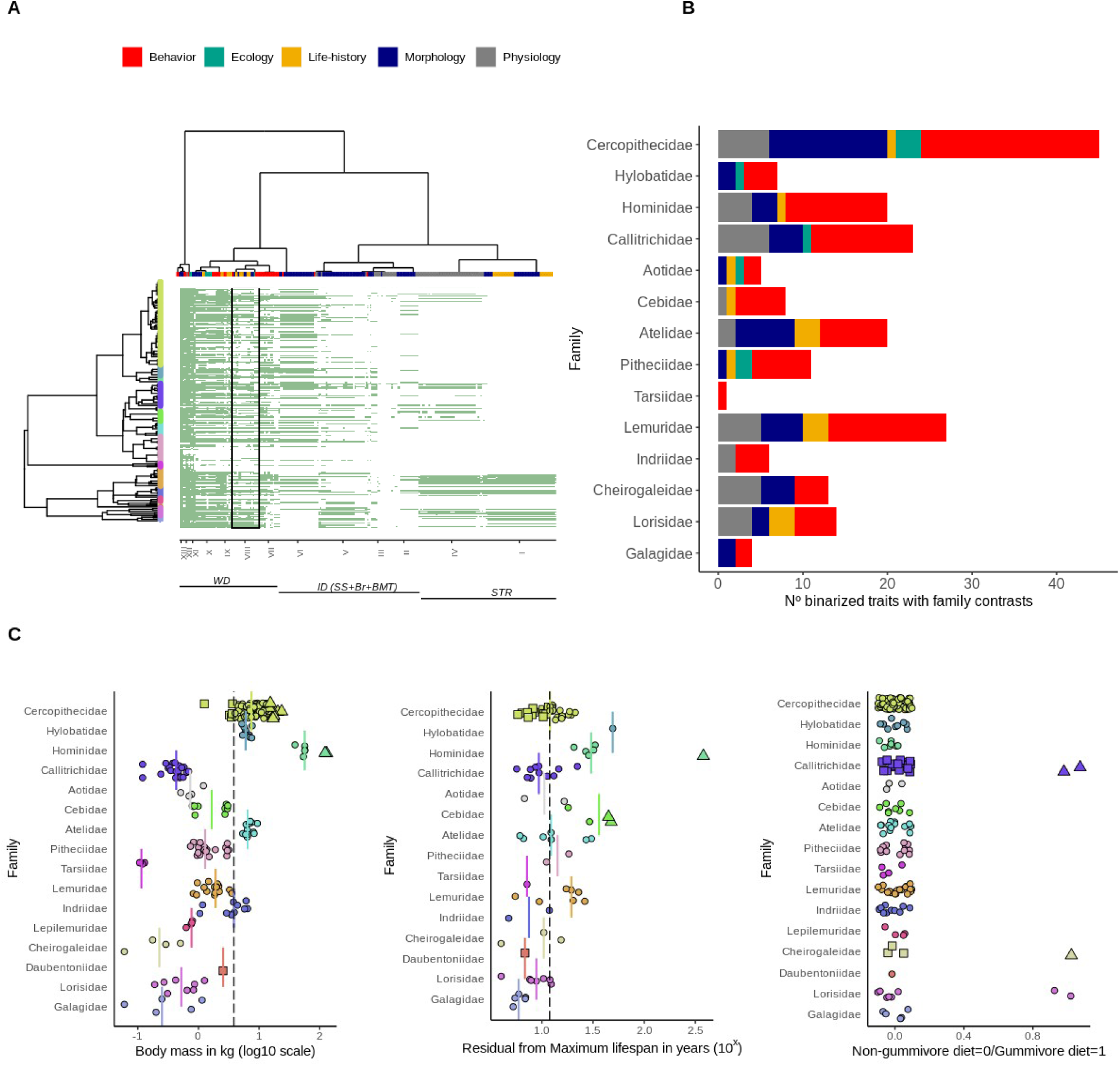
Overview of Primate Phenomic Data. (**A**) Hierarchical clustering of phenomic data mapped onto primate phylogeny and grouped by primate family. Based on data availability, the traits were classified into 13 data clusters [see Supplementary Material, Trait clustering by phylogenetic sampling]. The area within the square highlights traits from one of the clusters, Cluster VIII, which includes a broad range of ecological, life history, and morphological traits. ( **B**) Total number of traits (x-axis) showing extreme phenotypic differences between species within the same family. Top and Bottom values refer to species with contrasting trait values (see text for definitions). ( **C**) Examples illustrating the distribution of values for three traits: Body Mass (left), Maximum Lifespan (center), and Gummivore, sap-eating, Diet (right). Each point represents a species. Vertical bars indicate the median family values. Dashed lines correspond to the median global trait values. Species selected as Top and Bottom extremes (based on the criteria explained in the text) are indicated by triangles and squares, respectively.

We conducted a genome-phenome analysis of protein-coding sequences to create the first P3GMap (also available on the PGA web). This resource catalogues associations between protein-coding genes and diverse phenotypes across primates. The map comprises alignments for 16,133 clusters of one-to-one orthologous genes, over 75% of which include more than 200 species (Fig. S1, SM 2). These clusters of orthologous genes were retrieved as sets of best-bidirectional hits between 59 high quality reference genomes (21) and expanded to the full set of 224 primate whole-genome sequenced species using their mappings to the closest high quality reference genomes (28). Multiple sequence alignments were built and curated for these clusters of orthologous genes using codon-based approaches ((44–46), SM 2.2). To detect genetic factors that correlate with phenotypic evolution, we applied two conceptually distinct approaches: Convergent Amino Acid Substitutions (CAAS) using CAAStools (40; SM 3.1.2), and relative evolutionary rates (RERs) using RERconverge (39; SM 3.1.1). These methods capture genome-phenome associations across a broad range of evolutionary scenarios beyond classical positive selection signatures (9,11,13). To ensure consistency and scalability, we implemented a standardized strategy for cross-trait comparisons, providing a unified framework for mapping genome-phenome relationships in primates.

### 2.2 Primate Protein-coding Genome-Phenome Analyses

#### 2.2.1 Convergent Amino Acid Substitutions Associated with Phenotypic Diversity

For each trait, we defined two groups of species with contrasting trait values: Top and Bottom for continuous traits or distinct categories for discrete traits. These groups were constructed by selecting phylogenetically close species within the same clades (e.g., families), such that each species was paired with a closely related counterpart in the opposing group. This design results in the combination of cases in different families of intra-family pairs of species with drastically different phenotypes and increases the power to detect convergent amino acid substitutions associated with trait differences while controlling for phylogenetic distance.

CAASs were then identified from these pairwise *Contrasts* (see Fig. 1B–C, Figs. S18–S20, Data S1, and SM 3.1.2). To ensure independence, the standard strategy emphasizes *Contrasts* within families. Of the analyzed traits, 83 (37%) showed at least one intra-family contrast, and 63 showed at least one contrast in two or more families. Traits in widely distributed clusters (WD) exhibited multiple *Contrasts* (∼23%). Intermediately distributed (ID; ∼60%) and Strepsirrhine-centered (STR; ∼54%) clusters showed a high fraction of traits without *Contrasts*, reflecting both coverage limitations and the predominance of inter-family over intra-family variation in quantitative traits.

To apply the CAAS and RER methods, we constructed a dataset of curated multiple sequence alignments of one-to-one orthologs extracted from a recently sequenced set of 233 primate genomes (28), with 224 of these primate species having phenotypic information for at least one of the 200 curated traits. Using CAAStools for these alignments with phylogenetic correction via permulations (47) and per-trait multiple testing (SM 3.1.2.4), we detected 29,155 significant CAASs in 8,780 genes across 76 of the analyzed traits (p_permul-FDR_ < 0.05). This yielded a median of 49 genes and 51 positions per trait (Fig. 2A, Figs. S21–S24, Data S2), providing a large-scale resource for studying candidate molecular convergence in the future. These associations reflect fixed amino acid substitutions accumulated along evolutionary lineages, rather than segregating variation within species.

**Figure 2.**
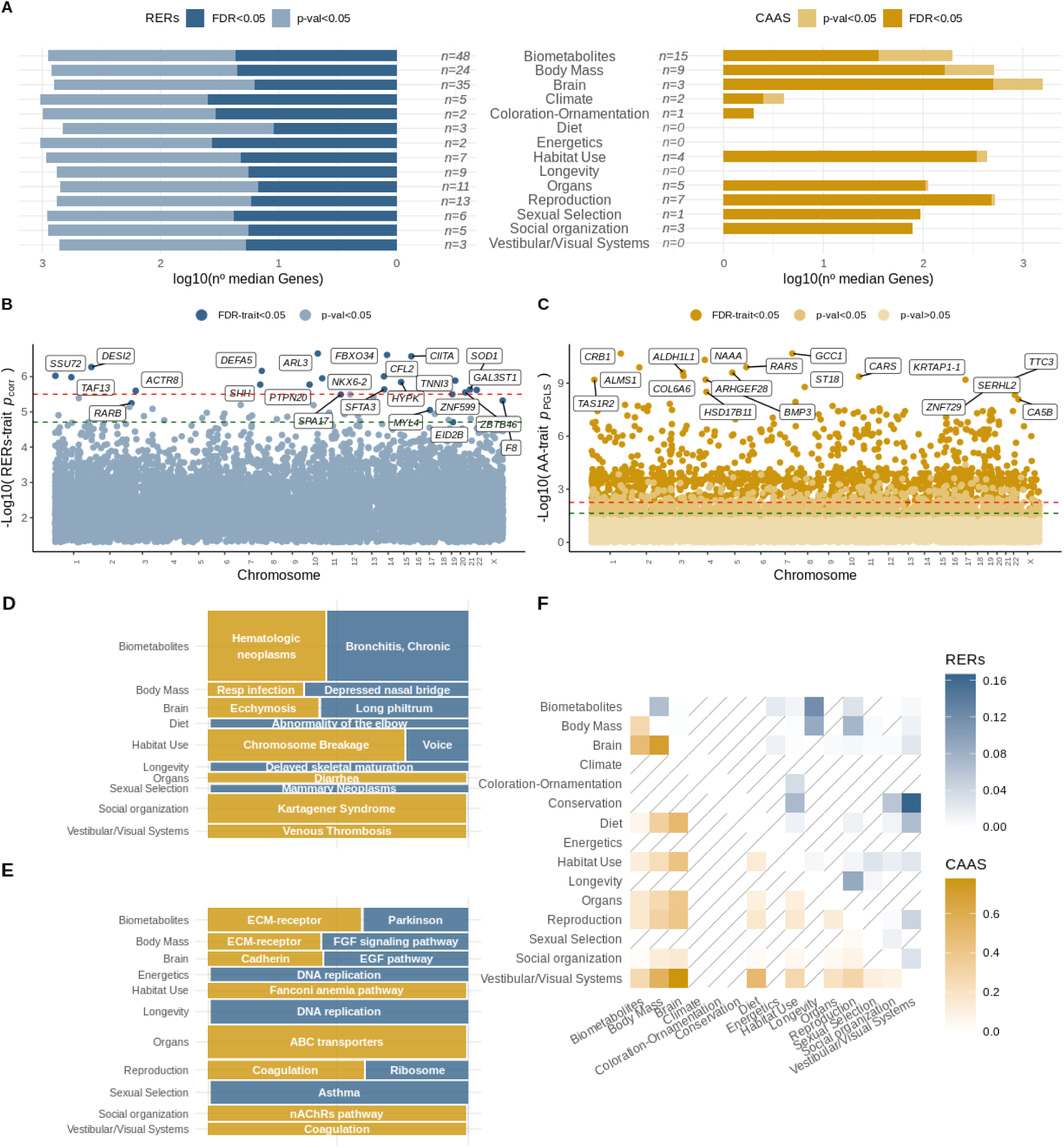
Exploring the Phylogenetic Protein-coding Genome–Phenome Map (P3GMap). (**A**) Gene–trait associations recovered across secondary biological domains using RERs (left) and CAAS (right), showing the median number of significant genes per trait (log10 scale) and the number of traits with associations (center). (**B**, **C**) Manhattan plots highlighting the top gene–trait associations detected by RERconverge and CAAS, with significance thresholds indicated (green, FDR; red, Bonferroni). Genes surpassing strict thresholds are labeled. (**D & E**) Functional enrichment identified by CAAS and RER across secondary domains, highlighting pathways recurrently implicated by each approach. (**F**) Heatmap showing the overlap of significant gene sets across secondary domains, with strip-lined boxes marking domain pairs without shared associations.

We assessed the consistency of these findings by testing whether the CAAS–trait associations identified in the discovery set could be recovered in an independent set of primate species not used during the discovery phase. This consistency check used phylogenetic generalized least squares (PGLS) for quantitative traits and phylogenetic logistic regression (phyloglm) for qualitative traits (see SM 3.1.2.5; Fig. 2C, Fig. S32). In this context, “primate consistent associations” are CAAS signals that remain significant in this independent dataset. This approach is conservative because it excludes the extreme phenotypic species used in the discovery step. To reduce the influence of non-specific or noisy signals, we selected representative biological examples from the most robust CAAS–trait relationships, defined as those significant after FDR correction in both PGLS and phylogenetic logistic regression (p_PGLS-FDR_ < 0.05, p_phyloglm-FDR_ < 0.05). We considered only the most informative traits, requiring at least five species in both the Top and Bottom groups and at least one intra-family *Contrast*. This approach recovered both known signals and novel candidate associations (Table S5).

Representative examples of CAAS–trait associations showing the strongest phylogenetic statistical support are listed below; these represent candidate links between amino acid variation and phenotypic traits across the phylogeny, but they do not imply direct causal effects of the genes on the associated traits. These include associations between *Body Mass* and the mitochondrial genes *NDUFS7* (substitutions Y223L, R230H, K231G, I232A; p_PGLS_ = 1.37 × 10⁻⁵–3.57 × 10⁻⁵) and *NOXO1* T/V223A (p_PGLS_ = 2.06 × 10⁻⁷); the gene of the immunoglobulin family *ICAM1* R/V418M (p_PGLS_ = 2.39 × 10⁻⁵) (48); and the association of *Lactation Period Length* with *MEFV* K/R546Q (p_PGLS_ = 1.66 × 10⁻⁶), an innate-immunity and familial Mediterranean fever gene affecting breastfeeding outcomes (49–51) and expressed stage-dependently in the lactating mammary gland. Additionally, we tested ten equivalent traits in non-primate mammals to determine whether their 4,968 CAASs exhibited consistent effects in mammalian phylogeny. Despite being highly conservative, this test yielded far more consistency than expected by chance (p_binomial_: 4.07×10e−^6^, see SM 3.1.2.7, Suppl. Table S6). Representative cross-mammalian consistent examples include *NELL1* Q176R for *Lactation Period Length* (p_PGLS_ = 1.09 × 10⁻⁴), which has been previously associated with delayed milk production in goats (52) and cranial bone development (53–55), and *MPDZ* H/P956Q for *Gestation Period Length* (p_PGLS_ = 1.65 × 10⁻⁵), a gene whose mutations cause congenital hydrocephalus (56).

#### 2.2.2 Relative Evolutionary Rates and Trait Associations

While CAASs detect genome–phenome associations by focusing on specific AA differences between species with the most different trait values, relative evolutionary rates (RERs) identify genes whose entire sequence evolutionary rates covary with trait changes. None of the methods implied the presence of positive or purifying selection. We applied RERconverge to 16,109 multiple sequence alignments of orthologous coding genes (24 alignments were skipped due to technical problems) and 200 quantitative traits (see SM 3.1.1), assessing significance using permulations (41; SM 3.1.1.1). This analysis yielded 3,911 genes significantly associated with 194 different traits (p_permul-FDR_ < 0.05), with a median of 19 genes per trait, summarized across domains (Fig. 2A, Figs. S21–S24, Data S2). Because RERconverge applies less conservative corrections than CAAStools (SM 3.1.1), we also required FDR-corrected parametric p-values, which were broadly consistent with permulation statistics (SM 3.1.1.1).

RERconverge revealed biologically relevant signals (SM 3.1.1, Table S3), including *NKX6-2* for *Percentage of Time Moving* (Rho = 0.62; p_permul-FDR_ < 10⁻³), which has been linked to spastic ataxia (57–58), or *DEFA5* for *Percentage of Fauna Ingest* (Rho = –0.5; p_permul-FDR_ < 10⁻³), which is a defensin gene associated with gastritis and gastric cancer (59–60).

### 2.3 Functional analysis of the P3GMap

We performed over-representation analyses to evaluate the biological functions associated with the identified P3GMap genes. Genes with significant CAAS were associated with 199 unique enriched terms across 27 traits, each exhibiting at least one enrichment (Suppl. Data S3). Body mass–related traits were enriched for DNA repair/replication/recombination categories, consistent with previous findings in Carnivora (61) and with the identification of DNA repair genes as anti-cancer targets in long-lived species (27,62–65) (Suppl. Table S7). The trait *Percentage of Time with Social Activity* showed enrichment for chromosome breakage (ER = 7.21; p = 4.53 ×10⁻⁷) and Fanconi anemia (ER = 4.73; p = 3.03 ×10⁻⁵), which are linked to developmental disorders (66–67) (Fig. 2D-E).

Across 48 traits, 94 enriched terms were found for the significant RERconverge genes. These included *Long Philtrum* (distance >2SD between nasal base and upper lip), which is associated with *Pallidum* (ER = 3.35; p = 2.97 ×10⁻⁶) and *Subthalamus Size* (ER = 3.34; p = 3.12 ×10⁻⁶). Nose morphology is a key distinguishing feature between strepsirrhines and haplorhines (68–69) and has been linked to human syndromes involving cognitive, sensory, and craniofacial abnormalities (70–74). Traits under sexual selection also showed enrichment: *Hair Dimorphism* with the MHC complex (p = 7.76 ×10⁻⁶), which has long been proposed to affect mate choice (75–79), although this remains debated in some taxa (80–84); and *Body Mass Dimorphism* with mammary neoplasms (ER = 2.16; p = 4.65 ×10⁻⁶), which is consistent with the effects of sex-dimorphism on cancer incidence (85–87).

The examination of trait relationships further refined the functional interpretation. Gene set overlaps across domains (Fig. 2F) revealed 16 significant pairs for CAAS (p < 1 ×10⁻⁵⁰), concentrated in *Trichromatic Color Vision* and *Brain/Body Mass*, suggesting pleiotropic effects across sensory and morphological systems. Five overlaps involved life history traits, such as *Age of Female at First Reproduction* with *Trichromatic Color Vision* and with *Maximum Young Adult Weight*; and *Gestation Period Length* with *Maximum Young Adult Weight* (including sex-specific values). These reproductive overlaps were prominent in Atelidae, where *Ateles* and *Lagothrix* show polymorphic color vision (88), and Alouatta displays routine trichromacy (89), suggesting a possible coordinated evolution of vision, reproduction, and somatic traits under ecological and social pressures.

RERconverge overlaps (11 pairs, p < 10⁻³⁰) emphasized longevity-related traits. For example, gene sets associated with the Percentage of Infant Mortality overlapped with those associated with Maximum Male Longevity and Gamma-Glutamyl Transferase Levels (GGT) (p = 4.46 ×10⁻⁷⁴, p = 7.13 ×10⁻³⁹), an enzyme central to glutathione catabolism that is widely used as a biomarker of oxidative stress and is linked to cardiovascular and neurological diseases (90–96), lung inflammation (97), cancers (98–100), and mortality prediction (101). These overlaps do not imply that the same genes directly affect multiple traits but may reflect shared biological pathways, correlated life history traits, or the underlying phylogenetic structure. Similarly, gene sets associated with Cholesterol Levels overlapped with those linked to Median Male Longevity (p = 6.74 ×10⁻⁴⁸), although cholesterol–mortality links remain debated (102). Gene sets associated with Serum Oxaloacetic Transaminase Levels (AST) overlapped with those linked to Median Female Longevity (p = 2.18 ×10⁻³⁸); elevated AST is associated with mortality in liver, neurological, and cardiovascular diseases (103–106).

Family-level analyses revealed lineage-specific patterns. *Cathemeral Activity* (sporadic intense activity periods during day and night) shared gene sets with *Dichromatic Color Vision* (p = 7.72 ×10⁻⁴⁹) and *Seasonal Breeding* (p = 2.34 ×10⁻²⁰⁰), mainly in Lemuridae. The trait *Solitary Social System* overlapped with *Monochromatic Color Vision* (p = 2.98 ×10⁻⁴¹) in Lorisidae and Galagidae, and with *Dichromatic Color Vision* (p = 9.85 ×10⁻⁴⁸) in Lepilemuridae, *Microcebus*, and *Nycticebus*. These patterns are consistent with the associations between sensory ecology, activity patterns, and social behavior across primates, particularly in relation to nocturnality and ecological niche differentiation. However, differences in the evolutionary history of visual systems across clades suggest that these relationships may reflect a combination of shared ancestry and ecological factors rather than uniform co-evolutionary processes.

Finally, we investigated whether P3GMap-associated genes overlapped with human GWAS loci. We did not detect significant enrichment, which is compatible with, but does not establish, a limited overlap between cross-species and within-population variation (SM 6.1–6.2). Moreover, 86.16% of CAAS positions are fixed in humans, consistent with these sites capturing divergence between species rather than polymorphisms within them. This suggests a decoupling between macroevolutionary divergence and microevolutionary variation, reinforcing the complementarity of cross- and within-species approaches in disentangling the architecture of complex traits.

### 2.4 Targeted Case Studies of Primate Traits

While the general analysis in the P3GMap framework enables consistent cross-trait comparisons across hundreds of phenotypes with RERconverge, CAAS, and other methods, tailored lineage-specific CAAS analyses can reveal mechanisms that are obscured in large-scale scans. To demonstrate this, we examined three unrelated traits from different domains: *Insectivorous Diet*, *White Blood Cell Count*, and *Maximum Lifespan*. First, *Diet* (a widely distributed trait) displayed one *Contrast* for insectivory, reflecting the propensity of broadly distributed, often qualitative traits to capture intra-family extremes. Second, *White Blood Cell Count* (a strepsirrhine-focused trait) presented two *Contrasts*, providing an example within a cluster that generally yielded fewer contrasting cases. However *Maximum Lifespan* showed no *Contrasts*, providing an example of a quantitative trait whose variation tends to manifest between families rather than within them.

In these three case studies, a larger set of candidate species with extreme trait values was available beyond those used in the main discovery framework from P3GMap, enabling alternative species selections and allowing us to further test the flexibility of the CAAS approach under expanded phylogenetic sampling. By tailoring species *Contrasts* to the phylogenetic distribution of each trait, we identified additional gene sets that refined the functional interpretation and highlighted recurrent adaptive pathways. Thus, the case studies (Fig. 3A–C) demonstrate the potential of the PGA for trait-focused exploration, which complements the global view of the general analysis presented above. Additionally, to leverage the biological relevance of our approach, we performed a natural selection analysis on these positions, revealing directional selection toward specific amino acids, hereafter referred to as “directional selection” for simplicity (SM 3.2)

**Figure 3.**
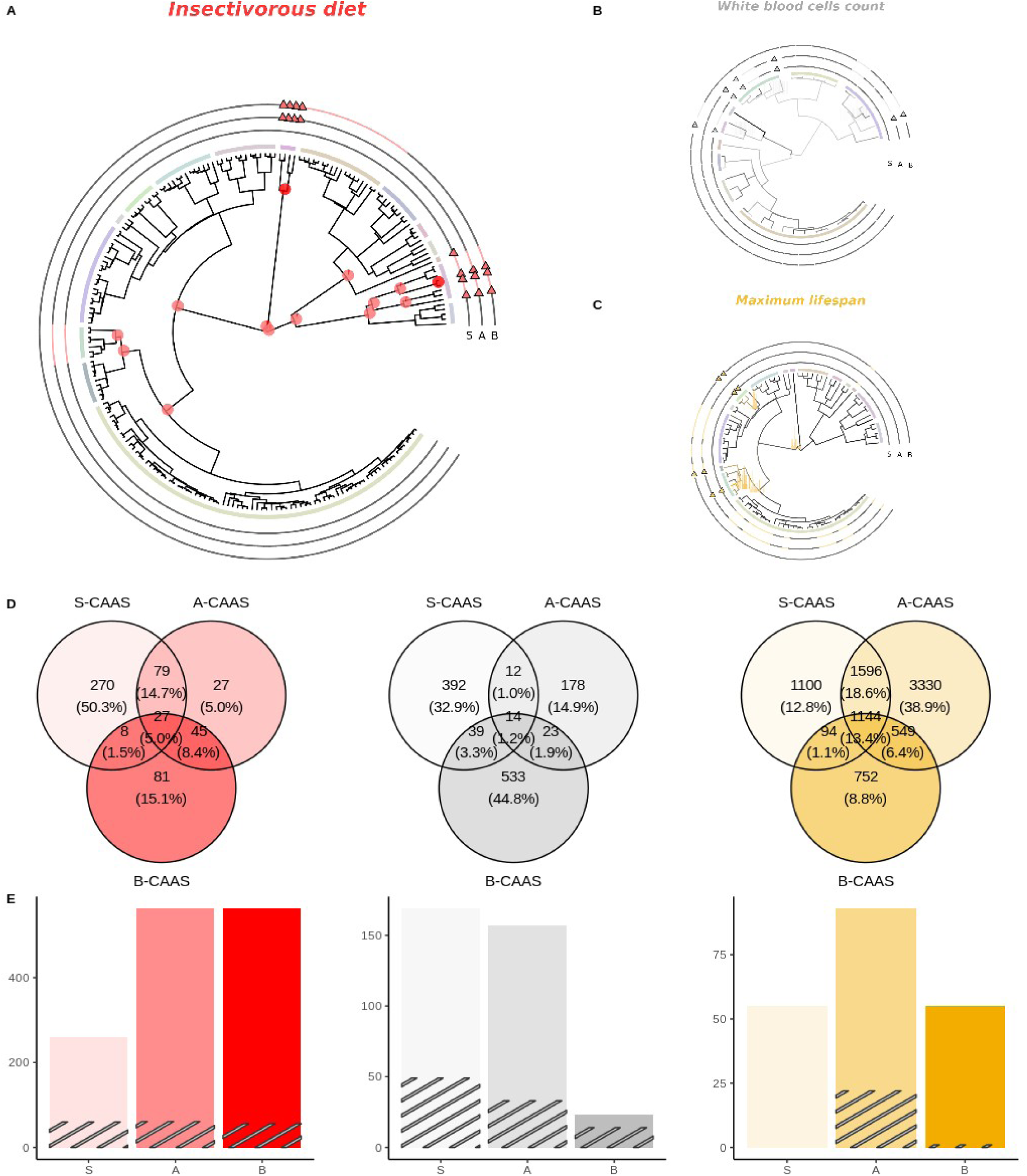
Case studies illustrating the CAAS selection approach for Insectivorous Diet, Maximum Lifespan, and White Blood Cell Count. (**A–C**) Selection of Top species (triangles) and Bottom species (colored lines) using the standard genome–phenome strategy (*S-*) and two alternative approaches (*A-* and *B-*). (**D**) Venn diagrams of gene overlap (p-perm < 0.05) between alternative family selections for each trait. (**E**) Number of sites under directional selection per trait; stray lines indicate CAAS positions showing positive directional selection.

#### 2.4.1 Insectivorous Diet

Diet, defined here as the proportional contribution of major food categories such as insects, fruits, or leaves based on field observations and literature-derived estimates, is closely associated with traits such as Brain Size (31–33) and Dentition (107–108) in primates, including strepsirrhines (33). For this case study, species were classified as insectivorous according to the ecological data compilation of Galán-Acedo *et al*. (37), in which insectivory corresponds to diets with a dominant arthropod component, typically above 50% when quantitative data are available, or to equivalent qualitative evidence from the literature. Under the standard strategy, the *contrasts* involved Lorisidae (slender versus slow lorises) and *Daubentonia madagascariensis* (Daubentoniidae) as insectivorous species compared with non-insectivorous species from other families. Species with more variable or mixed insect consumption were not included in this *Contrast* to maintain a conservative and comparable definition across distantly related lineages. In the tailored design, we included Tarsiidae as an additional insectivorous lineage alongside alternative non-insectivorous selections (Fig. 3A). This approach identified 619 positions across 537 genes, all of which were significant by permulations (Fig. 3D). These were enriched in four categories (pFDR < 0.05): apical part of the cell (GO:0045177, 19 genes), cargo receptor activity (GO:0038024, 8 genes), lipase activity (GO:0016298, 7 genes), and endopeptidase activity (GO:0004175, 14 genes). These categories were not recovered when the standard *Contrast*, the Tarsiidae-inclusive *Contrast*, or the alternative non-insectivorous comparison sets were analyzed separately, indicating that they emerged only when the fully tailored set of insectivory-related *Contrasts* was considered together. These enrichments suggest that the epithelial surface is a hub for nutrient absorption, consistent with the structural changes implicated in nutritional and inflammatory disorders (109–114). Among the candidate genes, Sucrase–Isomaltase (SI) carried CAASs A/R/T917K across all species sets and I627M in those including Tarsiidae, both under directional selection. SI is essential for carbohydrate digestion, and loss-of-function variants in Arctic human populations cause sucrose intolerance in childhood (115–117). Lipases also featured prominently, including ASPG (p.G374D), HPSE (A/F/L44V), PLA2G3 (L432P), and PNLIP (K394R), all of which were subjected to directional selection (118–119). Of the 27 genes and 14 positions shared across selections, DEFB113 V52I was the only site under directional selection in all the comparisons. DEFB113, a β-defensin gene central to host defense, has undergone repeated gains and losses in mammals (120–122), with diet proposed as a driver of this diversification (123–125).

#### 2.4.2 White Blood Cells Count

*White Blood Cell Count* (WBC) metrics have rarely been explored in macroevolutionary analyses, although they correlate with mating systems and sexually dimorphic traits, such as *Testis Size* (34–35). We contrasted species with extreme WBC counts from Hominidae and Lorisidae (standard strategy) and added Callitrichidae, which present outlier values. This identified 1,901 positions in 1,191 genes with significant permulation results. Enrichments included olfactory receptor activity (GO:0004984), also recovered in the general P3GMap analysis above, and two novel extracellular matrix (ECM) categories: ECM constituent (GO:0005201) and ECM–receptor interaction (hsa04512). These ECM categories partially overlapped with olfactory receptor genes (9 and 6 genes, respectively) but were only detected in the analysis tailored for the case study. Olfactory receptors (ORs), canonical sensory genes, are also expressed in immune cells, such as macrophages (126), and leukemia cell lines (127–129). Three ORs were particularly noteworthy: *OR4D5* (I48T) under directional selection in Hominidae; and OR52B6 and OR10AD1, the latter with L/M102I under directional selection. Although ORs constitute a large multigene family with a complex history of duplication that can complicate read mapping and generate spurious signals of sequence divergence, these three ORs are largely unduplicated across primates, have no close paralogs in Ensembl (130), and reside at distinct genomic loci. This pattern is consistent with independent, repeated co-option of ORs for immune and sensory functions. We also detected a xenobiotic metabolism cluster. *SULT1C3* carried four CAASs (K45R, H/R196Q, E227D, and Q243Y) under directional selection in Callitrichidae and Hominidae, plus 19 further associations (pPGLS < 1 × 10⁻³). CES2 showed a CAAS (L599F) with directional selection and consistent phylogeny-wide association (pPGLS = 5.46 × 10⁻⁴). Both genes are central to detoxification and belong to neuro-immune pathways involved in xenobiotic avoidance, allergies, and immune dysregulation (131–133). Immune recognition genes also showed lineage-specific signals: LILRB3 under directional selection in Hominidae, LILRA6 under selection despite lacking CAASs, HLA-DQA1 under directional selection in Hominidae, and HLA-DPB1 with a CAAS (Q75R) in Callitrichidae and Lorisidae (pPGLS = 1.67 × 10⁻⁴). Collectively, these results underscore the interplay between immune function, environmental exposure, and lineage-specific adaptation in shaping WBC variation.

#### 2.4.3 Maximum Lifespan

Longevity is a central focus of comparative genomics, with humans living markedly longer than their closest relatives (62,134–136). After correcting for *Body Mass*, the *Contrasts* used in the general analysis we performed for the P3GMap included humans (Hominidae), long-lived Cebidae, and several short-lived lineages. For the tailored use case, we refined these comparisons by adding Hylobatidae as another long-lived clade and including additional short-lived species (Fig. 3C). This analysis identified 30,886 positions in 8,565 genes (Fig. 3D). However, because lifespan *Contrasts* often span entire families, the initial set of associations is likely to include a higher proportion of nonspecific signals. To refine this, we assessed the consistency. In primates, from 62.72% of informative positions, defined as those containing Top and Bottom states across more than five additional species, the analysis yielded 5,225 positions in 3,338 genes with consistent associations. In mammals, this figure was 30.71% of informative positions, yielding 1,061 positions in 921 genes with consistent associations across primate and nonprimate mammals. These curated sets provide a robust basis for functional interpretation. First, enrichment analysis of primate-consistent revealed four categories, including chromosome segregation (GO:0007059) and complement/coagulation cascades (hsa04610). Eight categories emerged for mammal-consistent genes, including interleukin-10 production (GO:0032613) and regulation of multi-organism processes (GO:0043900). Across both clades, 139 amino acid positions in 136 genes were consistently associated with and enriched in cranial nerve paralysis (HP:0006824) and abnormal cranial nerve physiology (HP:0031910), conditions linked to diabetes (137), metabolic syndrome (138), ischemic (small-vessel) origin (139), or cerebral aneurysm (140). These enrichments should be interpreted as shared functional annotations within the associated gene sets, rather than as direct evidence that cranial nerve-related pathology mediates lifespan evolution. From the gene sets linked to these terms in this case study, seven genes overlapped in the general strategy, highlighting the added value of examining the overlaps across different selection configurations.

When we compared all strategies using the most phylogenetically restrictive set of consistent genes, six genes showed CAASs across all species selections. Notable examples include IL1B with CAAS L152M, a key regulator of immune and inflammatory responses (141–143), complemented by the detection of two other interleukins, IL12B and IL4I1. Additionally, we recovered IL23R and IL5 in the directional selection analysis. A second example is *KLK4* Q227P (Kallikrein-related peptidase 4), which exerts tumorigenic effects in prostate cancer linked to the IGF axis through cleavage of IGF-binding proteins (88), detected in comparative studies searching for genomic correlates of long-lived species (27). We also detected CAASs and directional selection in KLK1 P57R in Hylobatidae and Hominidae, although this site was not consistently associated across primates or mammals. Together, these results implicate immunity, inflammation, and growth factor pathways as recurrent contributors to lifespan evolution.

## 3. Discussion

This study presents the first P3GMap, which integrates 200 traits across 224 primate species. By combining CAAS (144) and RERs (145), we uncovered thousands of candidate associations linking coding variation to phenotypic evolution. The PGA and the associated P3GMap offer a unified resource for studying the genetic basis of traits related to ecology, morphology, life history, and physiology. This approach complements within-species studies, such as GWAS (4–7), by capturing evolutionary signals that are fixed along lineages (8–9), as illustrated by the observation that approximately 86% of CAAS positions are fixed in humans. This reinforces the decoupling between macroevolutionary divergence, which establishes species-specific trait differences, and microevolutionary variation, which can be readily observed in within-species studies (27,62–65).

Our framework has several limitations. It focuses on single-copy protein-coding genes (28), excluding regulatory variation, gene duplications, and structural variants, all of which are also important for trait evolution (146). CAAS and RERconverge rely on distinct statistical and conceptual foundations, which can result in partially different gene–trait associations (144–145). Although permulation tests are rigorous (47), they are based on Brownian motion models and may not fully capture all evolutionary dynamics (11,147–148). These caveats highlight the need for future expansions that incorporate regulatory, structural, and expression data (146), and refined evolutionary models (11,147–148).

Despite these constraints, our results demonstrate the power of macroevolutionary analyses in illuminating the genetic architecture of complex traits (13–16). The P3GMap reveals recurrent pathways ranging from immunity, inflammation, and DNA repair to growth factor signaling, linked to phenotypic change, echoing prior evidence from long-lived mammals (27,62–65) and other comparative frameworks (24–27,61). Our case studies connect genomic signatures to insectivory (31–33,107–108), immunity and WBC variation (34–35,126–129), and lifespan (24–27,134–136), illustrating the value of targeted *C*ontrasts for dissecting lineage-specific adaptations. As an open and scalable platform, the PGA provides a foundation for extending genome –phenome inference beyond protein-coding variation, thereby strengthening the link between comparative genomics and human biomedical research.

## Acknowledgements

We thank the members of the Human Genetics and Comparative Genomics laboratories at the Institute of Evolutionary Biology (IBE-UPF) for insightful discussions on phylogenetic comparative methods, trait curation, and the functional interpretation of our results. We are grateful to the researchers who compiled the primate phenotypic datasets used in this study, including the ecological trait database of Galán-Acedo et al. (31), and to the consortium that generated the 233-primate-genome resource underpinning our comparative analyses (28). We also thank the developers of RERconverge (the Clark and Chikina groups) for creating a methodological framework that was central to this work. This research was supported by the UPF Scientific Computing Core Facility. The authors further acknowledge the National Centre for Genomic Analysis (CNAG) and the Barcelona Supercomputing Center (BSC) for providing access to high-performance computing resources.

