## Supplementary Information for "A phylogenetic protein-coding genome-phenome map of complex traits across 224 primate species"

**The PDF file includes:**

Materials and Methods
Figs. S1 to S50
Tables S1 to S7

**Other Supplementary Materials for this manuscript include the following:**

Data S1 to S3

Materials and Methods 5
1. Phenomic Data and Trait Modeling 5
1.1 Retrieval and classification of primate traits 5
1.2 Comparable traits clusters definition and correlations 5
1.3 Preprocessing of phenomic dataset for phylogenetic comparative analyses 8
2. Genomic Data Processing 9
2.1 Estimation of Reference Orthology Clusters 9
2.2 Protein-coding gene alignments generation 9
3. Integrated Phylogenetic Analyses 10
3.1 Phylogenetic comparative methods (PCMs) application 10
3.1.1 RERconverge 10
3.1.1.1 Permulation statistics 11
3.1.2 Convergent amino acid substitution (CAAS) 12
3.1.2.1 Accounting for species with informative trait values 12
3.1.2.2.Search of within-family contrasts and Bottom refining 12
3.1.2.3 Accounting for pairs of informative phylogenetic clades 12
3.1.2.4 Study of two scenarios of AA change 13
3.1.2.5 Primate consistency test 13
3.1.2.6 Equivalent non-primate mammalian positions retrieval for CAAS positions 14
3.1.2.7 Cross-mammalian consistency test 14
3.1.2.8 Redundancy study of CAAS species selection 15
3.1.2.9 What CAAS is and what CAAS is not 15
3.2. Natural selection detection 16
4. Functional Enrichment Analyses 16
4.1 Overrepresentation analysis (ORA) 16
4.2 Trait domains gene set overlap 17
5. Human Genomic Mapping and Annotation of CAAS Positions 18
5.1 Human genomic mutations and conservation scores retrieval for CAAS positions 18
5.2 Human alternative allele frequency information retrieval (AF) 18
6. Functional Genomic and GWAS-Based Analyses of CAAS Genes 18
6.1 Gene set analysis in human GWAS with MAGMA 18
6.2 Gene set heritability enrichment in human GWAS with S-LDSC 19
6.3 Over-representation of high-impact mutation genes 20
Supplementary Figures 21
Supplementary Tables 71
Data S1 (separate file). 79
Data S2 (separate file). 79
Data S3 (separate file). 79

Technical Appendix 80
Software versions, command-line parameters, and reproducibility resources 80
TA1. Software used for genomic and phylogenetic analyses 80
TA2. Orthology detection 81
TA3. Multiple sequence alignment 81
Protein alignments 81
Codon-aware alignment of CDS 81
TA4. Alignment filtering 82
TA5. Phylogenetic tree processing 82
TA6. Detection of convergent amino acid substitutions (CAAS). 83
TA7. Figure generation 83
TA8. Reproducibility and data availability 83

### Materials and Methods

#### 1. Phenomic Data and Trait Modeling

##### 1.1 Retrieval and classification of primate traits

Trait measurements for primate species were compiled from diverse studies with varying sampling depths and strategies, harmonized meta-analyses, and several databases (Data S1). In all cases, we focused on large-scale studies involving a high number of species. We did not combine measures from different sources unless they were already integrated by the curation efforts of the included sources. Manual inspection identified several outliers that were evident typographical errors in the original sources; these were corrected when possible or removed. To facilitate traceability, we retained the trait names from the original sources. This effort resulted in an initial dataset of 835 redundant traits, which contained different implementations of the same traits from various sources. These redundant traits were initially included in the trait comparisons to test their consistency and to compare their species coverage.

From a biological perspective, we grouped traits into 15 categories (secondary domains) nested within five main trait domains—morphology, life history, physiology, behavior, and ecology—following a hierarchical approach from these primary domains to more specific levels of biological complexity. The definition of a biological trait, its selection for genomic architecture analysis, and its classification into well-delimited groups remain debatable (1–4). Using these domains, we grouped as many traits as possible into clearly defined sets representing the major areas of biological organization.

#### 1.2 Comparable traits clusters definition and correlations

| Primary Domain | Secondary Domain | Number of Traits |
|---|---|---|
| Behavior | Diet | 4 |
| | Habitat use | 11 |
| | Social organization | 19 |
| | **Total** | **34** |
| Ecology | Climate | 6 |
| | Conservation | 2 |
| | **Total** | **8** |
| Life history | Longevity | 12 |
| | Reproduction | 28 |
| | **Total** | **40** |
| Morphology | Body mass | 39 |
| | Brain | 39 |
| | Coloration / ornamentation | 2 |
| | Organs | 12 |
| | Sexual selection | 20 |
| | Vestibular and visual systems | 2 |
| | **Total** | **114** |
| Physiology | Biometabolites | 60 |
| | Energetics | 4 |
| | Vestibular and visual systems | 3 |
| | **Total** | **67** |

**Table S1. Summary of phenotypic trait classification across the biological domains.**

From our initial dataset of 835 traits obtained from public sources (Data S1), we excluded secondary domains of traits that contributed limited phylogenetic information and could add noise to the downstream analyses. Specifically:

*Morphometric quantitative traits* (N = 220): Most of these (185 traits) were primarily measured in the Cercopithecidae family, with data for at least 20 species from this family. Because these traits are largely restricted to a single family, they provide limited information on broader phylogenetic patterns.

*Parasite–host qualitative traits* (N = 352): Of these, 319 traits had phenotypic data for fewer than five species, making them too sparse to be informative across the dataset.

These groups were excluded to reduce dimensionality, minimize noise, and focus on traits that offered a meaningful phylogenetic signal across multiple taxa. After applying these filters, we retained a final dataset of 263 traits. A summary of the filtering steps applied to the initial phenomic dataset is presented in Table S2. Traits were organized into five primary biological domains: morphology, life history, physiology, behavior, and ecology, and further subdivided into secondary domains representing more specific biological processes (Table S1). A complete list of traits and associated metadata is provided in Data S1.

To characterize the heterogeneity in species coverage across traits, we performed Multiple Correspondence Analysis (MCA) using factorMineR on a binary matrix representing the presence/absence of trait measurements for each non-human primate species. We then applied Hierarchical Clustering on Principal Components (HCPC) using the first ten MCA components (which explained 82.22% of the variability in species coverage) to identify groups of traits with similar distributions of covered species. To determine the optimal number of clusters, we performed K-means clustering across a range of 2–90 clusters and selected the best solution using the elbow method to identify the optimal inertia (Fig. S1). This process resulted in the classification of traits into 13 clusters with comparable species coverage (Fig. 1A, Fig. S2, Fig. S3, Data S1).

This trait classification by species coverage accounts for the varying proportions of missing data across species and traits, indicating which species can be reliably compared within each group. These species–trait groups enabled us to study the main contributors to the principal components. To this end, we excluded species with less than 90% trait overlap within a cluster, retaining only the most comparable ones. We then applied Factor Analysis of Mixed Data (FAMD) and examined the phylogenetic evolutionary correlations between traits within each cluster and traits from external clusters sharing at least 90% of the same species. FAMD integrates quantitative and qualitative variables and reduces dimensionality while preserving the key patterns of covariation. Traits were projected onto the first two dimensions (Dim1 and Dim2) and represented as arrows, with the length and color reflecting their relative contribution to the dimension (percentage of variance explained). This approach highlights the main traits driving variation and allows the visualization of covariation patterns among secondary domains.

Clusters VII–XIII covered life history (27.6%), morphological (9.7%), and ecological data (100%) with a wide phylogenetic distribution. Cluster VIII (n = 46 species / 15 traits) contained brain mass, brain size in females, neonatal mass, and body mass measurements, with more than five traits contributing strongly to the first component of the cluster (see Supplementary Figs. S4–S6). Other clusters with broad and diverse trait representation, such as Cluster I (n = 23 species) and Cluster IV (n = 30 species) (see Supplementary Figs. S13–S19), provide sources of deeper phenotypic knowledge on the strepsirrhine clade, coming mainly from captive individuals monitored in conservation and protection programs at centers such as the Duke Lemur Center (DLC) and the Henry Doorly Zoo (HDZ).

Finally, to detect similarities in the evolution of different traits and the consistency between highly related or redundant traits, we obtained correlation matrices corrected for the phylogenetic non-independence of the species represented in each cluster (Supplementary Figs. S5, S7, S9, S11, S13, S15). For this, we used the phylogenetic covariance matrices implemented in R for continuous traits (5):

```
obj <- phyl.vcv(X, vcv(tree), 1)
corr <- cov2cor(obj$R)
```

In contrast, for clusters containing categorical variables, we incorporated the concept of liabilities (underlying continuous traits that describe an existing discrete character) proposed in the threshold model (6) and implemented it using a Bayesian threshold framework, allowing the estimation of correlations between pairs of variables (two qualitative or one qualitative and one quantitative):

```
fit <- threshBayes(tree, X, types = c(type1, type2))
corr <- mean(fit$par[, “r”])
```

| Filtering step | Description | Traits removed | Traits remaining |
|---|---|---|---|
| Initial dataset | Traits compiled from public sources (Data S1) | 0 | 835 |
| Removal of morphometric traits | Quantitative morphometric traits largely restricted to Cercopithecidae species | 220 | 615 |
| Removal of parasite–host traits | Qualitative parasite–host traits with sparse species coverage (<5 species) | 352 | 263 |
| Final dataset | Traits retained for downstream analyses | 0 | 263 |

**Table S2. Filtering steps applied to the initial phenomic dataset.**
From an initial set of 835 traits compiled from public sources (Data S1), traits that provided limited phylogenetic information were excluded. Morphometric traits largely restricted to Cercopithecidae species and parasite–host traits with sparse species coverage were excluded from the analysis. The final dataset retained 263 traits for downstream comparative analyses.

#### 1.3 Preprocessing of phenomic dataset for phylogenetic comparative analyses

Because our dataset integrated multiple existing phenotypic databases of related primate traits, we removed redundancy when the same biological trait appeared in multiple databases. In such cases, we retained the version with the largest number of sequenced species and primate families ( Table S3).

From a data-encoding perspective, we classified the traits as quantitative continuous or qualitative (binary, multinomial, and discrete <10 states). Discrete traits were binarized by coding the minimal number of higher states of the trait representing at least two primate species as 1 and the remaining lower states were coded as 0. Dummy variables were created for each category of multinomial traits. Species with the presence of the corresponding category were coded as 1 (*e.g.*, nocturnal species from the diel activity trait when studying nocturnality), and those without the category were coded as 0 (diurnal and cathemeral species from the diel activity trait when studying nocturnality). Quantitative traits measured in fewer than 10 species and qualitative (binarized) categories covering fewer than two families in the sequenced primates were excluded from further analyses, as these traits provided insufficient data to capture meaningful patterns and could potentially bias the results. After applying these filters, we retained 200 traits as the final set of curated traits used in the phylogenetic genome-phenome association analyses.

Allometric relationships between body size and many biological traits strongly influence the evolutionary history of morphological, life-history, and physiological phenotypes (7–13). Larger body sizes in the Hominidae and Cercopithecidae families could confound the identification of species with extreme values for other traits. This influence is evident in the high correlations between body mass and many other traits in our dataset (see Supplementary Figs. S5, S7, S9, S11, S13, S15, and S21). To control for this effect, for traits scaling with body mass, we performed phylogenetic generalized least-squares regression (PGLS) (14) using the caper package (v1.0.1) to obtain residuals that simultaneously accounted for body mass covariation and phylogenetic effects based on the relevant body or body part size metric for each dataset. For all brain region size traits, corrections were applied directly to their respective brain size traits, thereby accounting for both brain and body size scaling in a single step.

### 2. Genomic Data Processing

#### 2.1 Estimation of Reference Orthology Clusters

Using protein-coding gene annotation equivalences of 20,417 human gene models from Ensembl v98, we focused on the predicted human-based peptides from nonhuman primates in a dataset comprising 59 high-quality reference primate genomes (15). For each human gene model, we identified orthologous pairs of protein-coding genes by conducting the best bidirectional hit searches (16) across the initial set of primate species. An iterative approach was used for all human gene models to generate orthology clusters for each reference primate species, in which reciprocal best hits were identified between that species and every other reference species.

After assigning all peptides to one or more species-centered orthology clusters, we retained the cluster whose reference recovered the largest number of orthologous relationships for each human gene, prioritizing *Homo sapiens* as the main reference, based on annotation quality. We validated this clustering method using reciprocal best chains from six UCSC primate genomes (*Chlorocebus sabaeus*, *Otolemur garnettii*, *Pan troglodytes*, *Pan paniscus*, *Papio anubis*, and *Pongo abelii)*. Most genes suitable for liftover were successfully validated indicating a very high concordance and confirming the reliability of the clustering approach.

From the 18,858 orthologous clusters obtained, we retained 16,177 clusters with genes in ≥40 of the 59 reference species to ensure the consistency of orthologous relationships across the phylogeny of primates. Multiple sequence alignments of the final peptide clusters were generated (17) and subsequently filtered (18) to remove poorly aligned regions and sequences with low alignment coverages. Detailed information on the software versions and command-line parameters used for orthology detection and alignment generation is provided in the Technical Appendix.

#### 2.2 Protein-coding gene alignments generation

For the resequenced individuals from the largest genomic catalog of world primates (19), we selected one representative individual per species, choosing the individual whose read mapping to its closest reference genome recovered the largest number of genes within the first decile of the callability overlap distribution (19). Reconstructed coding DNA sequences (CDS) from these selected individuals were assigned to orthology clusters established in the previous step.

Reconstructed coding sequences depend on base-calling uncertainties that can lead to alterations in coding sequences, potentially resulting in incorrect amino acid translations or codon frameshifts. To minimize the impact of these uncertainties, coding sequence alignments were generated using a codon-aware alignment strategy that preserved the reading frame and enabled the identification of frameshifts and premature stop codons (20–21).

Reference coding sequences were first aligned and subsequently enriched with resequenced primate sequences to generate multiple sequence alignments containing all available species in each orthology cluster. Because frameshifts are more likely to result from sequencing or gene model definition errors than from true biological events, they were masked before downstream analyses.

To further ensure the reliability of the alignments, we applied two filtering steps to the data. First, we performed a chi-square test for each gene alignment to detect and remove the least reliable sequences, which were characterized by a significant number of unresolved ambiguous codons relative to well-aligned codon positions. Second, poorly aligned codon positions were filtered (22) by removing positions containing a high proportion of gaps across the species.

This procedure led to the recovery of a final set of 16,133 multiple sequence alignments ( Data S1), with 11,885 (>70%) containing alignments for more than 200 resequenced species (Fig. S16), which constituted the final genomic dataset used to build the genetic predictor variables. The detailed command-line parameters and software versions used for sequence alignment and filtering are reported in the Technical Appendix.

### 3. Integrated Phylogenetic Analyses

#### 3.1 Phylogenetic comparative methods (PCMs) application

##### 3.1.1 RERconverge

For the filtered set of primate coding sequence alignments, we obtained the corresponding gene trees based on the multiple sequence alignments generated in the previous steps. To do so, we fixed the tree topology to the previously obtained UCE-based primate species tree topology (19). For each multiple sequence alignment, this topology was pruned (23) so that only the species present in the corresponding gene alignment were included in the analysis.

Branch length estimates for the gene trees were obtained using a codon-based likelihood estimation approach (24). The relative evolutionary rates (RERs) for each alignment were estimated using the RERconverge package (25) in R. A square-root transformation was applied to the set of RERs to reduce heteroscedasticity, following the benchmarking recommendations of the software. Variance across the tree was accounted for by scaling individual branches, and weighted regression was performed to correct for the relationship between the mean and variance, as suggested by the RERconverge guidelines.

For quantitative non-allometric variables, RER–trait correlations were estimated using log10 transformations for traits rejecting the Shapiro–Wilk test null hypothesis of normality (p-value $< 0.05$), whereas raw trait values were used when the null hypothesis was not rejected (p-value $> 0.05$). Allometric-corrected traits were used directly after phylogenetic generalized least-squares (PGLS) correction with the corresponding body mass traits from their respective sources when available.

For qualitative binarized traits, gene–trait correlations were estimated by providing information for both the foreground branches for the trait of interest and the sister species in the master tree, as required by the software for binary correlation analysis. In both quantitative and qualitative cases, before the correlation step, we converted the phenotype vector into phylogenetic paths comparable to those in the RER matrix using functions from the RERconverge package.

Because multiple genes were tested per trait, we used the multiple-gene-corrected p-value based on the Benjamini–Hochberg false discovery rate correction (Data S2). The software versions used for these analyses are listed in the Technical Appendix.

###### 3.1.1.1 Permulation statistics

In addition, to obtain empirical p-values, we generated null phenotypes for our traits by permulating the traits following a Brownian Motion (BM) model of evolution. An empirical p-value metric ($p_{\text{Perm}}$) was estimated by measuring the proportion of permulated gene-trait correlations between the RERs of the corresponding gene and these null phenotypes, which were as extreme or more extreme than the parametric correlations estimated in the previous step.

For quantitative traits, 10 batches of 100 permulation runs (for a total of 1,000 permulations per trait) were performed for both non-allometric and allometric-corrected traits. For each trait, we obtained estimates for each gene-trait *Pearson* correlation (*Rho* statistic). For qualitative binarized traits, 10 batches of 100 permulation runs (again totaling 1,000 permulations per trait) were performed using the Complete Case (CC) method, producing one set of permulated phenotypes for all genes using the *master tree* obtained after parsing the corresponding gene trees from the previous steps. Because computational efficiency was lower for binarized traits than for quantitative traits in our dataset, only binarized categories for which each batch of 100 permulations required <=24h of computational time were included in our analyses. Different batches of permulations were combined using the combinePermData function.

##### 3.1.2 Convergent amino acid substitution (CAAS)

###### 3.1.2.1 Accounting for species with informative trait values

To identify the most relevant shifts in each primate trait, we searched for primate families containing species with *Top* extreme and/or *Bottom* extreme trait values.

First, the *Top* candidate species were selected. For quantitative traits, we selected the most extreme families with member species above two median absolute deviations (MAD) from both the median of the family and the median of the global trait distribution. Second, primate families with >2 MAD from the median of the trait not included in the first step were added to the *Top*. For qualitative binarized traits, we selected all species that displayed the corresponding category.

Second, we selected the *Bottom* candidate species. For quantitative traits, we applied the same criteria as the *Top* configuration, but with species below 2 MAD from both the median of the family and global trait distribution. In the second step, primate families with MAD values <2 that were not included in the first step were added to the *Bottom*. For qualitative binarized traits, we selected all species that displayed phenotypes other than those of the corresponding category.

###### 3.1.2.2.Search of within-family contrasts and Bottom refining

We identified the family contrasts in our dataset. These contrasts reflect strong trait shifts between closely related species in the phylogeny, thus helping to reduce phylogenetic noise in our criteria.

For quantitative traits, we searched for species with values below or equal to the median of the global trait distribution whose family was also present in the *Top* group of the trait as *Bottom* candidates. For qualitative binarized traits, we searched for species lacking the corresponding category whose family was present in the *Top* group of the trait as *Bottom* candidates.

###### 3.1.2.3 Accounting for pairs of informative phylogenetic clades

For quantitative traits, when ≥ 2 *family contrasts* were available, we selected those from the two families with the *Top* species displaying the highest trait values. When only one *family contrast* was available, we selected it and the *Top* species from the family with the next highest trait values. When no *family contrasts* were available for the trait, but ≥ 2 *Top* families met the extreme selection criteria, we selected the two *Top* families whose species had the maximal patristic distance between them to avoid phylogenetic contiguity in this selection. In the selection of Top species without contrasts, we gathered the remaining *Bottom* families for each case, meeting two criteria: a) species with trait values below or equal to the median of the trait distribution, and b) families with the smallest median proportion of alignments lacking their species (see Fig. S17 - Fig. S18).

For qualitative binarized traits, when ≥ 2 *family contrasts* were available for the trait, we selected the two most informative families, defined as those with the lowest presence/absence ratio of the respective category in the family. When only one *family contrast* was available, we selected the second *Top* family with the lowest proportion of present species. In this case, the added *Bottom* family corresponded to the one with the smallest median proportion of missing alignments (Fig. S19).

The selection of informative primate species for our trait measurements yielded 77 quantitative traits (56 with at least one primate family having species in both the *Top* and *Bottom* sets, hereafter referred to as *family contrast*, and 21 with *Top* and *Bottom* species from different families) and 57 qualitative binarized categories with at least one primate *family contrast*.

3.1.2.4 Study of two scenarios of AA change

Consistent with previous implementations of the CAAS algorithm in genome-phenome studies (26–27), prior to the development of CAAStools (v1.0) (28), we considered two CAAS scenarios for amino acid positions across groups of species. In Scenario 1, all species within the Top group exhibited identical amino acids, whereas those in the Bottom group shared a different amino acid. In Scenario 2, all species in the Top group shared the same amino acid, whereas species in the Bottom group displayed various amino acids that differed from that in the Top group.

To assess the significance of these two scenarios at every position where they were detected, we conducted 1,000 trait permulations and calculated an empirical p-value as the ratio of times the position reflected these scenarios for the permulated traits. We excluded seven genes from the analyses because of the extensive computational time required, which exceeded 24 h to run 1,000 permulations for all traits.

We applied strict criteria to search for CAAS, considering only positions where all species in the *Top* and *Bottom* groups were present in the protein-coding alignment and no gaps were present for any of the species included, using the options --max_fg_miss, --max_bg_miss, --max_bg_gaps, and --max_bg_gaps. Positions with missing amino acids in either the Top or Bottom groups were excluded from the analysis.

3.1.2.5 Primate consistency test

We tested the significance of our amino acid-trait links in the entire primate phylogeny for each discovered position. We only considered the same direction of the CAAS pattern (*one-tailed* $p_{PGLS}$) using primate species with phenotypic values not selected for our *Top* and *Bottom* groups in the previous phase. As noted in previous studies, many comparisons in which few species were present in one of the two groups may be statistically underpowered. Therefore, for the search of top-ranked AA-trait links and estimation of percentages, we only considered positions where both *Top* and *Bottom* amino acids were present in >5 internal primate species.

For quantitative traits, we used phylogenetic generalized least squares (PGLS) with the R package caper (v.1.0.1), fitting Pagel's λ, and applying Randomization of Residuals in a Permutation Procedure (RRPP) (29) under a Brownian-motion evolutionary model for quantitative measurements. For PGLS, we obtained at least one amino acid position with FDR-trait <0.05 for 16 measurements with inter-family contrasting phenotypes, six measurements with one family with contrasting phenotypes, and 11 measurements with two or more families with contrasting phenotypes, while for RRPP, we obtained associated gene sets for Minimum Female Parent Age At Conception, Uric Acid Levels and Cholesterol Levels measurements with FDR-trait <0.05.

For qualitative measurements, we performed a phylogenetic logistic regression (phyloglm) (30), only obtaining putatively associated amino acid positions for trichromatic color vision with an FDR trait < 0.05.

3.1.2.6 Equivalent non-primate mammalian positions retrieval for CAAS positions

TOGA alignments were retrieved for 36,509 orthologous transcripts from 427 mammalian species (31). Filtering was performed at the codon sequence alignment level. We removed poorly aligned species (<50% aligned residues in positions with >50% coverage) and codon columns with >50% gaps using trimAL (v1.2) (18) and BMGE (v1.1) (22). For each transcript alignment, species with >1 orthology were discarded from the analyses, and aligned sets of 1-to-1 mammalian orthologs were retrieved. For each codon alignment, the corresponding amino acid alignments were produced using BMGE, considering codon structure.

We obtained mammalian alignment positions corresponding to the primate alignment positions recovered by CAAS based on the reference genomic coordinates in the human CDS that are present in both sets of alignments (see Figs. S27–S28).

3.1.2.7 Cross-mammalian consistency test

We obtained information on 11 equivalent traits with CAAS detected in our primate phenomic dataset for non-primate mammalian species from the PanTHERIA resource (32), including age of females at first reproduction, body mass, gestation period length, interbirth interval length, lactation period length, litter size, maximum lifespan, mean-niche precipitation, mean group size, mid-range latitude, and nocturnal activity. Trait data processing before genome–phenome analysis was performed consistently with that used for primate-equivalent traits in this study. We focused on 191 species in the

Zoonomia TOGA codon alignments (31,33) with trait information, from which we excluded the 36 primate species already included in our main dataset.

We obtained information from our Top and Bottom sets of amino acids discovered by CAAS and searched for positions where these amino acids were present in mammalian protein-coding gene alignments. We performed two different analyses, following the same approach as our primate consistency test for CAAS discovery:
(a) We ran PGLS (fitting the λ model) between the predictor variable (species with *Top* and *Bottom* amino acids) and the response variable (quantitative trait), examining the association in the direction of the *Top* amino acid.
(b) We applied the Randomization of Residuals in a Permutation Procedure (RRPP), assuming the Brownian Motion model, to the quantitative trait set to test the significance of links between our Top and Bottom amino acid sets across the entire mammalian phylogeny, in the same direction as the CAAS pattern.

For qualitative traits, we performed phylogenetic logistic regression (phyloglm) using the phylolm package in R. We used the multiple-gene-corrected p-value from RERconverge based on the Benjamini–Hochberg correction (FDR) (Data S2). The final set of phenotypes included in the genome–phenome analyses is summarized in Data S1. For the search of top-ranked amino acid–trait links and the estimation of percentages, we only considered positions where both Top and Bottom amino acids were present in >5 external mammalian species, obtained a p-value of association, and retained traits with at least ten available positions. The significant gene–phenotype associations identified in these analyses are presented in Table S5.

###### 3.1.2.8 Redundancy study of CAAS species selection

To assess the degree of redundancy in our species selection for CAAS analysis, for each pair of traits with available CAAS selection for the Top and Bottom species, we computed a corresponding metric of the Jaccard distance (34) (1-Jaccard similarity) for the overlap of species between the Top groups of the pair of traits and the Bottom groups of the same pair of traits (see Fig. S19). A value of 1 corresponds to the maximum possible distance between the categories for each pair of traits, indicating that there is no overlap of species, whereas 0 reflects the same configuration of species between the groups of both traits. A summary of the overlapping gene sets identified across the analytical approaches is provided in Table S6.

###### 3.1.2.9 What CAAS is and what CAAS is not

CAAS analysis identifies amino acid substitutions whose distribution across species is statistically associated with phenotypic differences in the phylogenetic context. In this framework, CAAS positions are best understood as candidate protein-altering variants linked to trait evolution, rather than as linkage-based markers typically identified in genome-wide association studies. Because the associations are evaluated across lineages separated by millions of years of evolution, the signal is unlikely to reflect a long-range linkage disequilibrium persisting from nearby causal variants. However, the amino acid substitution itself is a plausible candidate for functional relevance.

In addition, we must bear in mind that CAAS associations do not by themselves demonstrate direct or simple causation. A significant CAAS may reflect a direct effect of the amino acid change on the focal phenotype, but it may also capture broader pathway-level relationships or indirect causal routes involving development, physiology, immunity, ecology, and life-history evolution. Therefore, CAAS results should be interpreted as candidate evolutionary links between coding variations and phenotypes, the mechanistic basis of which may vary from trait to trait.

This distinction is important for interpretating of individual examples. For instance, the evolutionary association reported in the main text between CAAS in **MEFV** and lactation period length does not necessarily imply that **MEFV** directly determines lactation duration. A more plausible interpretation is that immune and inflammasome-related biology may contribute to the broader physiological context in which lactation evolved. Similarly, an evolutionary association between **MPDZ** and gestation period length does not require a direct effect of this gene on gestational timing, but may instead reflect the involvement of epithelial barriers or neurodevelopmental processes that co-evolve with reproductive timing. In this sense, CAAS signals should be viewed as biologically informative candidates that motivate downstream functional and experimental validation, not as definitive proof of one-gene, one-trait causality.

#### 3.2. Natural selection detection

Directional selection analyses were performed on the protein-coding gene datasets for alternative species selections in our study cases (Insectivorous Diet, Maximum Lifespan, and White Blood Cell Count). We applied the FUBAR Approach to Directional Evolution (FADE) method (35) implemented in HyPhy (36), which detects substitution biases toward specific amino acids relative to the background evolutionary process.

This approach searches for sites of directional selection in protein-coding gene alignments and has recently been applied in comparative analyses, such as studies on the directional selection of social spiders. In our context, FADE provides a complementary framework to the CAAS approach by identifying substitution patterns biased toward specific amino acids in specific evolutionary lineages (37).

For each case study, we labeled the phylogeny of the analyzed species (38) to define the foreground and background branches. Foreground branches corresponded to species selected based on their trait values according to the CAAS framework, whereas the remaining species in the alignment were used as the background.

For the Insectivorous Diet, we selected branches leading to species from Daubentoniidae, Lorisidae, and

Tarsiidae. For Maximum Lifespan, branches leading to *Cebus*, *Sapajus*, *Hylobates*, and *Homo* were used as foreground genera. For White Blood Cell Count, we selected species from three primate families with contrasting phenotypes: Hominidae, Lorisidae, and Callitrichidae. The remaining species in each alignment were used as background sets.

Detailed information on the software versions and execution parameters used for these analyses is provided in the Technical Appendix.

### 4. Functional Enrichment Analyses

#### 4.1 Overrepresentation analysis (ORA)

We performed overrepresentation analyses (39) to identify enriched functional categories associated with the candidate gene sets obtained from genome–phenome analyses. Functional enrichment was evaluated using Gene Ontology categories (Biological Process, Cellular Component, and Molecular Function), human diseases from the DisGeNET database, human phenotypes from the Human Phenotype Ontology, and biological pathways from KEGG and Panther.

For each analysis, enriched categories were identified using a hypergeometric framework and were considered significant when passing multiple testing corrections with a false discovery rate threshold of FDR < 0.05. Category size filters were applied to exclude very small or excessively large functional categories from the analyses. Additional enriched Gene Ontology terms associated with body mass-related traits are listed in Table S7.

First, we analyzed gene lists corresponding to a subset of genes showing nominally significant permulation p-values (p-perm < 0.05) for both the RER and CAAS methodologies, providing a broad exploratory overview and enabling comparison between the two analytical approaches. The results obtained for each candidate gene set were evaluated relative to a background set of genes corresponding to the list of genes analyzed for the respective traits and methods.

Second, for CAAS analyses, we explored gene lists corresponding to genes containing internally significant CAAS positions identified in phylogenetic tests for each trait. The results obtained for each candidate gene set were evaluated relative to the background list of genes containing CAAS positions detected for the corresponding traits.

The complete set of enriched functional categories obtained from the overrepresentation analyses is shown in Table S7. The software versions used for these analyses are listed in the Technical Appendix.

#### 4.2 Trait domains gene set overlap

For each of the two PCM methodologies applied, we performed a hypergeometric test of the gene list overlap between the gene sets recovered for each pair of traits with a significant permulation analysis (p-perm < 0.05). We used the phyper package in R (40), implementing the hypergeometric distribution with our universe of protein-coding gene alignments for each pair of traits and method. The resulting p-values were corrected for multiple tests (FDR < 0.05) for each trait.

We used the gene lists corresponding to the subset of total genes showing a significant permulation p-value (p-perm <0.05) for both the RERs and CAAS methodologies to obtain both a comprehensive exploratory view and consistency between methods. Additionally, for CAAS analyses, we used gene lists corresponding to a subset of total genes showing primate-consistent significant p-values ($p_{PGLS} < 0.05$) per trait.

### 5. Human Genomic Mapping and Annotation of CAAS Positions

#### 5.1 Human genomic mutations and conservation scores retrieval for CAAS positions

We obtained a full set of genomic mutations underlying the CAAS positions detected at the amino acid level in primate protein-coding gene alignments. The corresponding positions were retrieved using the variant annotation software Transvar (v2.3.4) (41) and the panno function to search for specific protein variant annotations. For SNP mutations in our protein variants that mapped to multiple transcripts, we filtered the results to retain the genomic mutation corresponding to the transcripts used in the respective human gene models. After obtaining the set of genomic positions, we determined their corresponding conservation scores using the Zoonomia phyloP dataset (31,33) for mammalian alignments based on the human genome assembly version hg38. We then divided our CAAS results into three sets 1 of positions: all CAAS positions (including Scenarios 1 and 2), CAAS positions in Scenario 1, and CAAS positions in Scenario 2. As a background reference for CDS variability in terms of PhyloP scores, we sampled a total of 1 × 107 human CDS positions and compared them with our three CAAS-derived sets (see Fig. S29). This analysis allowed us to evaluate the evolutionary constraint at CAAS positions, showing that these sites generally occur in a more genetically variable background than the majority of the exome, which tends to be highly conserved.

#### 5.2 Human alternative allele frequency information retrieval (AF)

To determine whether the CAAS variants detected in primate alignments are fixed or variable in human populations, we used the gnomAD (v.3.0) (42) database to retrieve the alternative allele frequency (AF) for biallelic SNP positions in the hg38 human genome, based on genomic coordinates obtained with TransVar and corresponding to the mutation linked to the CAAS position detected (see Fig. S30). By retrieving this information, we can assess whether the CAAS variants detected in primate alignments are fixed or variable in the human population, respectively. This provides an estimate of the number of fixed versus variable positions in humans, offering a context for comparing the genetic background associated with primate genes to that of humans.

### 6. Functional Genomic and GWAS-Based Analyses of CAAS Genes

#### 6.1 Gene set analysis in human GWAS with MAGMA

Per-trait gene sets with significant results from permulations of CAAS and RERconverge analyses were tested for functional enrichment in human GWAS for the corresponding human traits using basic competitive analysis in MAGMA (43) against the background gene sets of all per-trait genes analyzed. First, every gene in the definition file from MAGMA (NCBI build 37 genome annotation, with a total of 19427) considers a range of 5 kb upstream and downstream. The 1000G data were used as the base population for LD and MAF correction using MAGMA (g1000_eur for EUR GWAS and g1000_eas for EAS GWAS). Using gene-level analysis, the P-values for each gene were estimated for each trait.

For the basic competitive analysis, the gene sets for each trait were used in the model using the *—set-annot flag*, and the total number of analyzed genes (background) for the same trait was used with the *-include flag* to limit the analysis to genes within or outside the permutation list but within the background list. The resulting beta is the beta difference between the gene set and the complementary gene set in the background list. The results were meta-analyzed using a random-effects meta-analysis for the 19 (CAAS) and 26 (RERs) GWAS/traits analyzed.

#### 6.2 Gene set heritability enrichment in human GWAS with S-LDSC

The relevance of the per-trait hit gene 5% and per-trait associated pathways from both RERs and CAAS was evaluated in terms of their impact on the trait heritability of the corresponding human equivalent traits (Data S1). We estimated relative heritability enrichment with S-LDSC (44) using GWAS summary statistics for the corresponding human heritable traits (SNPh2 > 5 % and SNP z >7), homologous to the primate traits analyzed. Enrichment in the partition corresponding to (A) trait-RERs/CAAS significant genes or (B) trait-ORA pathways was estimated and compared to the partition comprising the genes in the corresponding analysis background, as described below.

A) A) Genes strategy: The partition was defined as a trait-RER/CAAS gene body + 5 kb. In every case, the background was the gene body + 5 kb flanking all genes analyzed per trait and the comparative methodology used. The results were meta-analyzed using a fixed-effects model.

B) B) Pathways strategy: Partition was defined as the gene body + 5 kb flanking the genes in the set of all genes annotated to any of the trait-RERs/CAAS over-represented categories in the ORA. In every case, the background was the gene body + 5 kb flanking the genes annotated to any category in the annotation databases (GOslim BP, GOslim MF, GOslim CC, HPO, and DisGeNET). The per-trait results were meta-analyzed using a fixed-effects model.

Analyses were limited to human traits with a GWAS SNPh2 z score > 7 and annotations (gene sets) covering at least 0.5% of the heritability SNPs, following the authors' recommendations. To create annotations, the gene coordinates corresponding to the CDS used in the phylogenetic analyses (hg38, Ensemble v98) were obtained using the biomaRt R package, and the positions were converted to hg19 coordinates using the UCSC liftOver tool (45).

Because lower conservation (median phyloP scores) in the CAAS gene sets compared to the background category could bias the relative enrichment towards the null hypothesis, we performed additional tests: 1) analysis removing conserved positions (phyloP scores > 0); 2) quantitative analysis estimating enrichment per quintile of phyloP scores; and 3) analysis and meta-analysis restricted to traits showing no statistically significant differences in *phyloP* between the RER/CAAS result and background gene sets.

#### 6.3 Over-representation of high-impact mutation genes

Considering the possibility that common variant heritability is not a good representation of gene involvement in a trait, we evaluated whether the RER/CAAS gene sets also have a high impact on human phenotypes and diseases. Specifically, we tested whether the RERs/CAAS genes were overrepresented among genes annotated as harboring high-impact mutations in human phenotype/disease databases (Fig. S49-Fig. S50), using the autosomal inheritance OMIM database (HPO high frequency, >30 %) (46).

#### Supplementary Figures

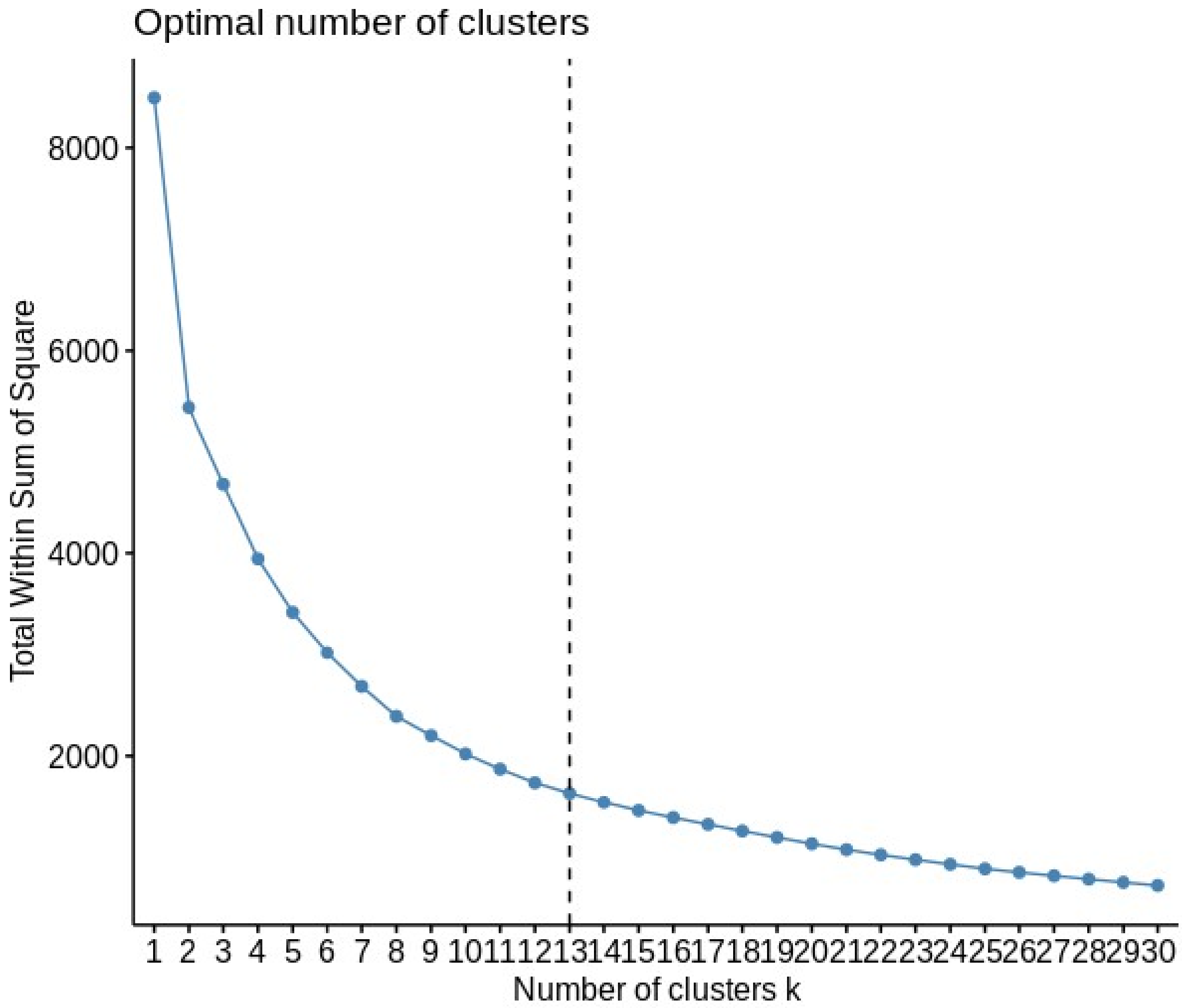


**Fig. S1.**

Optimal number of clusters (k=13) for our phenomic dataset using the elbow method and k.max=90

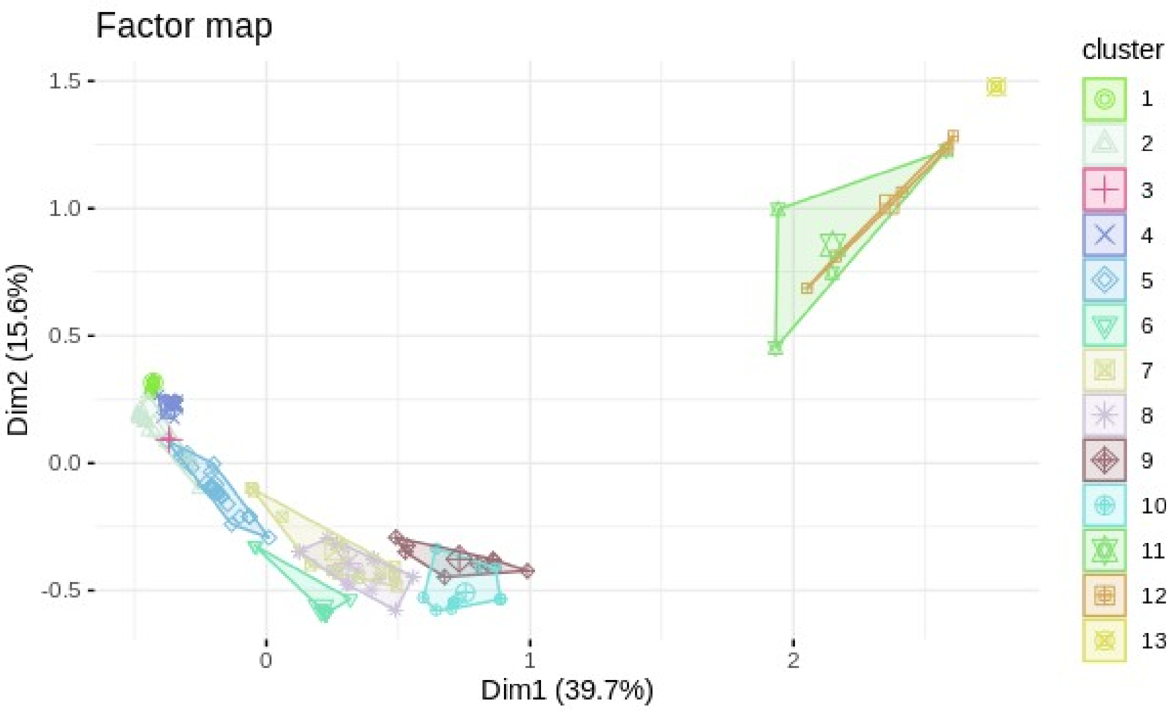


#### Fig. S2.

Representation of 2 first dimensions of the factor map for our set of trait clusters based on the absence/presence of data for primate species

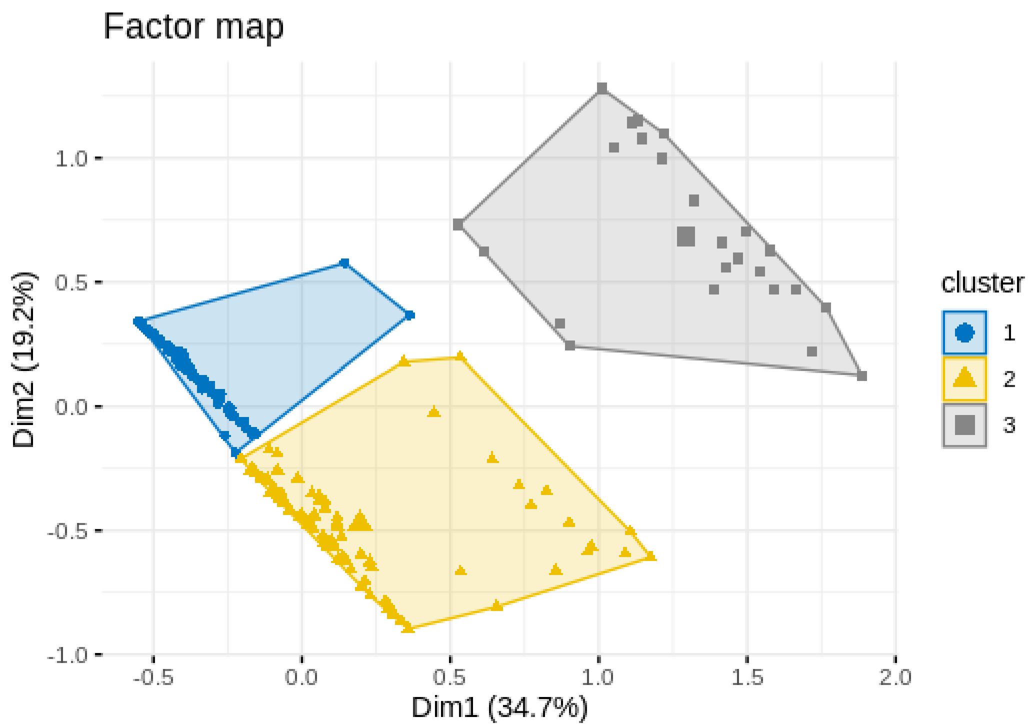


**Fig. S3.**

Representation of 2 first dimensions of the factor map for our set of species clusters based on the absence/presence of data for primate species

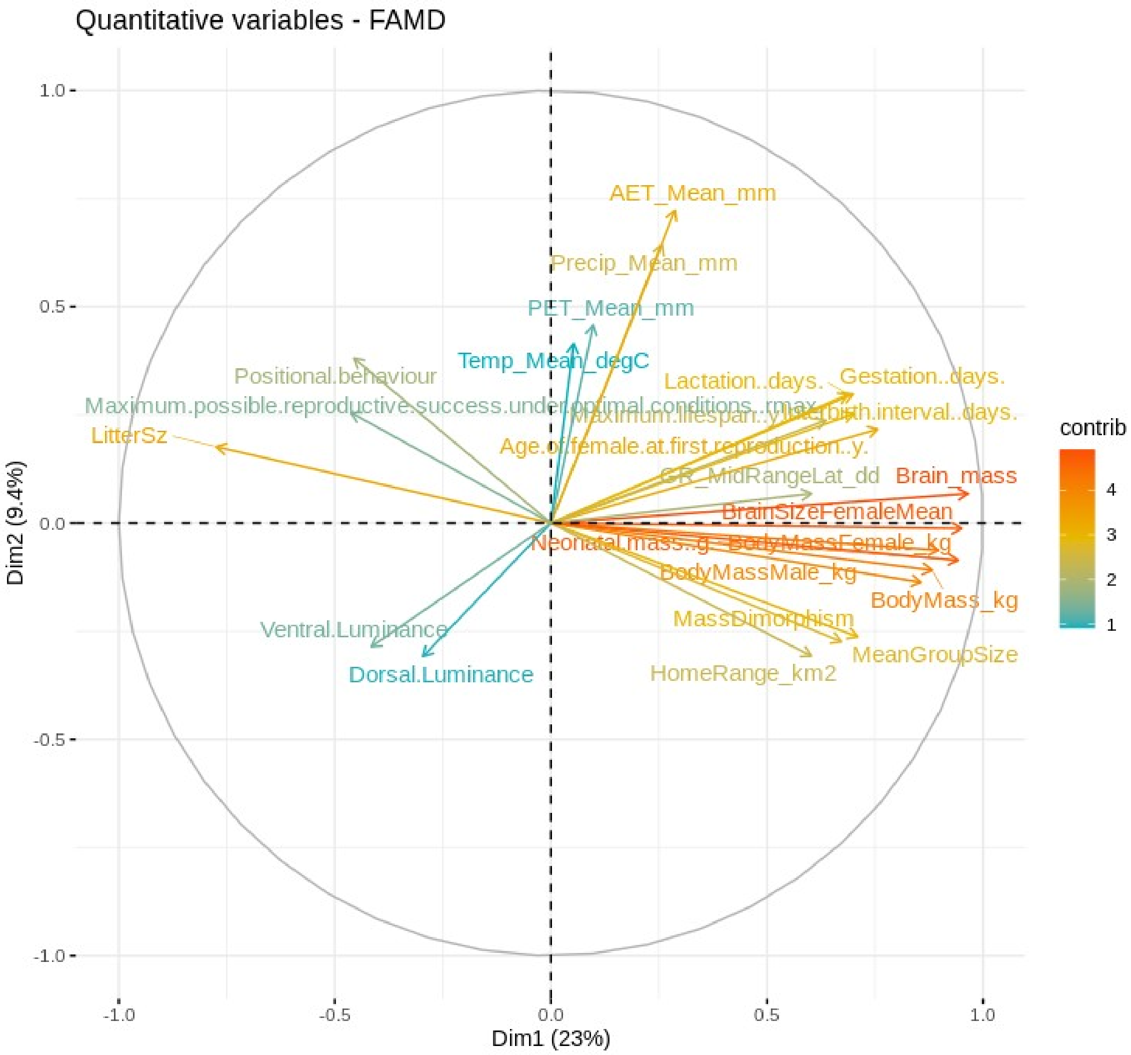


**Fig. S4.**

Cluster VIII contained high coverage life history, ecological, and morphological traits. The variable contributions reported here explain the determination of a given principal component (in percentage) as follows: (var.cos2 * 100) / (total cos2 of the component)

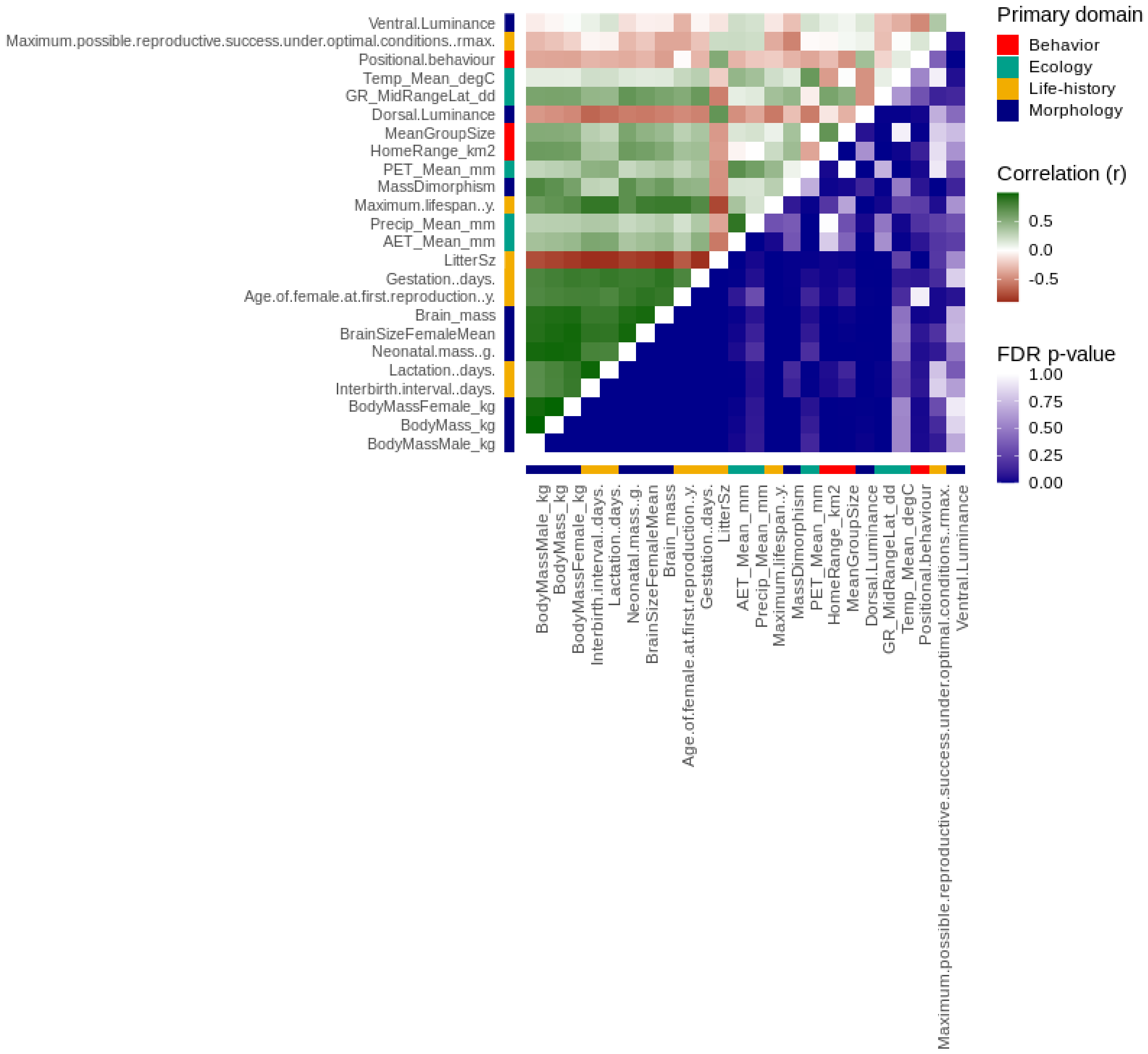


**Fig. S5.**

Cluster VIII comprised high-coverage life-history, ecological, and morphological traits. For each trait pair, the phylogenetic correlation values are shown above the diagonal, and the corresponding FDR-adjusted P-values are shown below the diagonal. The assignment to the primary biological domains is indicated next to the trait label.

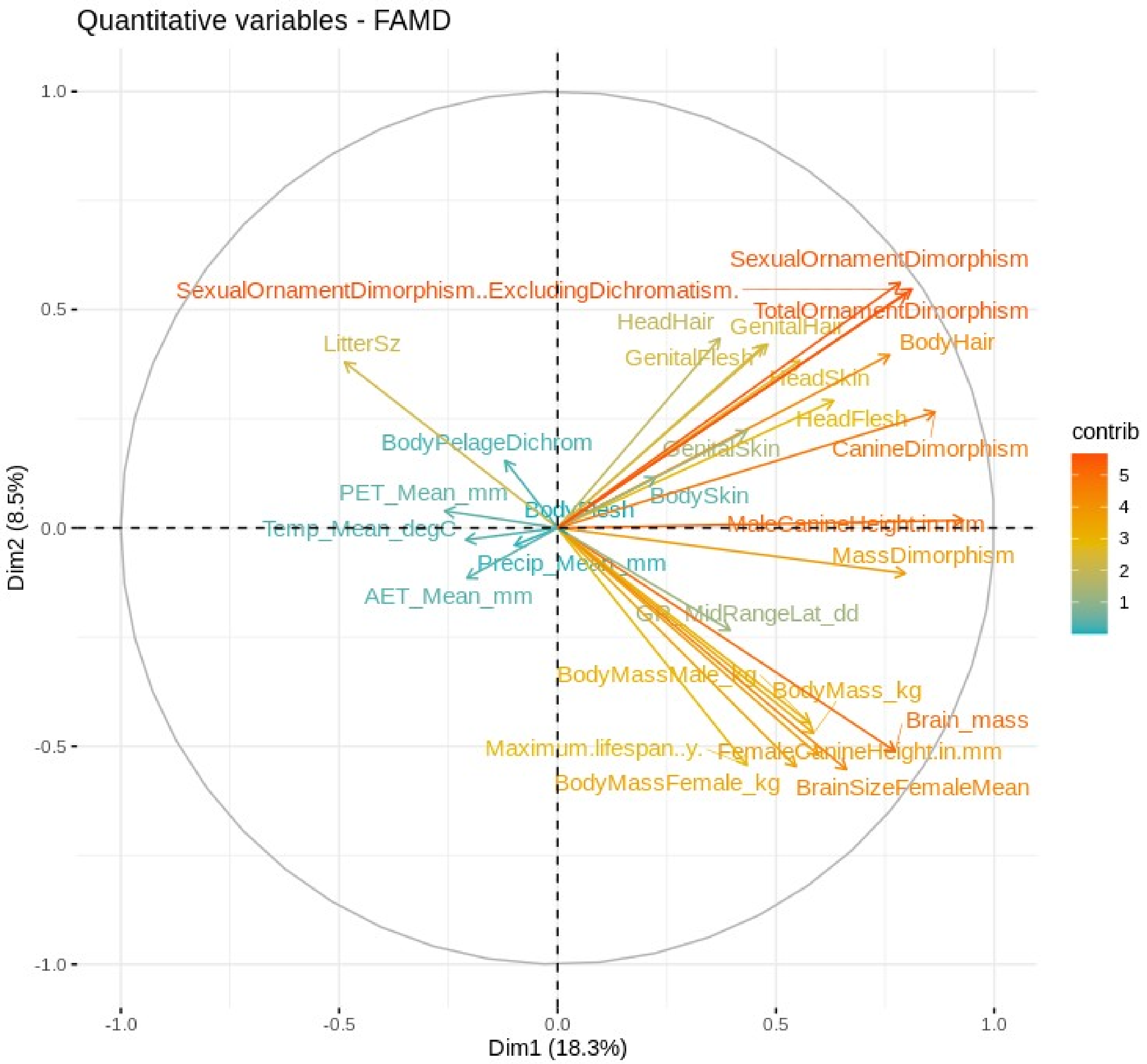


**Fig. S6.**

Cluster VI contained sexually dimorphic traits. The variable contributions reported here explain the determination of a given principal component (in percentage) as follows: (var.cos2 * 100) / (total cos2 of the component)

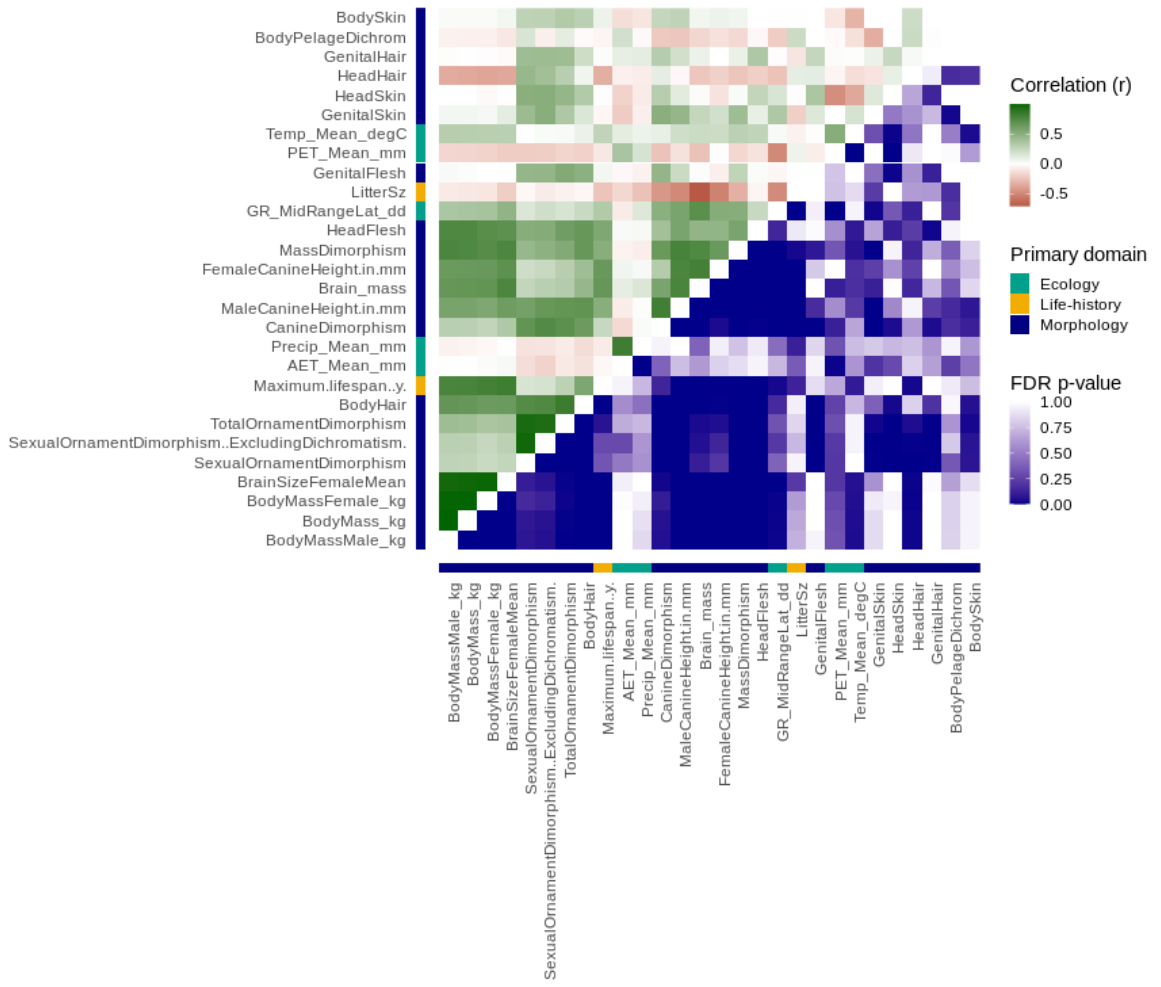


#### Fig. S7.

Cluster VI comprised sexually dimorphic traits. For each trait pair, the phylogenetic correlation values are shown above the diagonal, and the corresponding FDR-adjusted P-values are shown below the diagonal. The assignment to the primary biological domains is indicated next to the trait label.

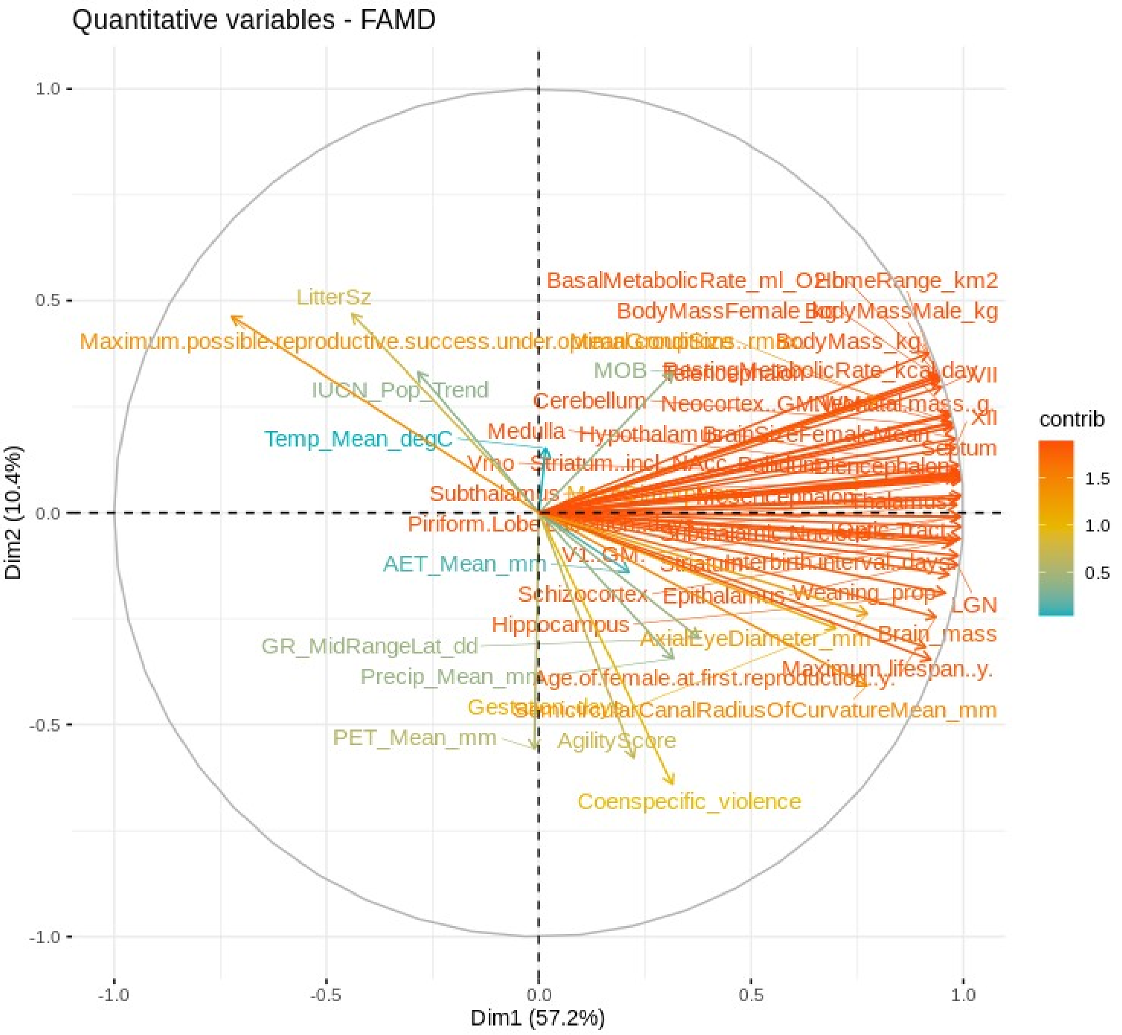


#### Fig. S8.

Cluster V contained brain region-related traits. The variable contributions reported here explain the determination of a given principal component (in percentage) as follows: (var.cos2 * 100) / (total cos2 of the component)

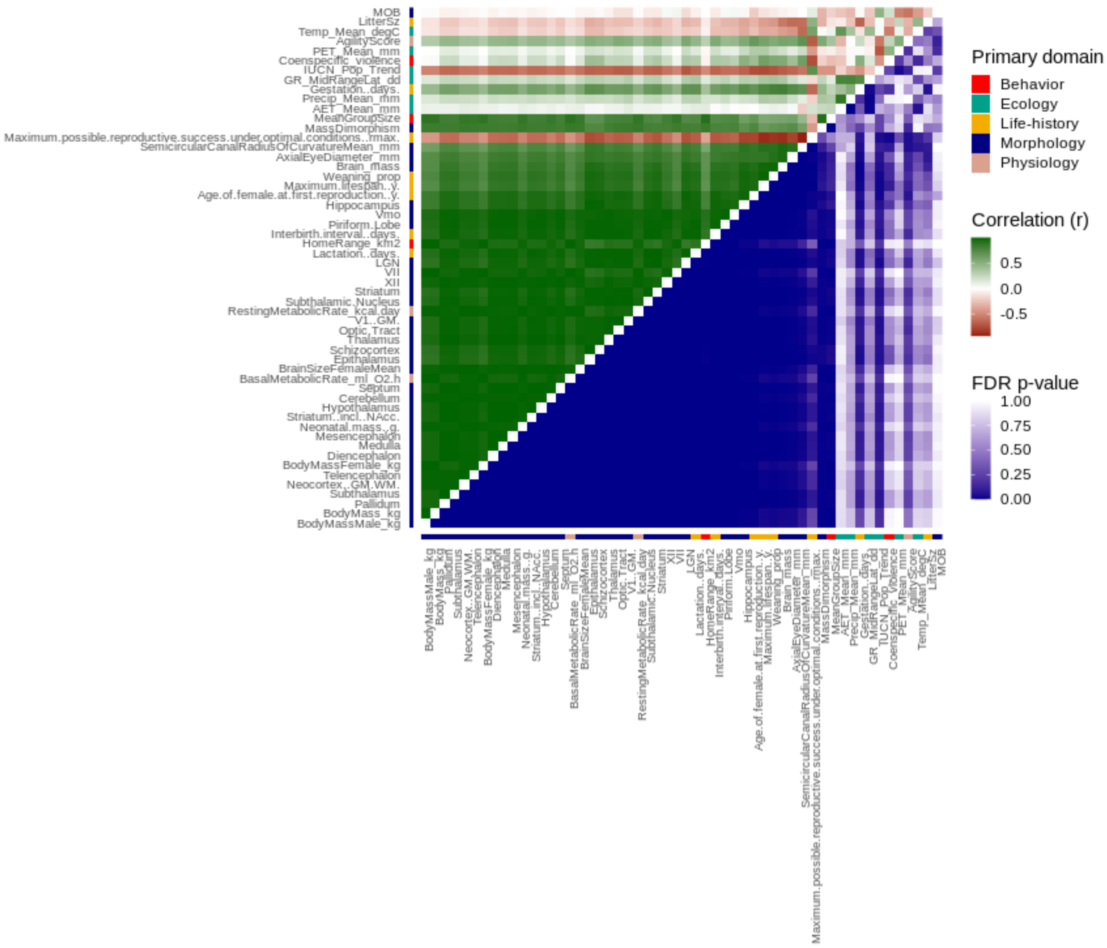


#### Fig. S9.

Cluster V comprised brain region-related traits. For each trait pair, the phylogenetic correlation values are shown above the diagonal, and the corresponding FDR-adjusted P-values are shown below the diagonal. The assignment to the primary biological domains is indicated next to the trait label.

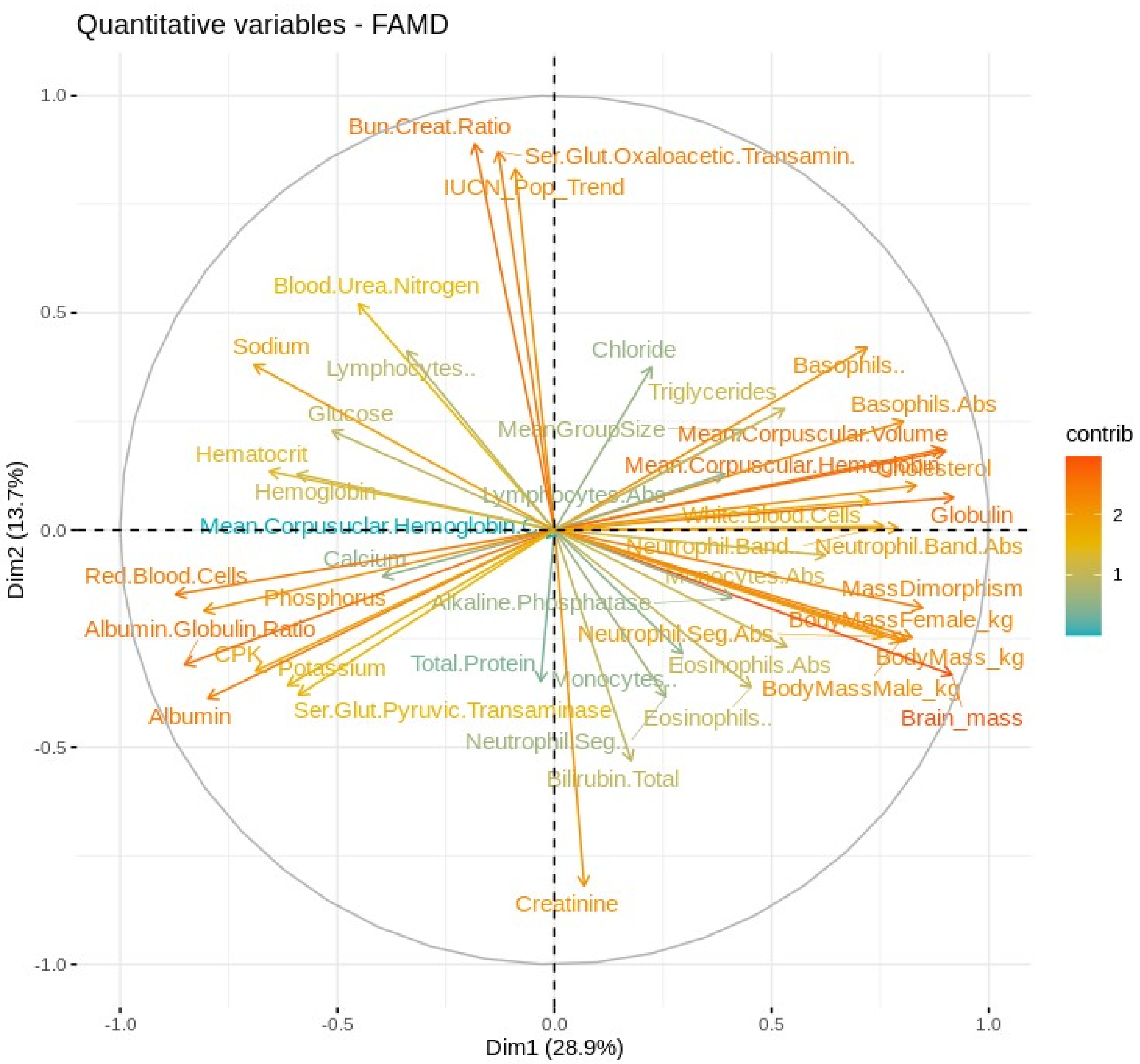


**Fig. S10.**

Cluster IV contained biometabolite traits. The variable contributions reported here explain the determination of a given principal component (in percentage) as follows: (var.cos2 * 100) / (total cos2 of the component)

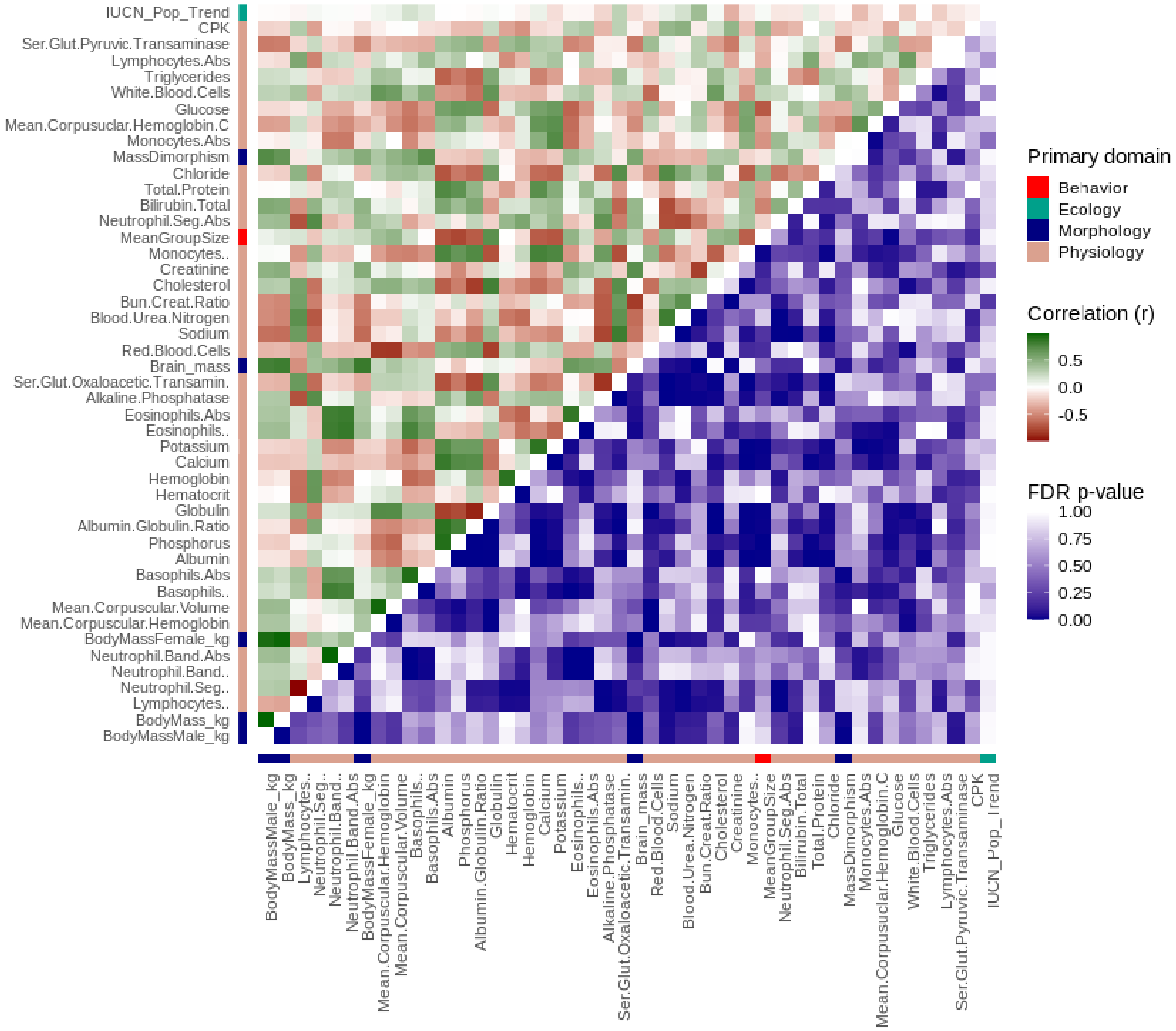


#### Fig. S11.

Cluster IV comprised biometabolite traits. Phylogenetic correlation matrix showing the correlation values for each pair of traits. For each trait pair, the phylogenetic correlation values are shown above the diagonal, and the corresponding FDR-adjusted P-values are shown below the diagonal. The assignment to the primary biological domains is indicated next to the trait label.

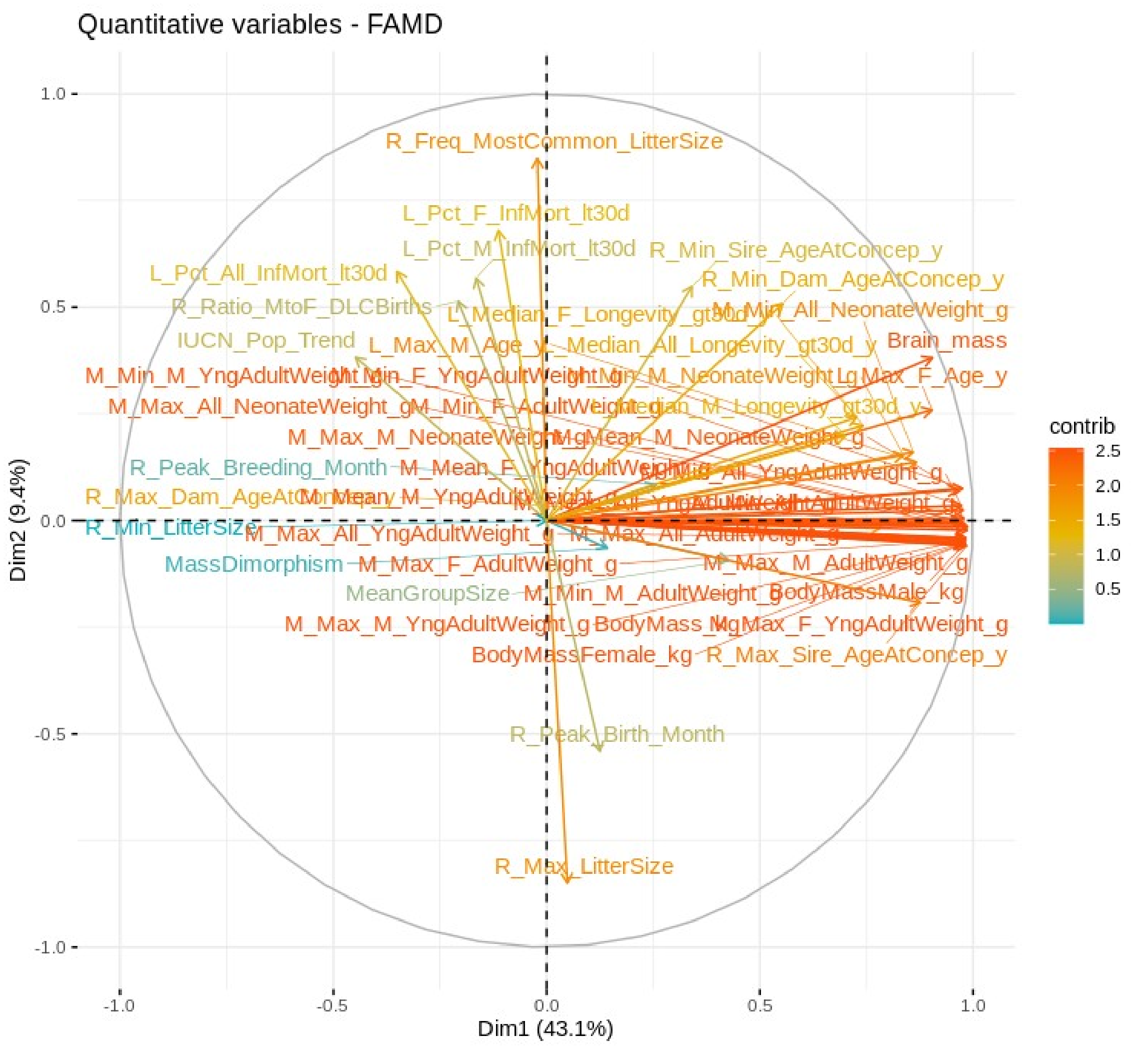


#### Fig. S12.

*Cluster I*. Contained life-history traits mainly from *Duke Lemur Center*. The variable contributions reported here explain the determination of a given principal component (in percentage) as follows: (var.cos2 * 100) / (total cos2 of the component)

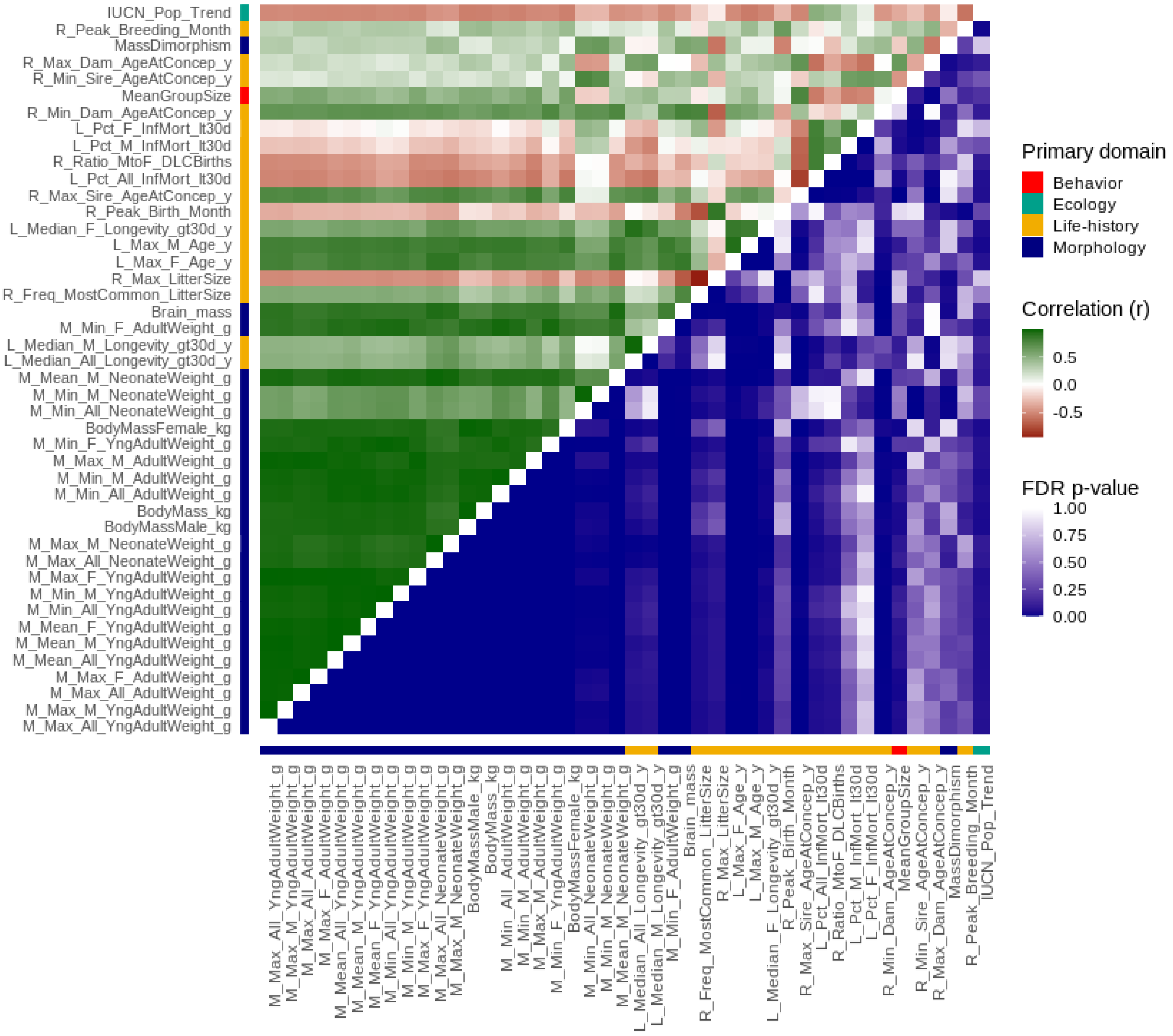


**Fig. S13.**

Cluster I comprised life-history traits mainly from the *Duke Lemur Center*. Phylogenetic correlation matrix showing the correlation values for each pair of traits. For each trait pair, phylogenetic correlation values are shown above the diagonal and the corresponding FDR-adjusted P-values are shown below the diagonal. The assignment to the primary biological domains is indicated next to the trait labels.

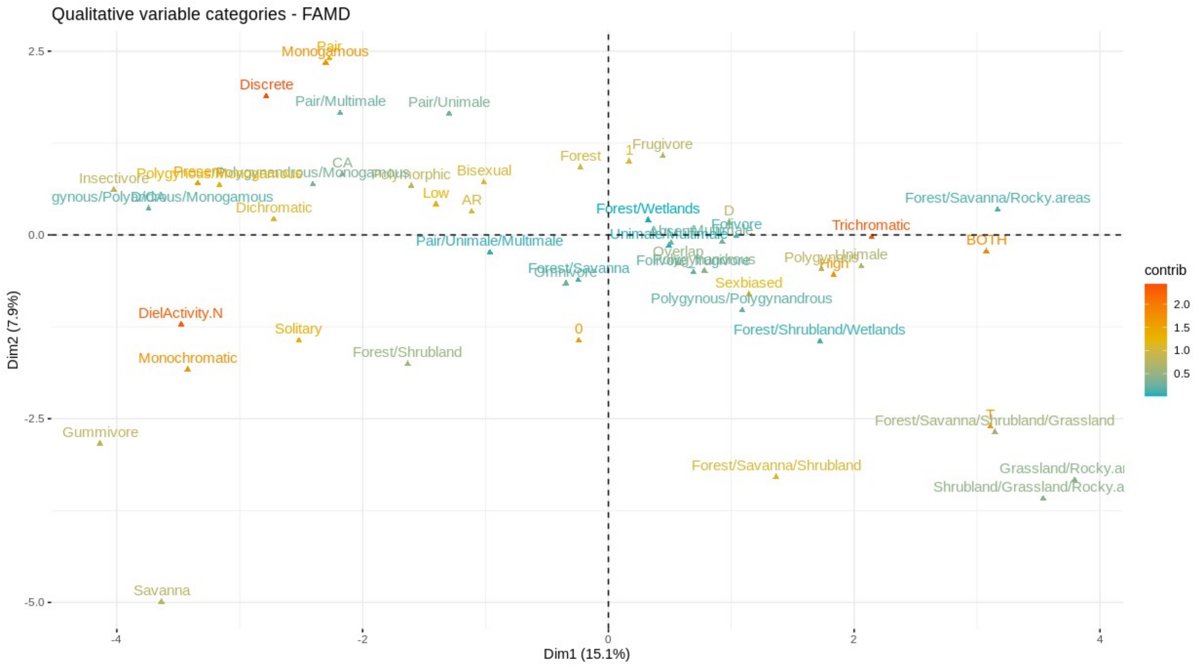

#### Fig. S14.

Cluster X (qualitative) contained qualitative binary and multinomial traits. The variable contributions reported here explain the determination of a given principal component (in percentage) as follows: (var.cos2 * 100) / (total cos2 of the component)

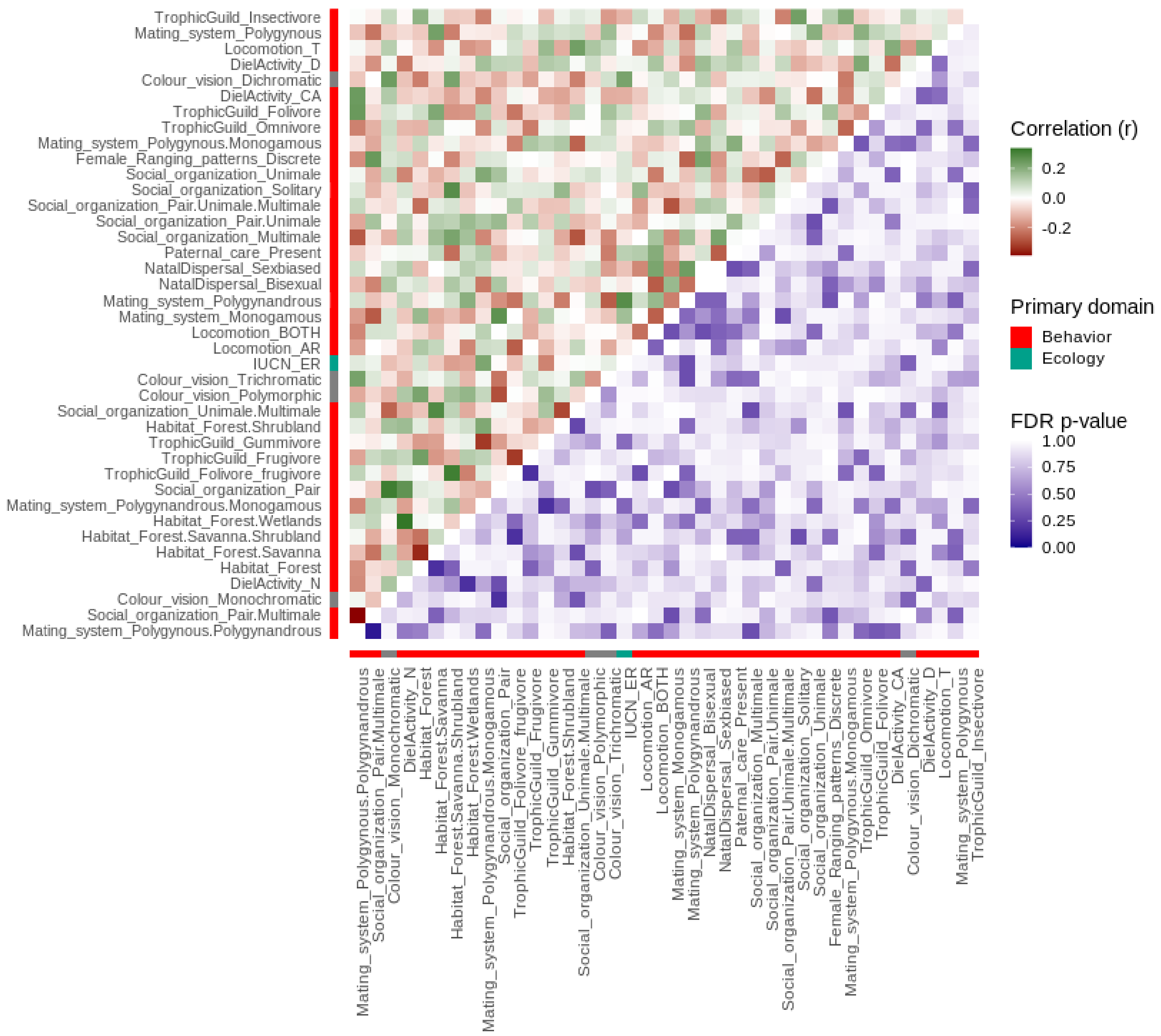


#### Fig. S15.

Cluster X (qualitative) comprised qualitative binary and multinomial traits containing liability correlations. Phylogenetic correlation matrix showing the correlation values for each pair of traits. For each trait pair, the phylogenetic correlation values are shown above the diagonal, and the corresponding FDR-adjusted P values are shown below the diagonal. The assignment to the primary biological domains is indicated next to the trait label.

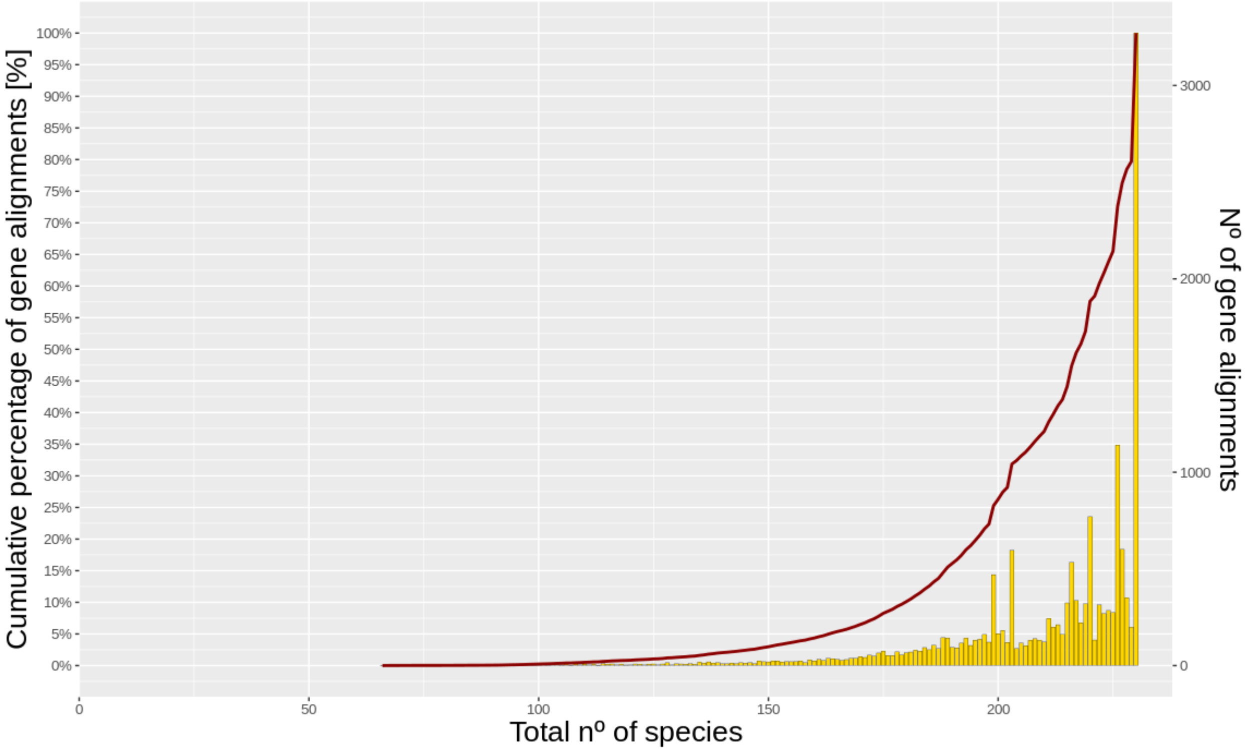


**Fig. S16.**

Histogram of the number of primate species (x) per protein-coding gene alignment (y). The red line indicates the cumulative distribution of the total number of gene alignments.

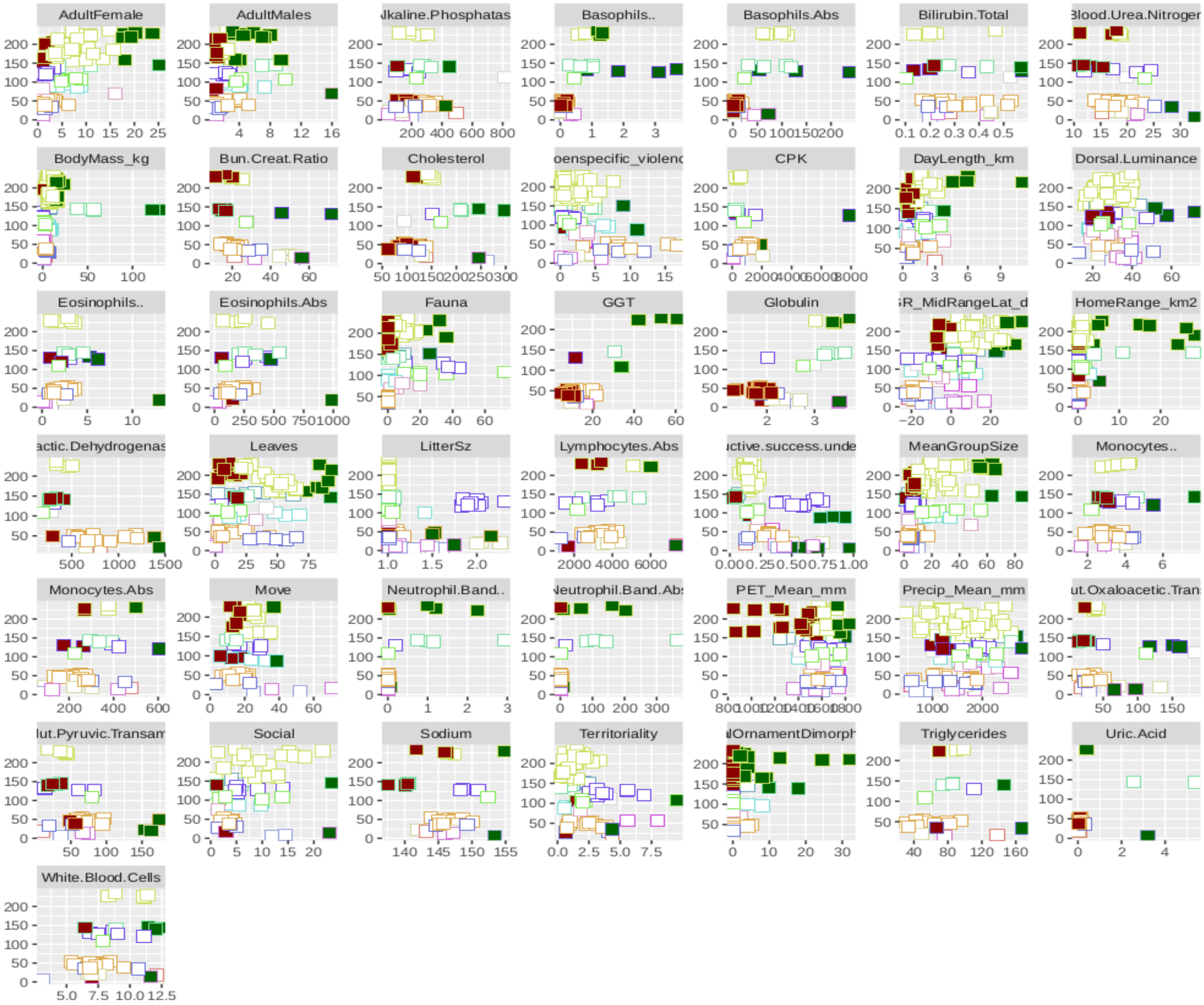

**Fig. S17.**

*Top* and *Bottom* binarizations for quantitative traits non-allometrically corrected (green for *Top* species and red for *Bottom* species) after the selection criteria.

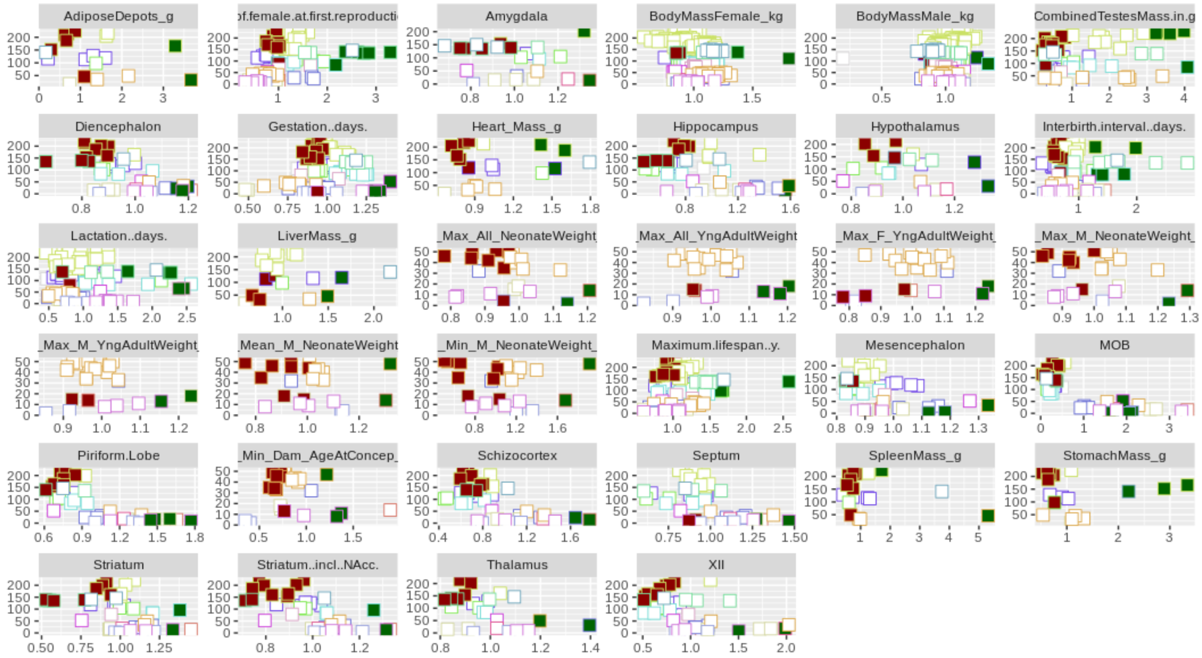


**Fig. S18.**

*Top* and *Bottom* binarizations for quantitative traits were allometrically corrected (green for *Top* species and red for *Bottom* species) after the selection criteria.

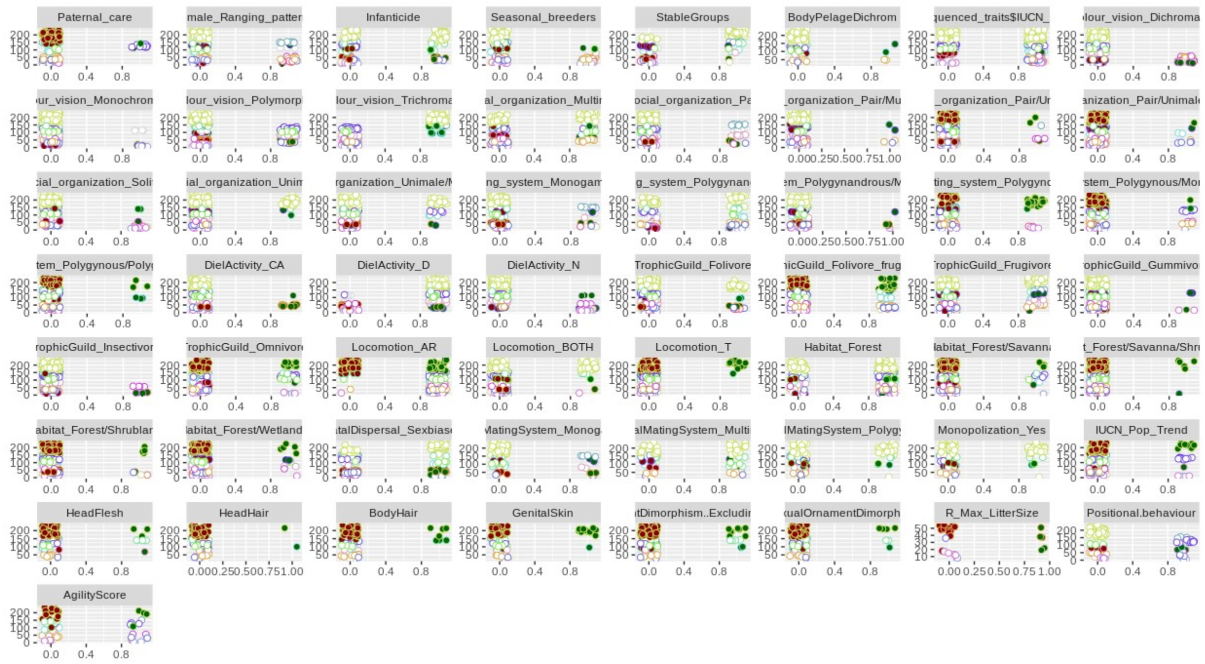

**Fig. S19.**

*Top* and *Bottom* binarizations for qualitative traits (green for *Top* species and red for *Bottom* species) after the selection criteria

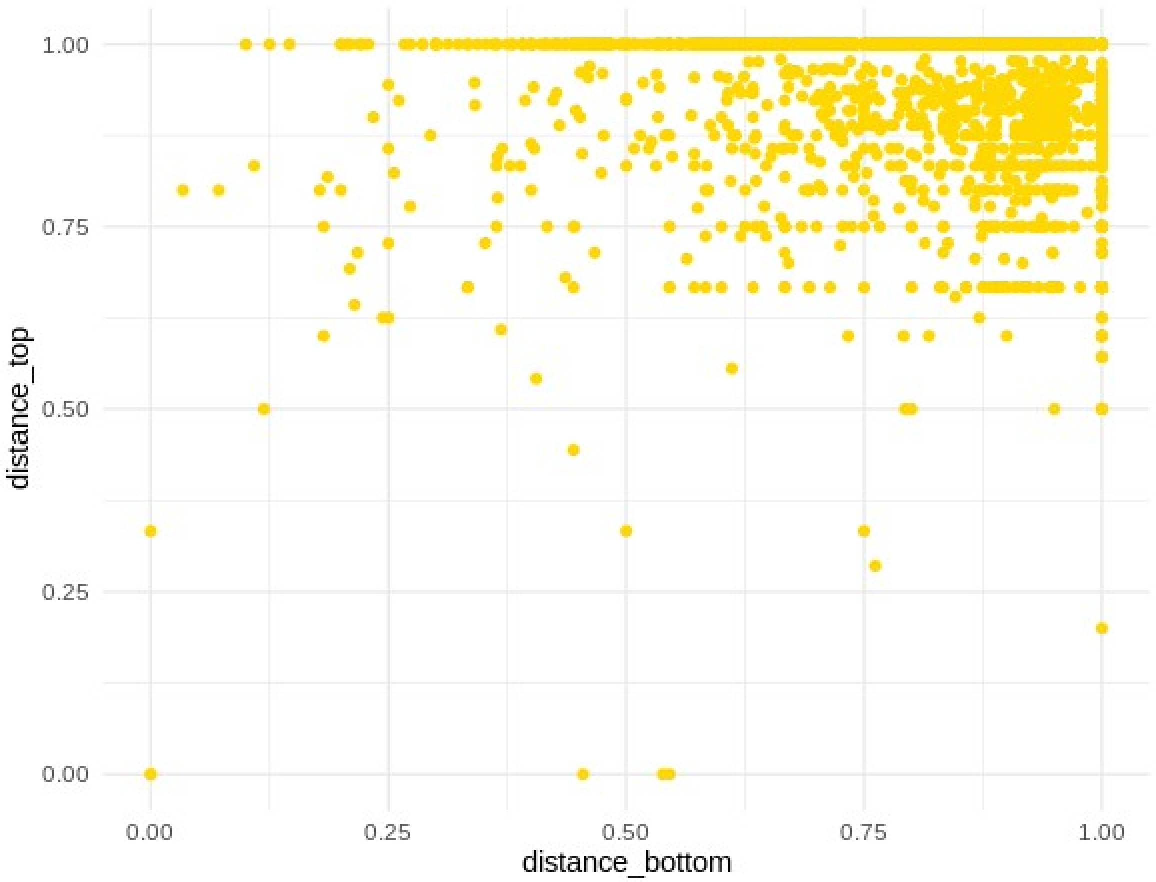


**Fig. S20.**

The corresponding distance between *Top* species and *Bottom* species from our trait selection for CAAS between each pair of traits with extremes was selected. Distance is measured as 1-*Jaccard similarity* metric.

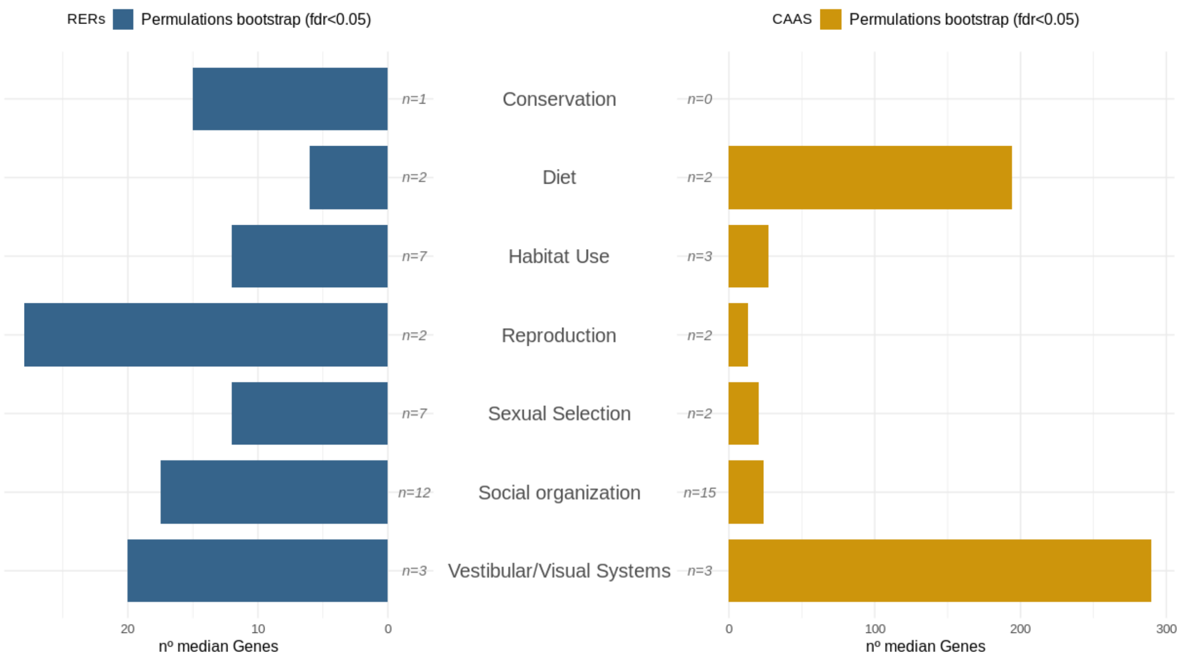


### Fig. S21.

Median number of genes with significant p-value after multiple-gene test correction (FDR<0.05) by the main biological domains for each phylogenetic test applied and each corresponding qualitative trait in the P3GMap.

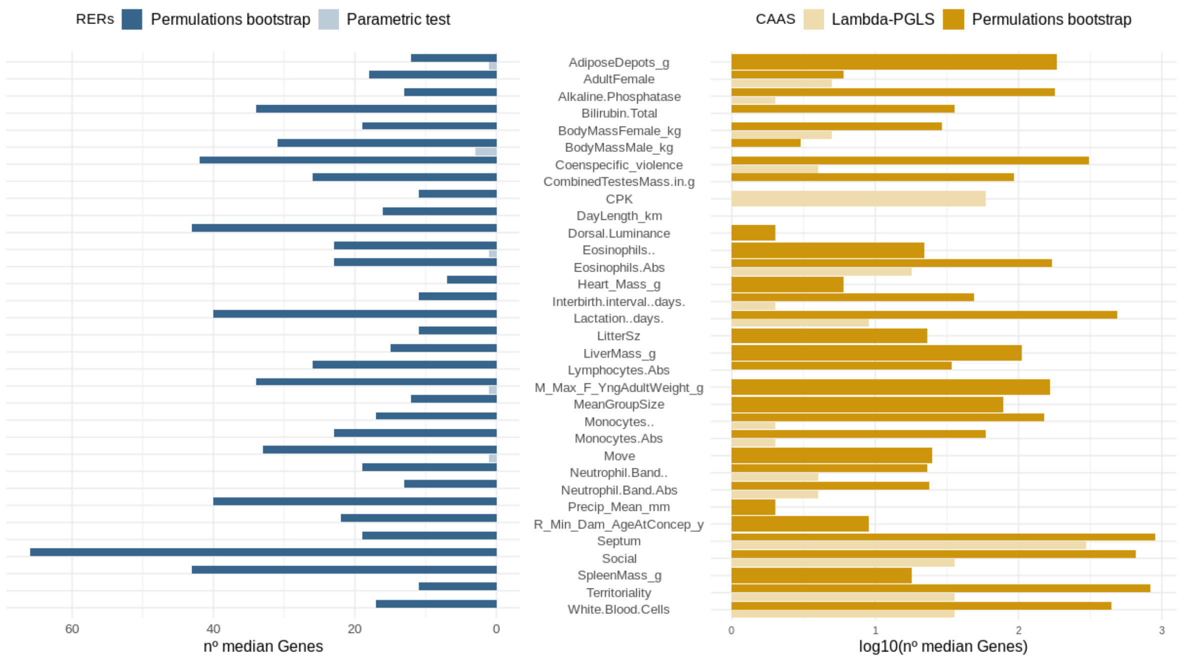


### Fig. S22.

Number of genes with significant p-values after multiple-gene test correction (FDR<0.05) per trait for each phylogenetic test applied and each corresponding quantitative trait with >=2 families with contrasting phenotypes in the P3GMap

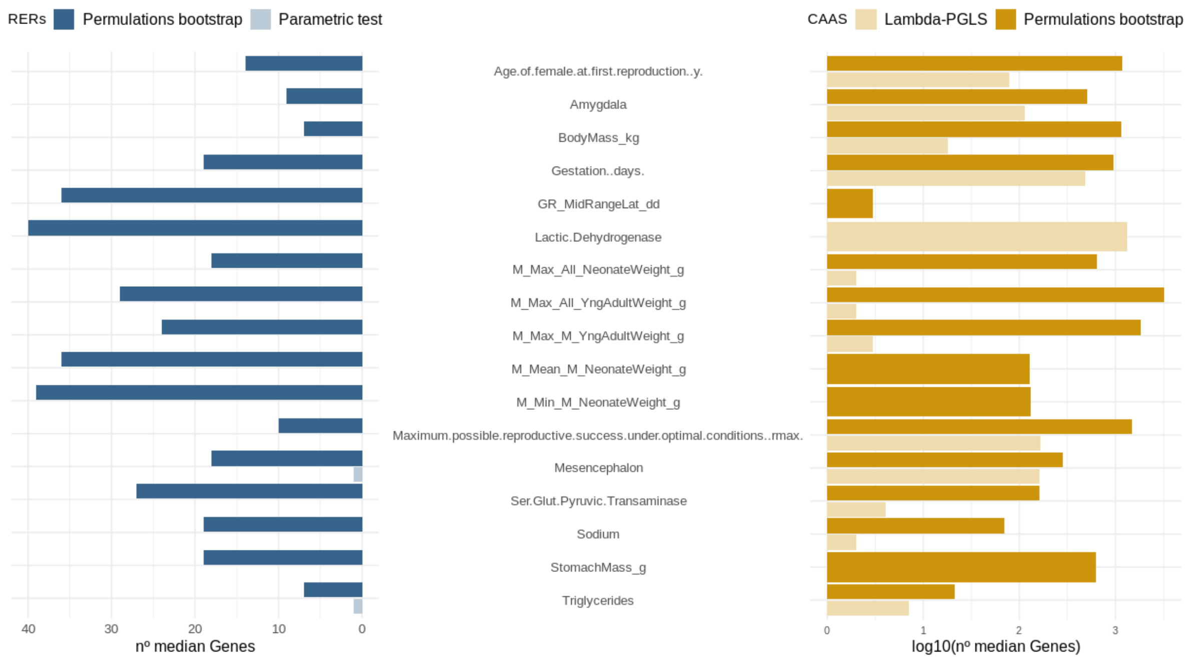


**Fig. S23.**

Number of genes with significant p-values after multiple-gene test correction (FDR<0.05) per trait for each phylogenetic test applied and each corresponding quantitative trait with only one family with contrasting phenotypes in the P3GMap

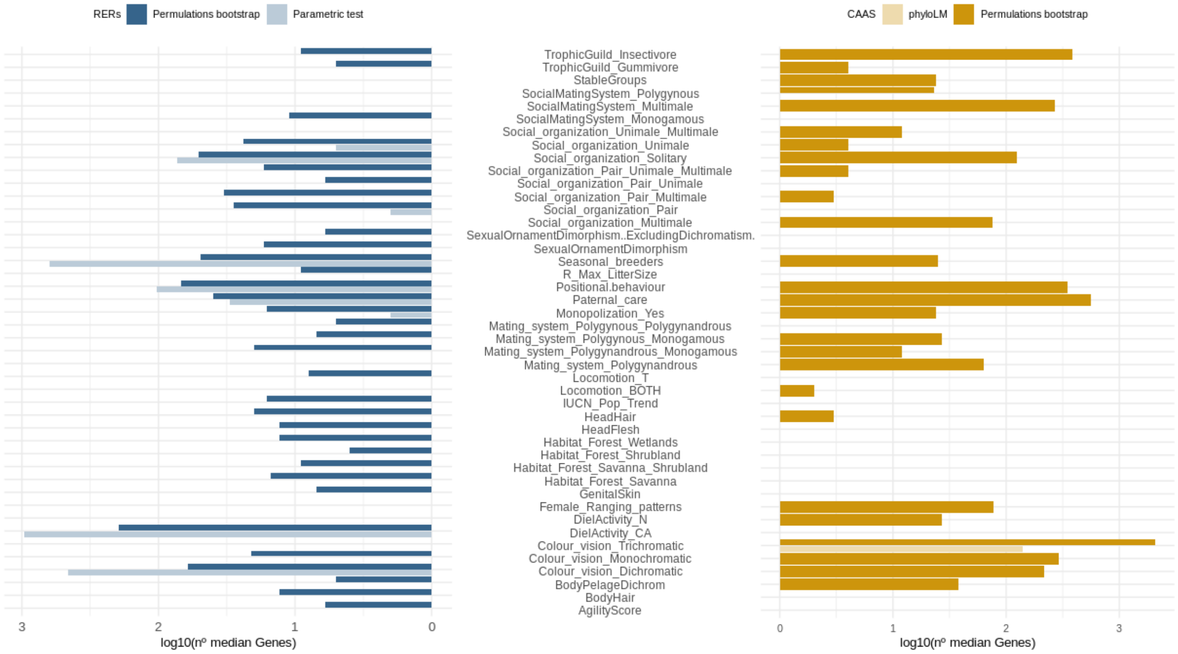


### Fig. S24.

Number of genes with significant p-values after multiple-gene test correction (FDR<0.05) for each phylogenetic test applied and each corresponding qualitative trait in the P3GMap

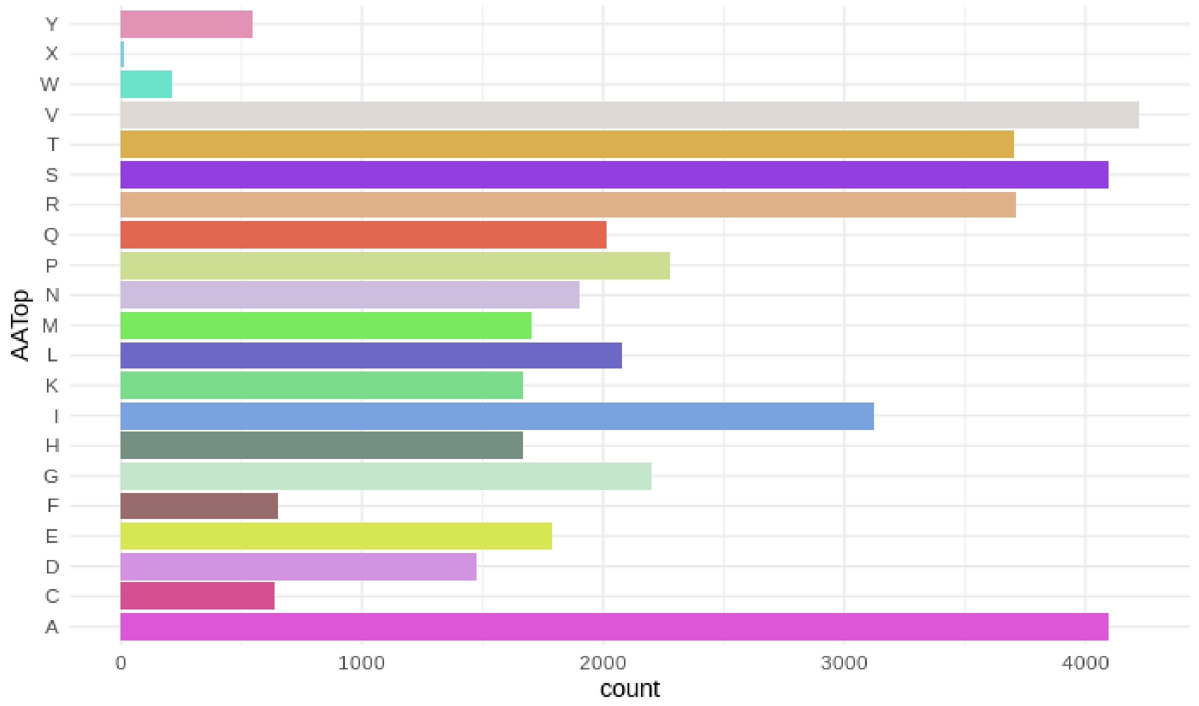


**Fig. S25.**

Frequency for each AA in *Top* species with positions with perm p-value<0.05 for all our set of traits with CAAS in the P3GMap

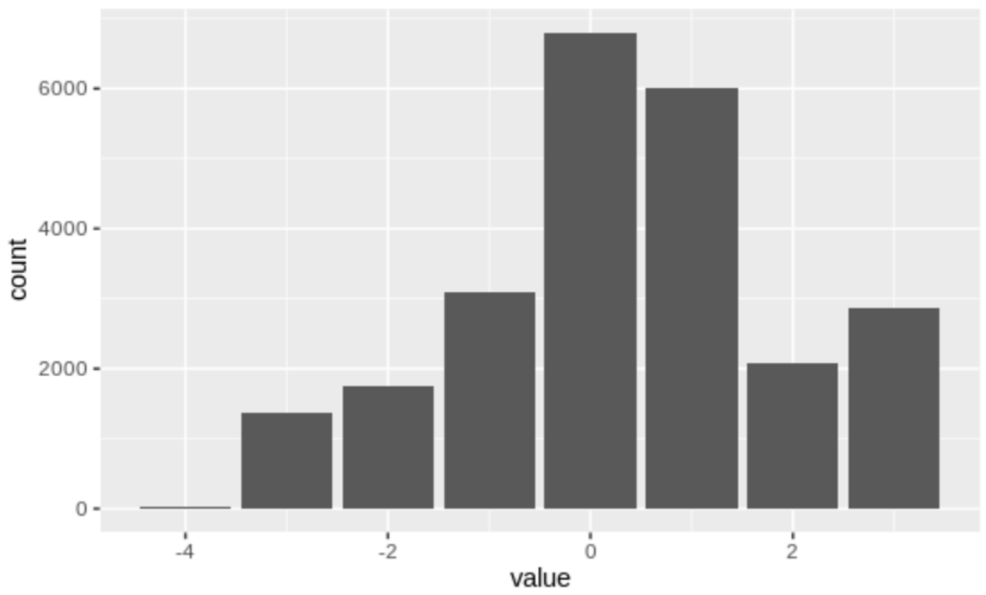


**Fig. S26.**

Amino Acid transition scores from BLOSUM62 considering CAASs recovered for Scenario1 and perm p-value < 0.05 in the P3GMap

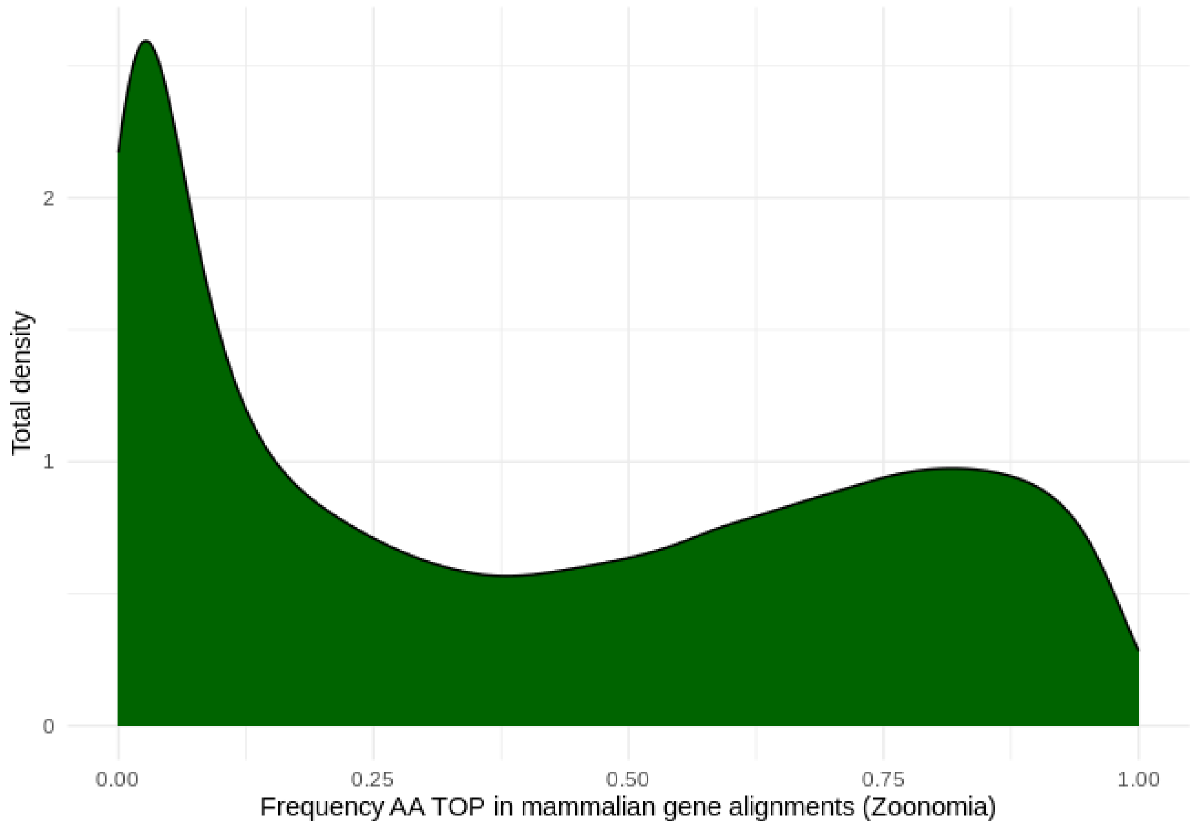


**Fig. S27.**

Density distribution of the frequency of *Top* AA detected in primates with CAAS in the Zoonomia mammalian protein-coding gene alignments for each respective position in the P3GMap.

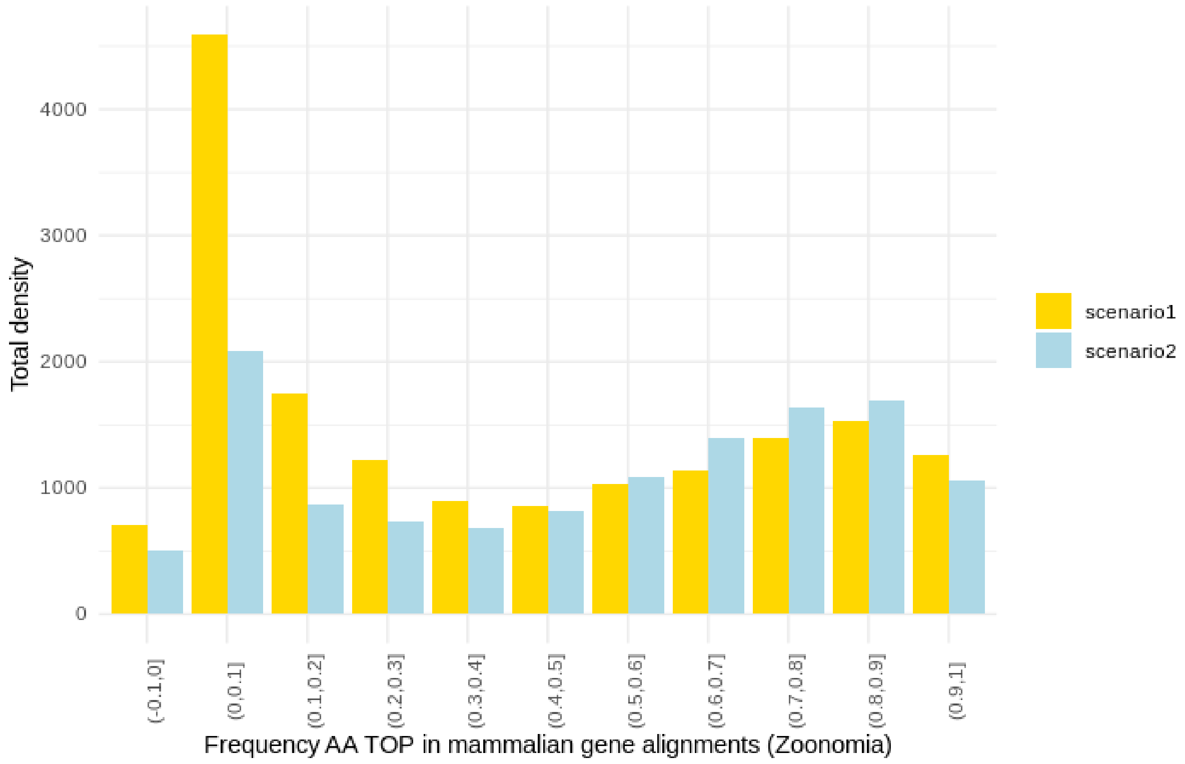


**Fig. S28.**

Histogram for the 10-deciles frequency of *Top* AA detected in primates with CAAS in the Zoonomia mammalian protein-coding gene alignments for each respective position with perm-p-value <0.05 and each respective binarized substitution *Scenario* considered in the P3GMap.

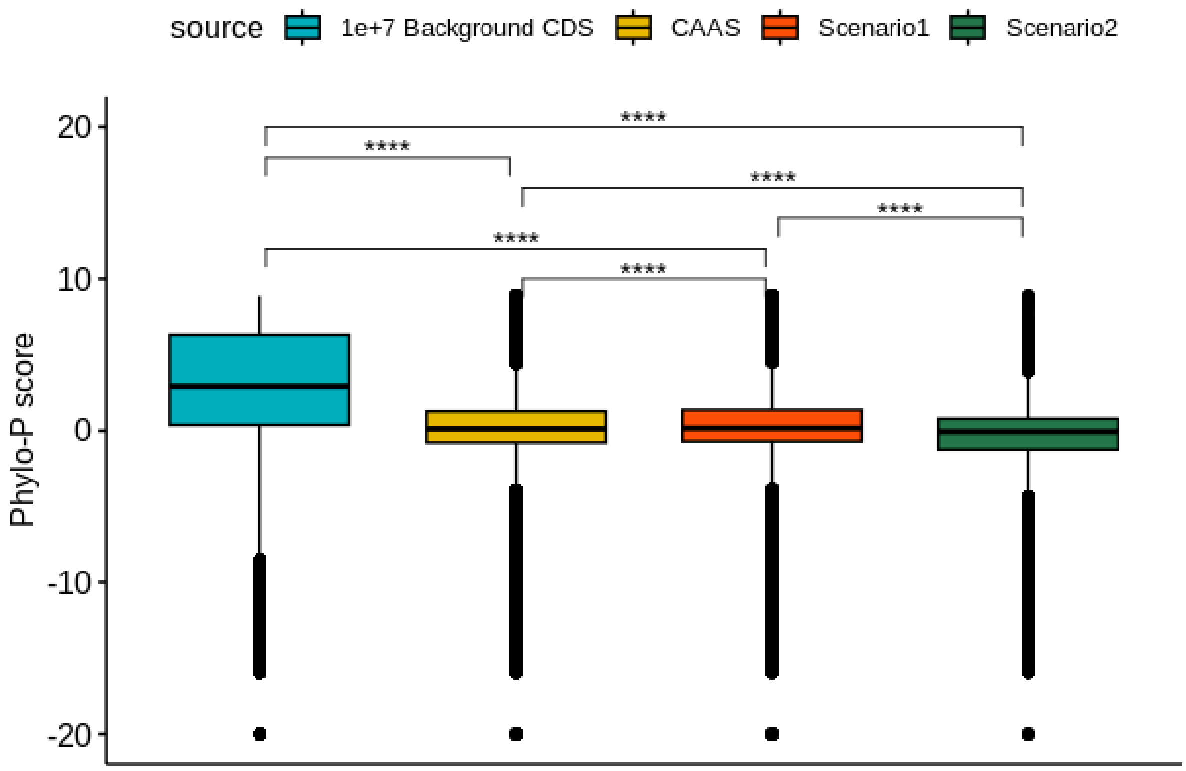


**Fig. S29.**

Boxplot showing PhyloP conservation score (from *Zoonomia*) for recovered positions with CAAS (both *scenario1* and *scenario2*) with perm-p-value <0.05 against random 10M background CDS positions in primate protein-coding genes.

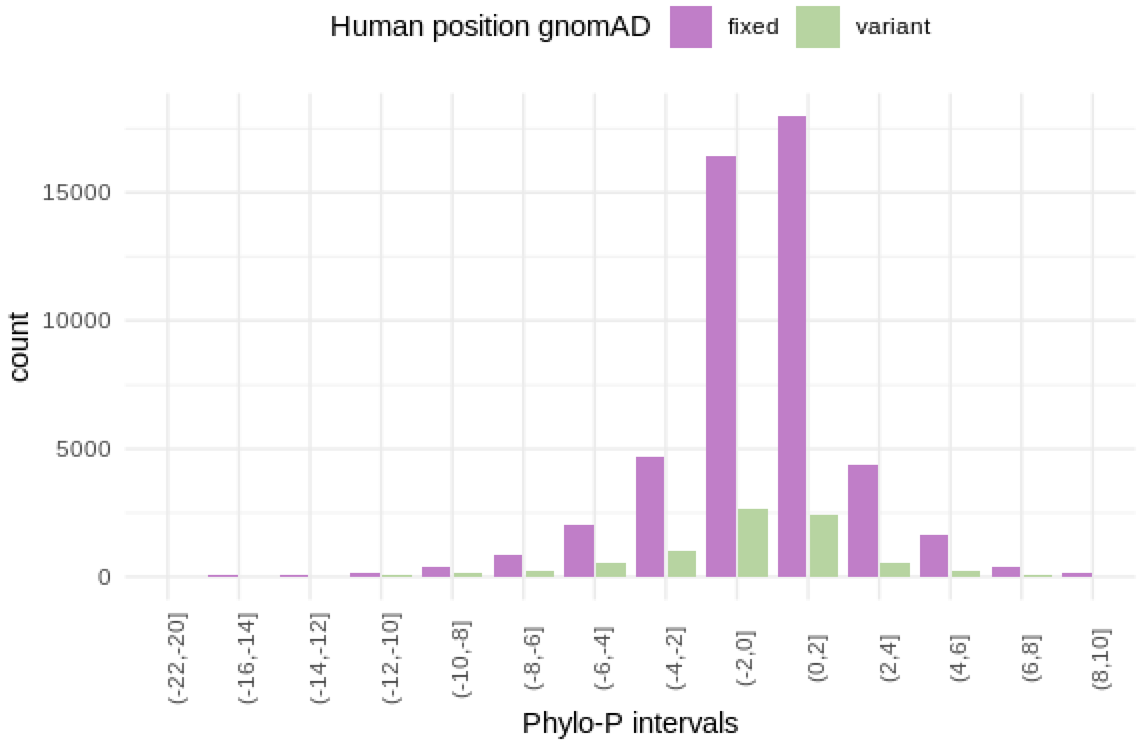


#### Fig. S30.

The alternative allele frequency (AF) of CAAS-recovered positions was determined using gnomAD 3.1, including biallelic variant positions in the analysis and splitting information into intervals according to the respective PhyloP score assigned to each position in the P3GMap.

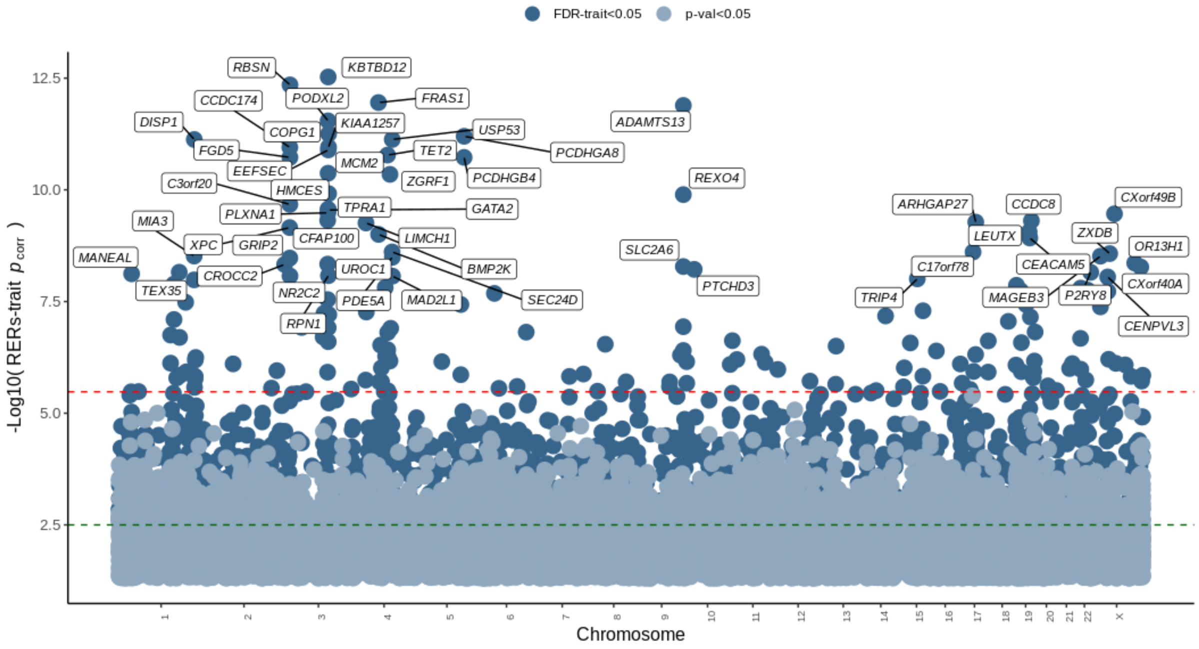


#### Fig. S31.

Manhattan plot of RERs gene-trait qualitative associations for the parametric test in P3GMap. The dashed green line corresponds to the FDR threshold, and the dashed red line corresponds to the Bonferroni thresholds. Genes labelled with their corresponding gene names have ($p_{corr}$<1e-8).

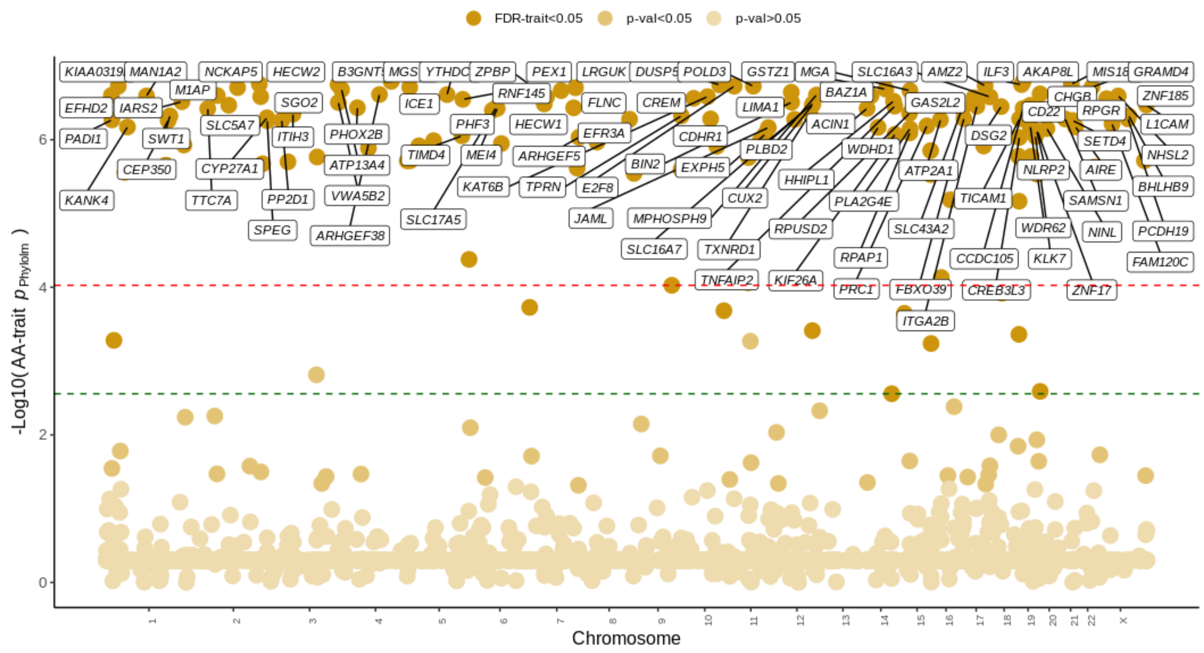


**Fig. S32.**

Manhattan plot for CAAS gene-trait qualitative associations after computing the lowest p-value per trait from the phylogenetic test applied in the P3GMap. The dashed green line corresponds to the FDR threshold, and the dashed red line corresponds to the Bonferroni thresholds Genes labeled with their corresponding gene names had a -log10 p-value > 4.

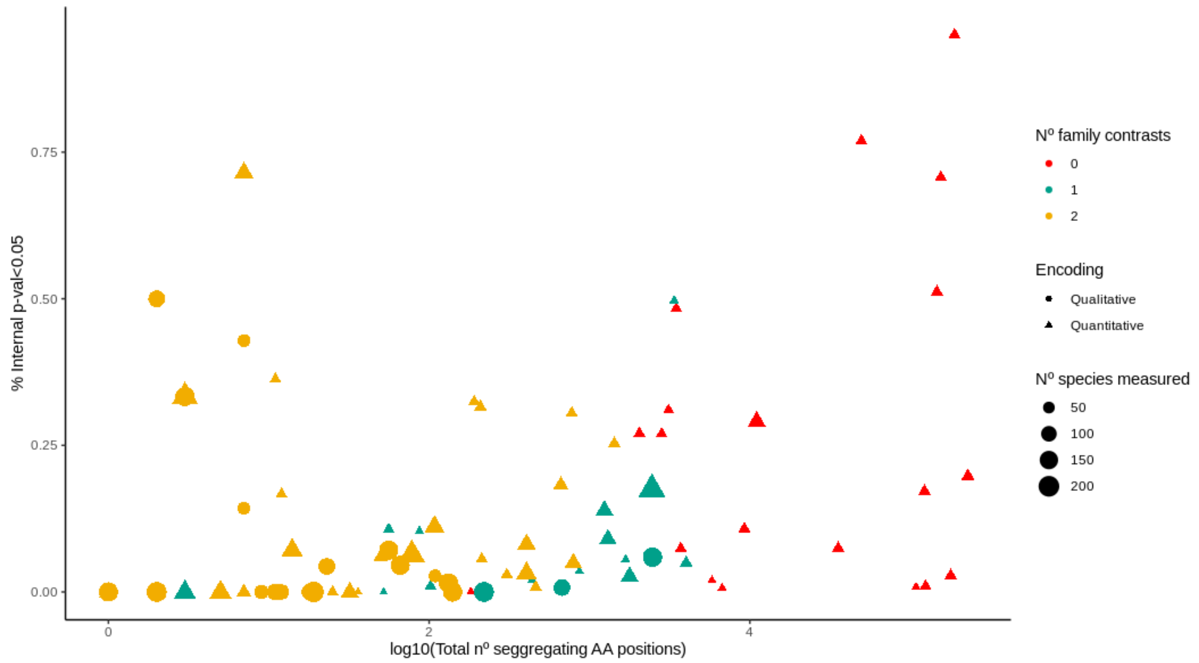


**Fig. S33.**

Descriptive CAAS plot between the total number of CAAS segregating positions (>5 *Top* & *Bottom* species) recovered per trait (in *log10)* (x) and % of internally significant positions (y), accounting for the total number of species measured (per trait basis) and classifying them according to A) encoding type (shape), and B) number of *family contrasts* (color).

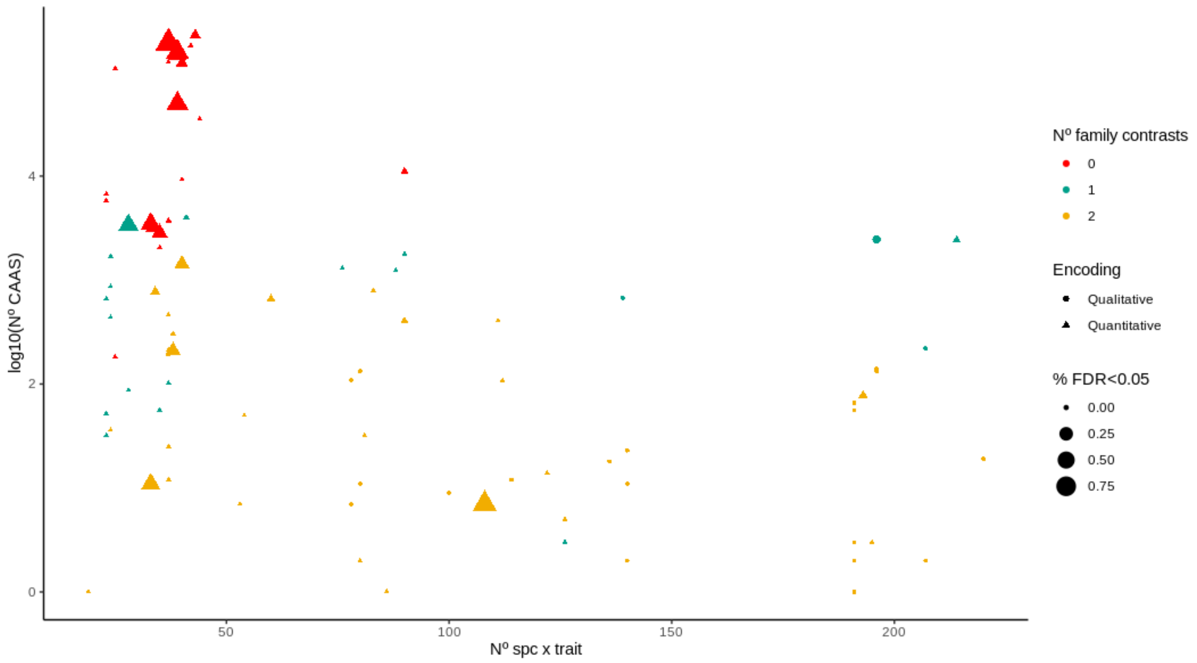


**Fig. S34.**

Descriptive CAAS plot between the total number of CAAS positions recovered per trait (y) and the number of species tested per trait (x), accounting for the proportion of positions with internal test FDR p-value <0.05 (per trait basis) and classifying them according to A) encoding type (shape) and B) number of *family contrasts* (color).

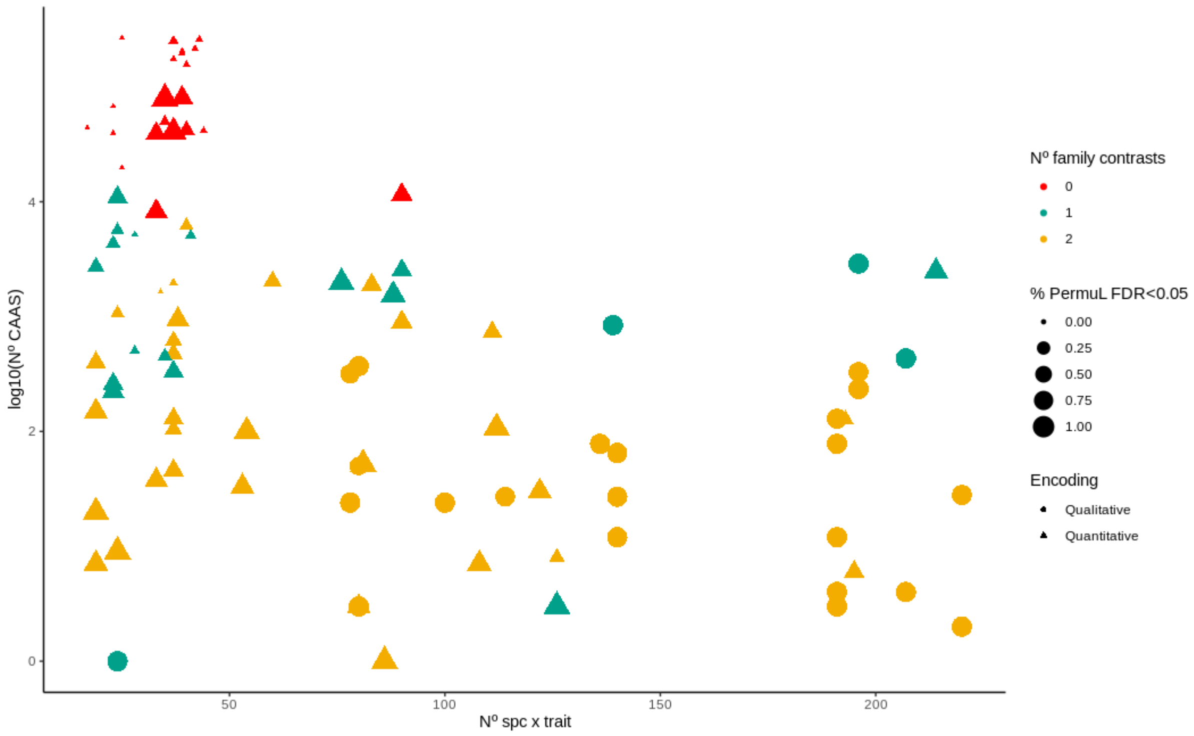


**Fig. S35.**

Descriptive CAAS plot between the total number of CAAS positions recovered per trait (y) and the number of species tested per trait (x), accounting for the proportion of positions with permulations FDR p-value <0.05 (per trait basis) and classifying them according to A) encoding type (shape) and B) number of *family contrasts* (color).

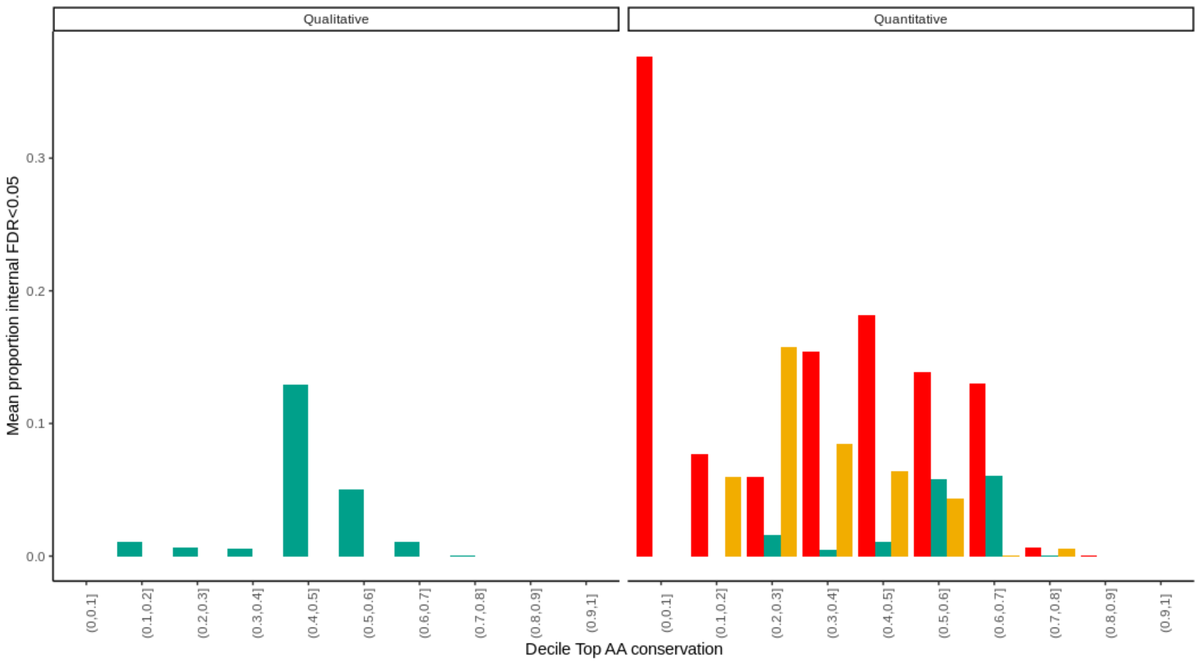


**Fig. S36.**

Barplot of the mean number of CAAS positions with primate consistency test FDR<0.05 (by trait basis) (y) against deciles of the respective range of AA conservation scores (in percentage %) (x) and classifying them according to A) encoding type (facet) and B) number of *family contrasts* (red=0, turquoise=1, gold=2).

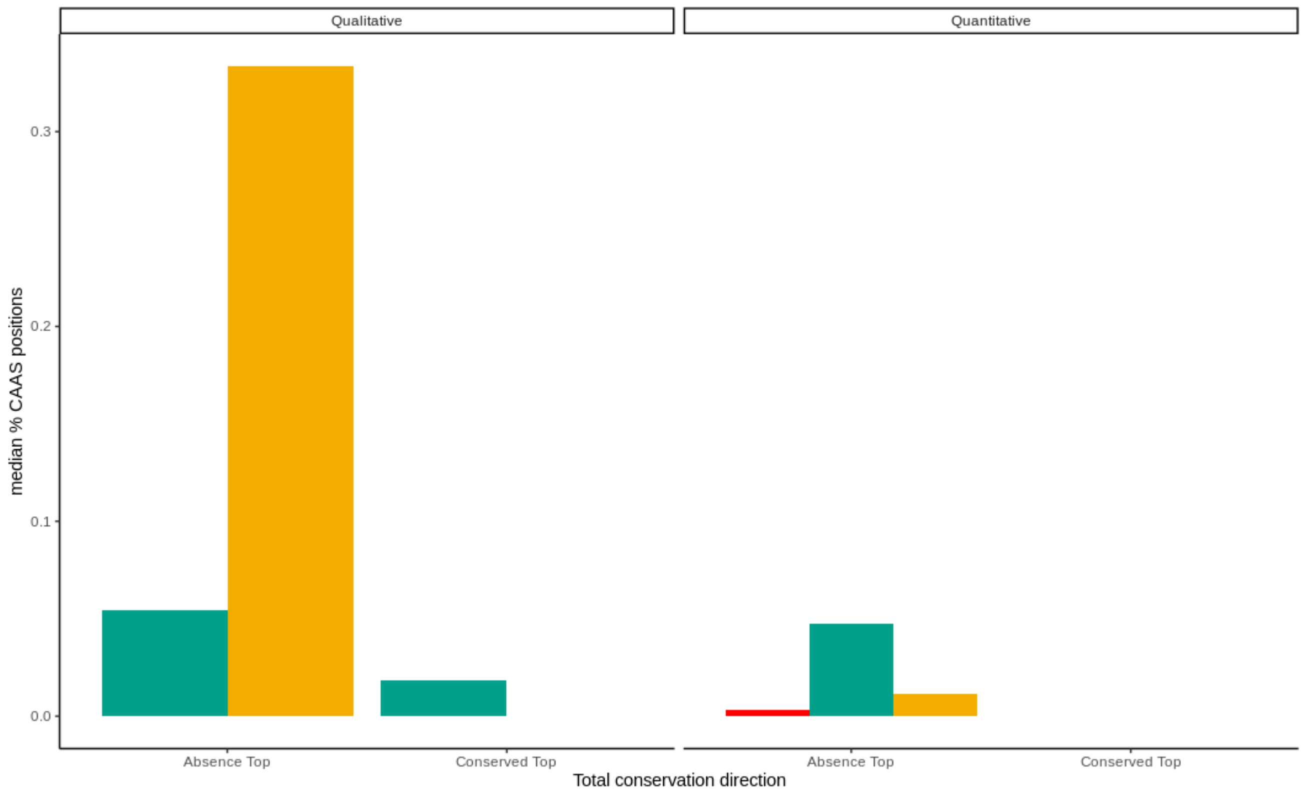


**Fig. S37.**

Barplot comparison of the median number of CAAS positions lacking internal species with *Top AA* (absence *Top*) or *Bottom AA* (conserved *Top*) and classifying them according to A) encoding type (facet) and B) number of *family contrasts* (red=0, turquoise=1, gold=2). These correspond to amino acid positions with CAAS detection but without sufficient power for primate consistency tests.

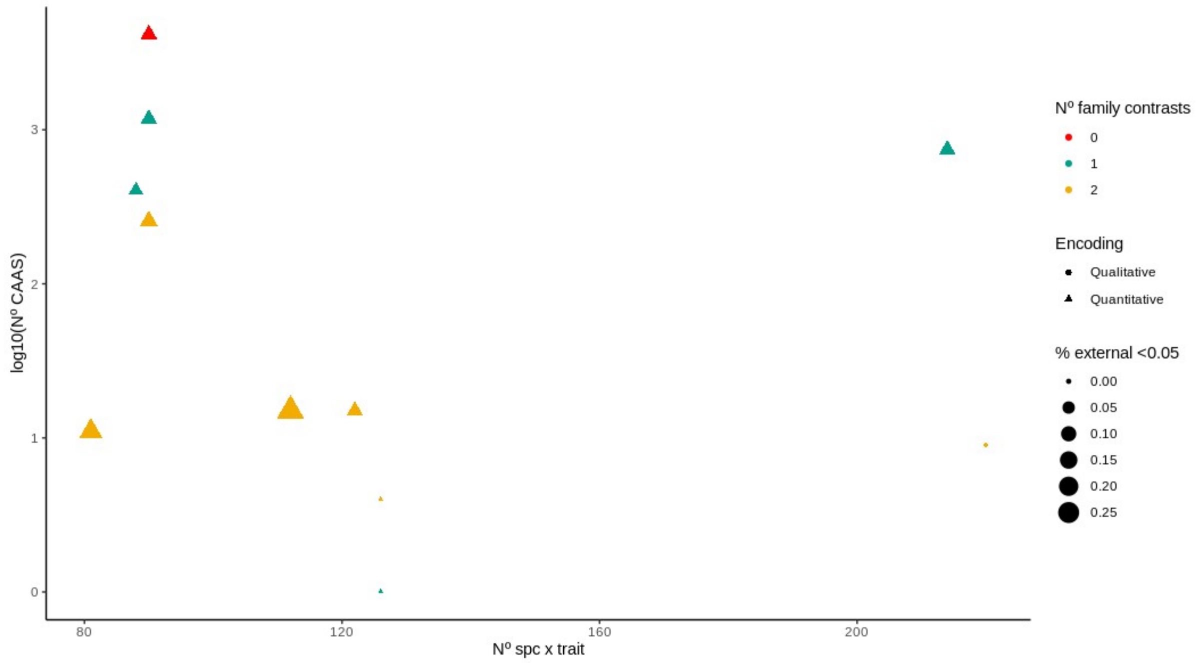


**Fig. S38.**

Descriptive CAAS plot between the total number of CAAS positions recovered per trait (y) and the number of species tested per trait (x), accounting for the proportion of positions with cross-mammalian consistency test p-value <0.05 (per trait basis and >5 member species in both *the Top* and *Bottom* groups) and classifying them according to A) encoding type (shape) and B) number of *family contrasts* (red=0, turquoise=1, gold=2).

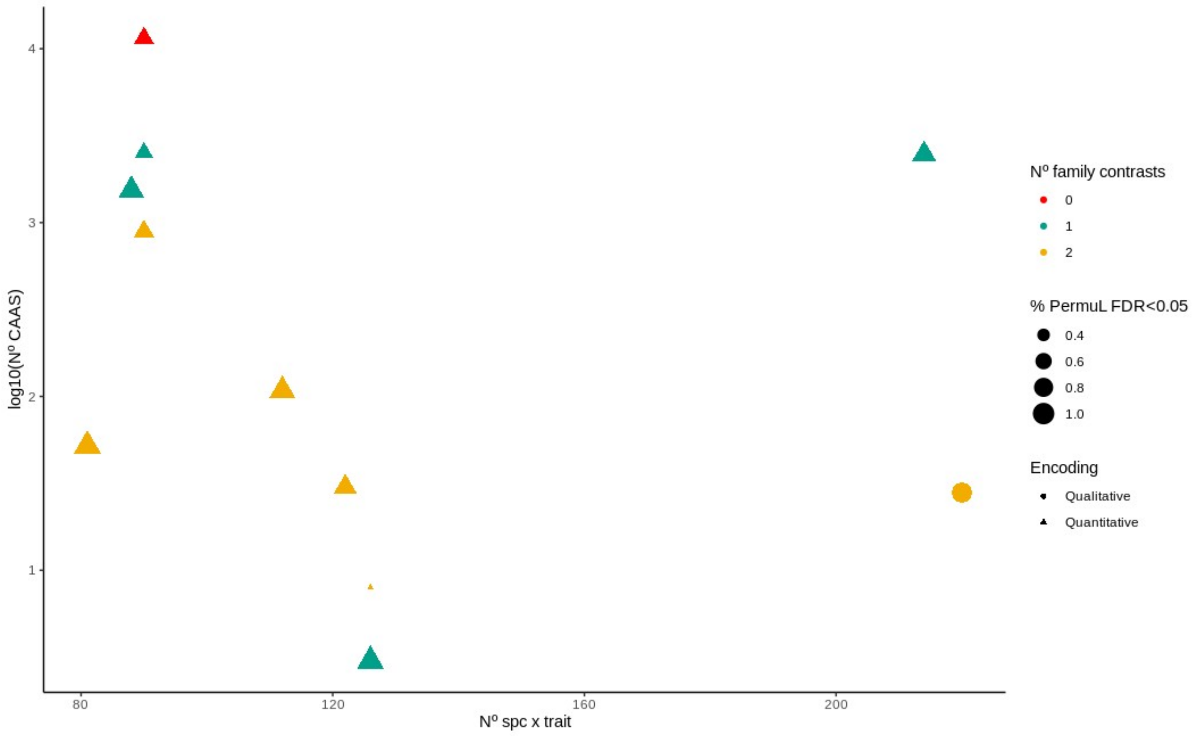


**Fig. S39.**

Descriptive CAAS plot between the total number of CAAS positions recovered per trait (y) and the number of species tested per trait (x), accounting for the proportion of positions (used in the external mammalian analysis) with permulations FDR p-value <0.05 (per trait basis) and classifying them according to A) encoding type (shape) and B) number of *family contrasts* (color).

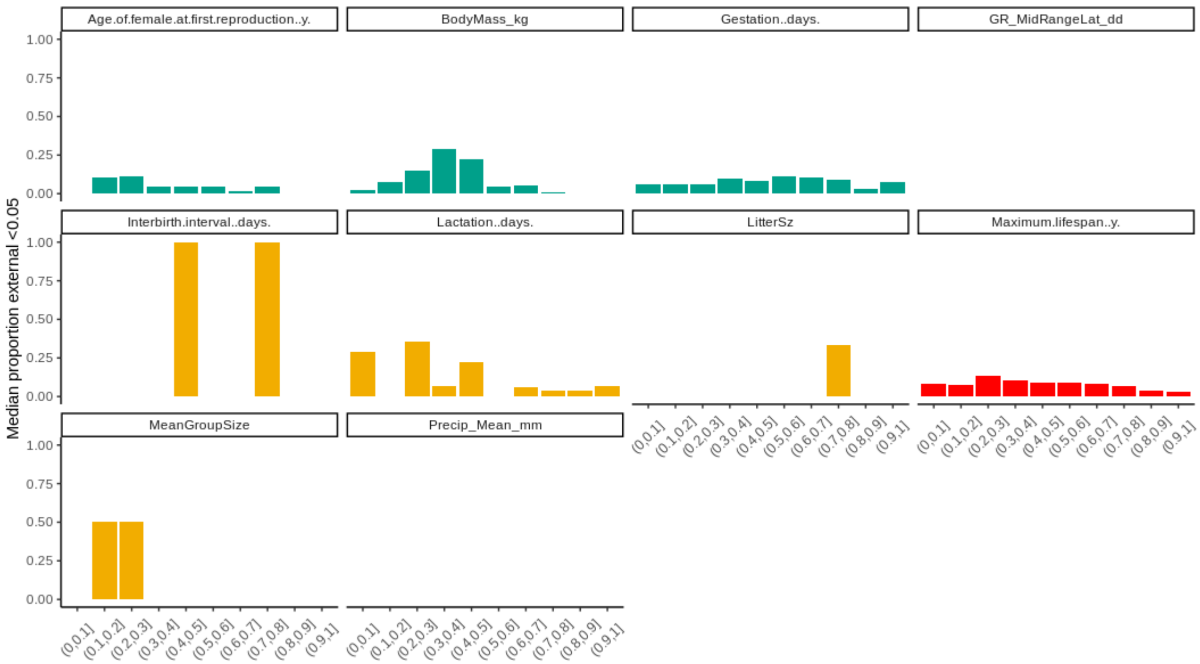


#### Fig. S40.

Descriptive CAAS plot between the median proportion of positions with nominal external p-value<0.05 per trait (y) and deciles of amino acid position conservation (x), accounting for quantitative traits (used in the external mammalian analysis) and classifying them according to A) number of *family contrasts* (red=0, turquoise=1, gold=2).

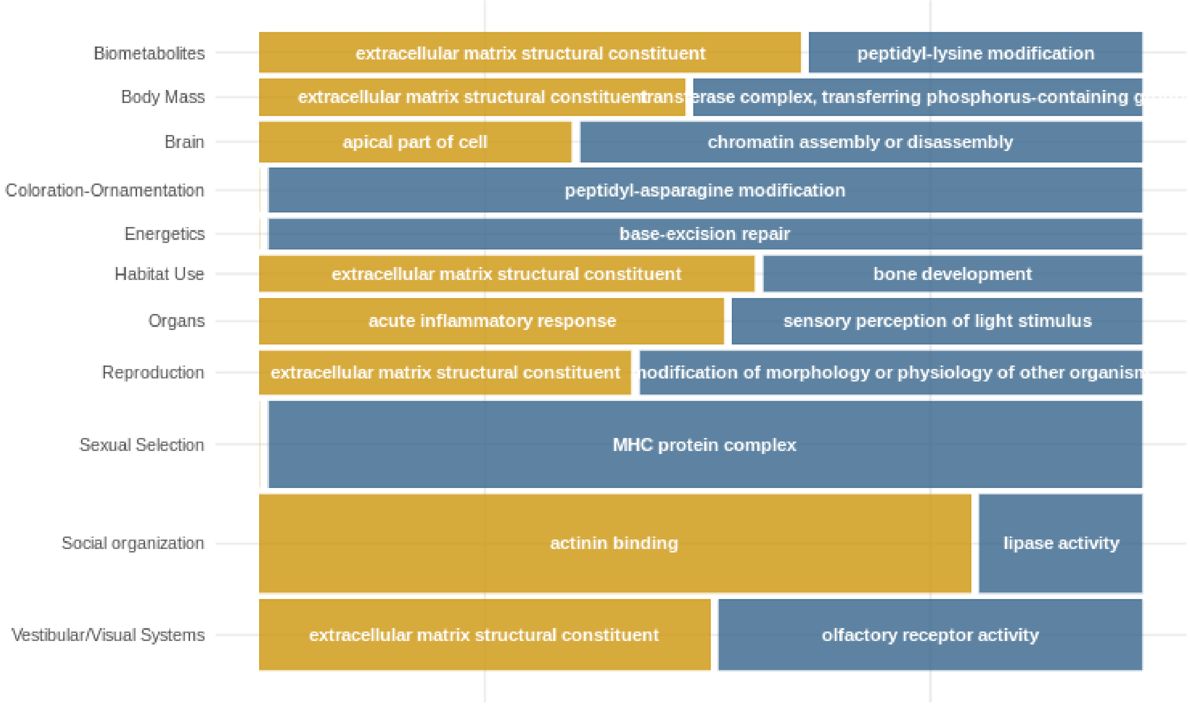


#### Fig. S41.

Mosaic plots with the most common functional descriptive term (FDR<0.05) for perm p-val<0.05 gene lists for GO non-redundant ontologies (Biological Process, Cellular Component, and Molecular Function) in the *PPGPM*. The square areas are proportional to the median enrichment ratio of the respective terms.

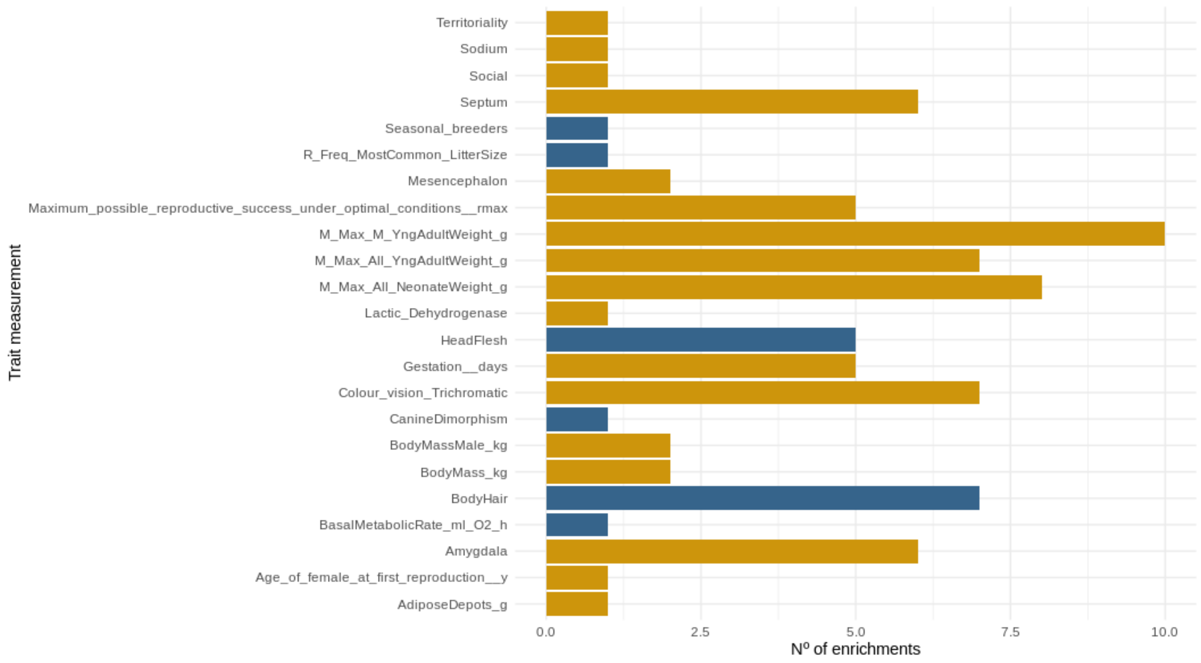


**Fig. S42.**

Number of functional enrichments for KEGG pathways database with FDR<0.05 for both methodologies (CAAS and RERs) per trait in the *PPGPM*

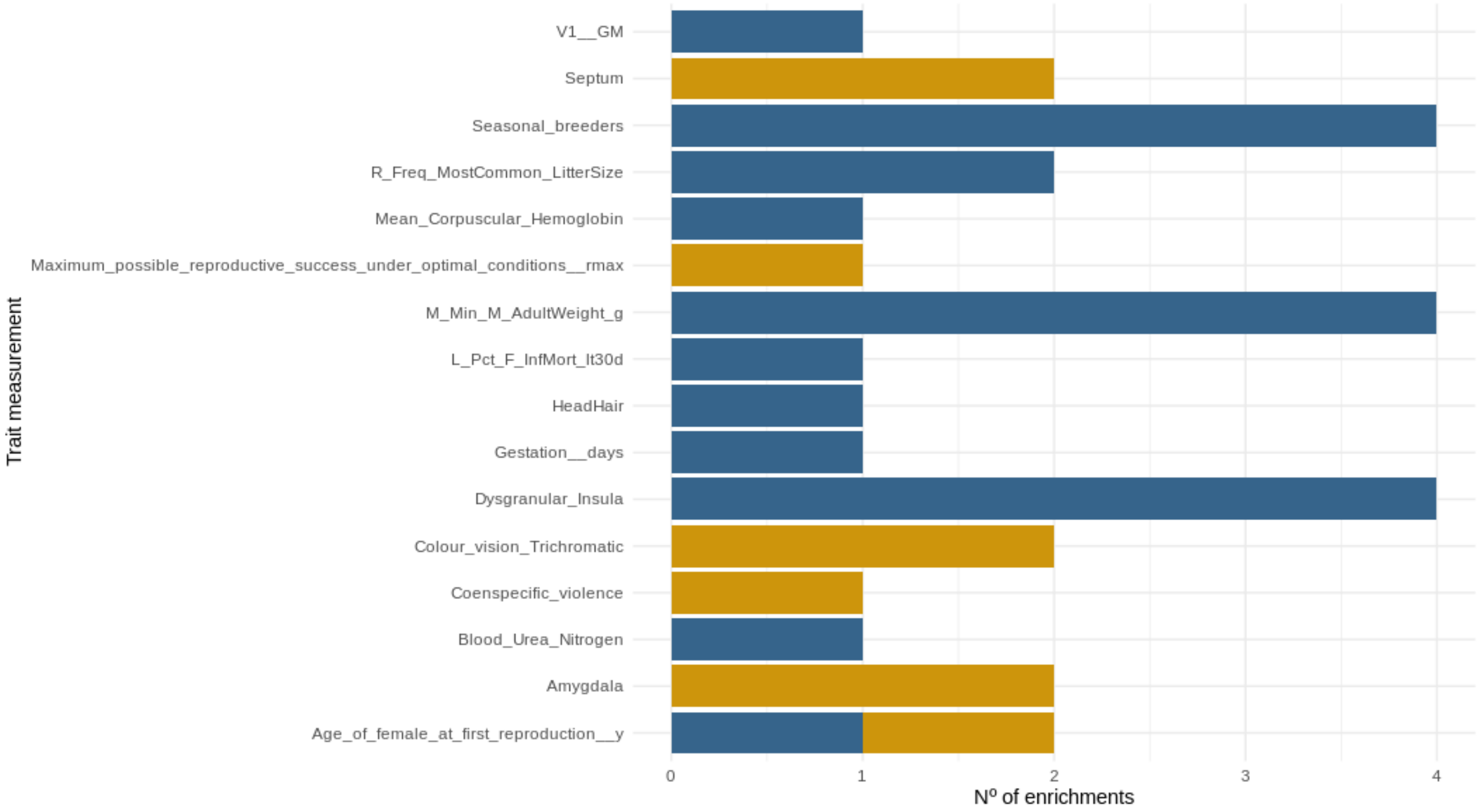


**Fig. S43.**

Number of functional enrichments for Panther pathways database with FDR<0.05 for both methodologies (CAAS and RERs) per trait in the *PPGPM*

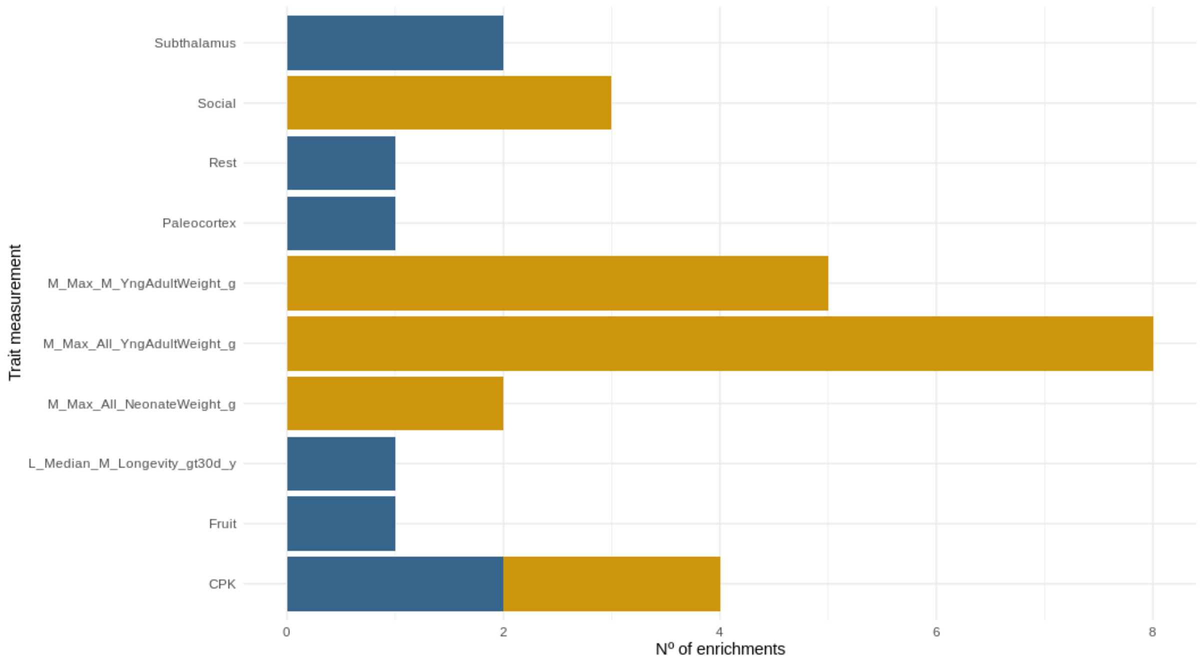


**Fig. S44.**

Number of functional enrichments for the human phenotype ontology (HPO) database with FDR<0.05 for both methodologies (CAAS and RERs) per trait in the *PPGPM*

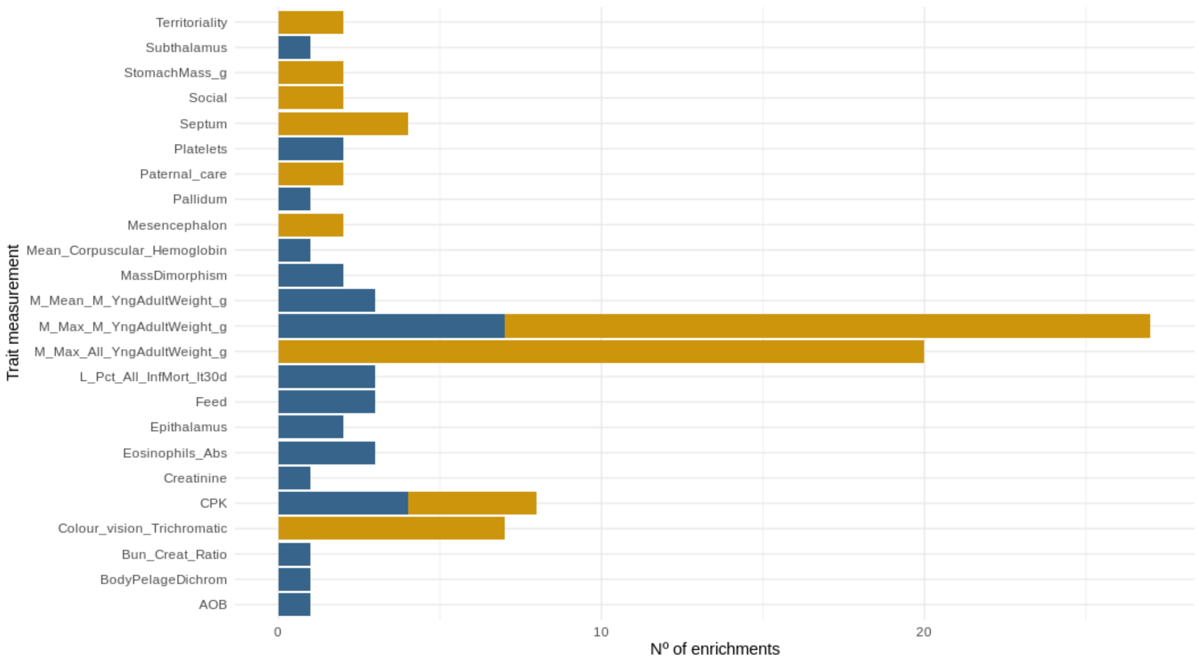


**Fig. S45.**

Number of functional enrichments for the DisGeNET disease database with FDR<0.05 for both methodologies (CAAS and RERs) per trait in the *PPGPM*

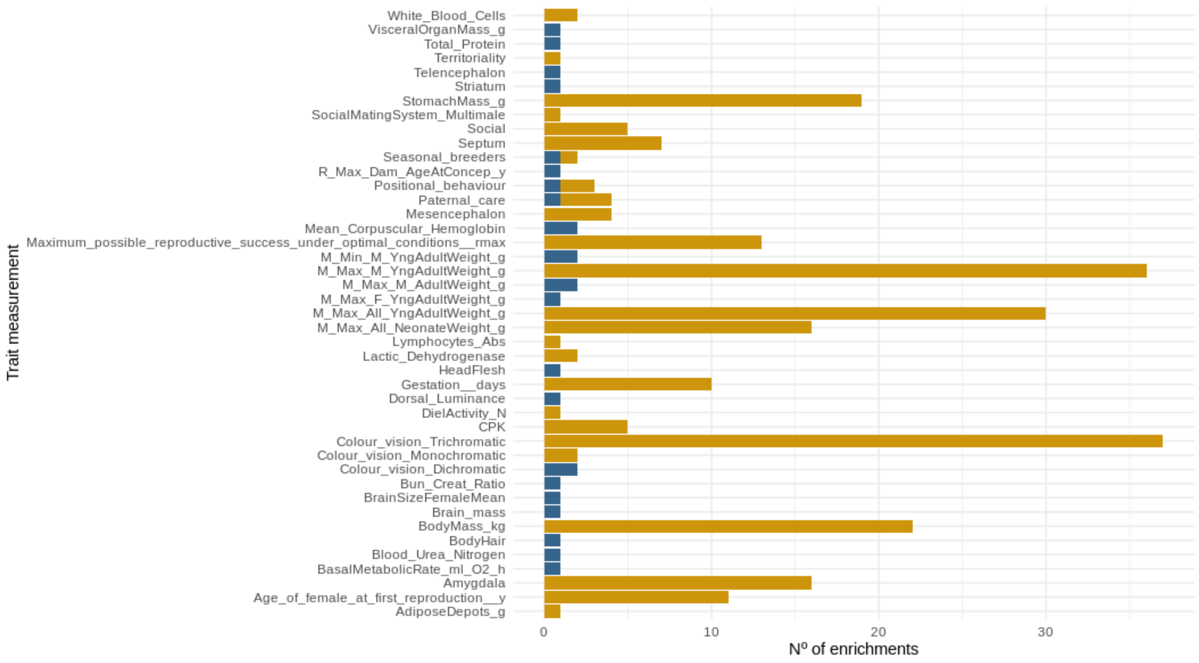


#### Fig. S46.

Number of functional enrichments for *GO ontologies* (Biological Process, Cellular Component, and Molecular Function) database with FDR<0.05 for both methodologies (CAAS and RERs) per trait in the *PPGPM*

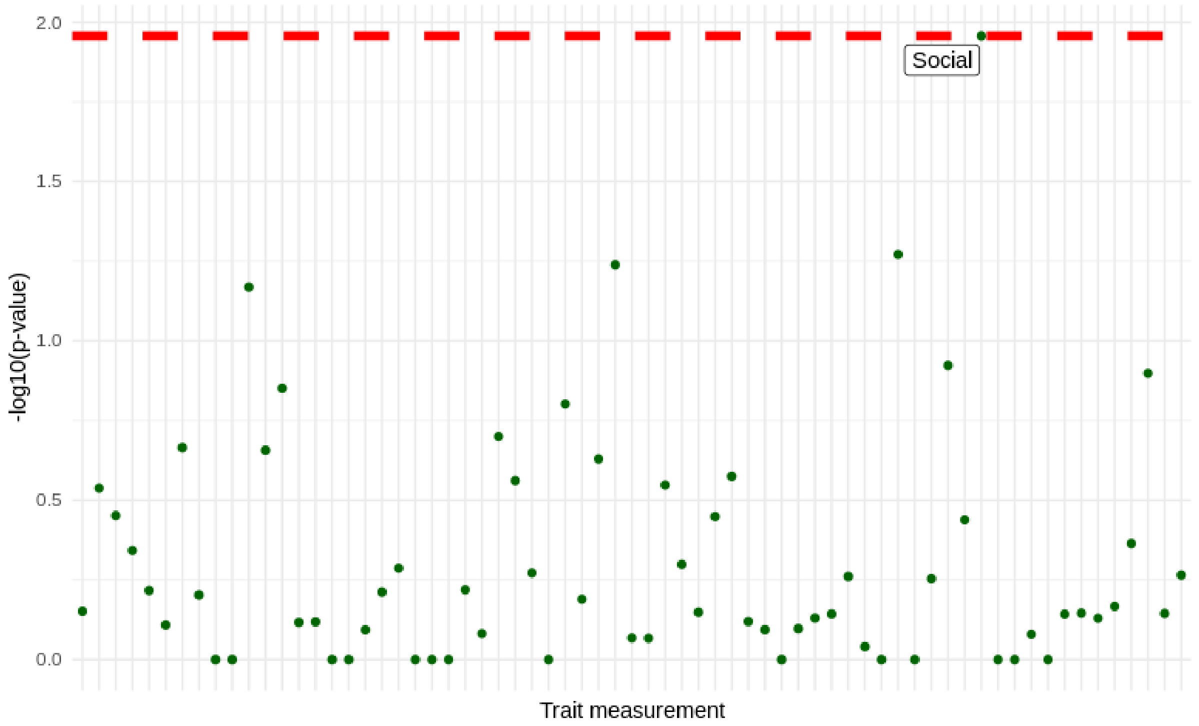


#### Fig. S47.

Gene sets overlap for each trait with within-family contrasting phenotypes between the CAAS and RERS methodologies associated with gene sets with permulations <0.05 in the PPGPM. The dashed red line indicates FDR<0.05 for the hypergeometric test of gene set overlap.

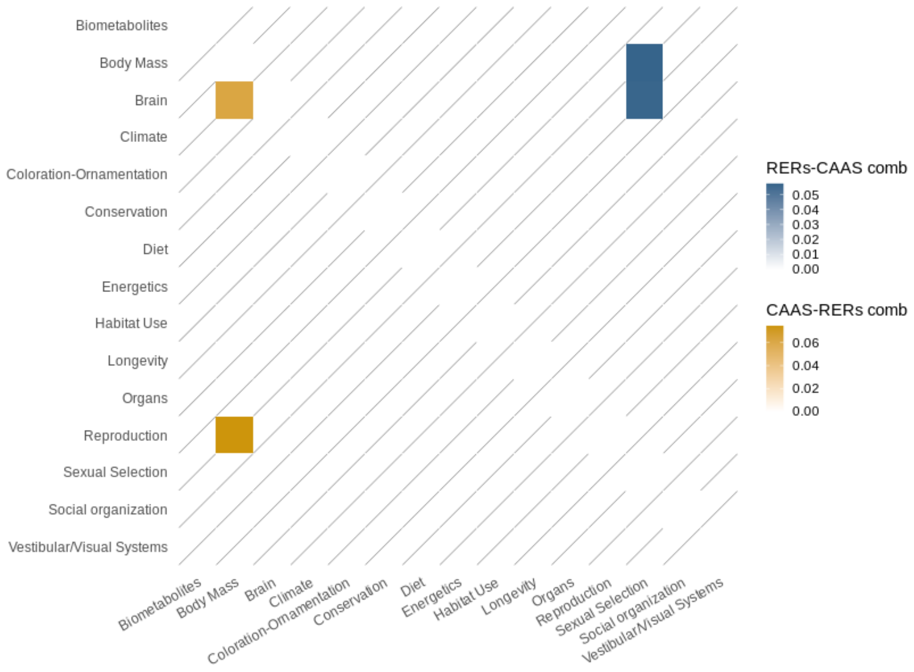


#### Fig. S48.

Heatmap representing the median proportion of overlapping genes between each CAAS trait gene set against other RER trait gene sets (gold) and each RERs trait gene set against other CAAS trait gene sets (blue) from different biological secondary domains with an FDR<0.05 from the hypergeometric test of gene set overlap. Each color corresponds to a PCM method.

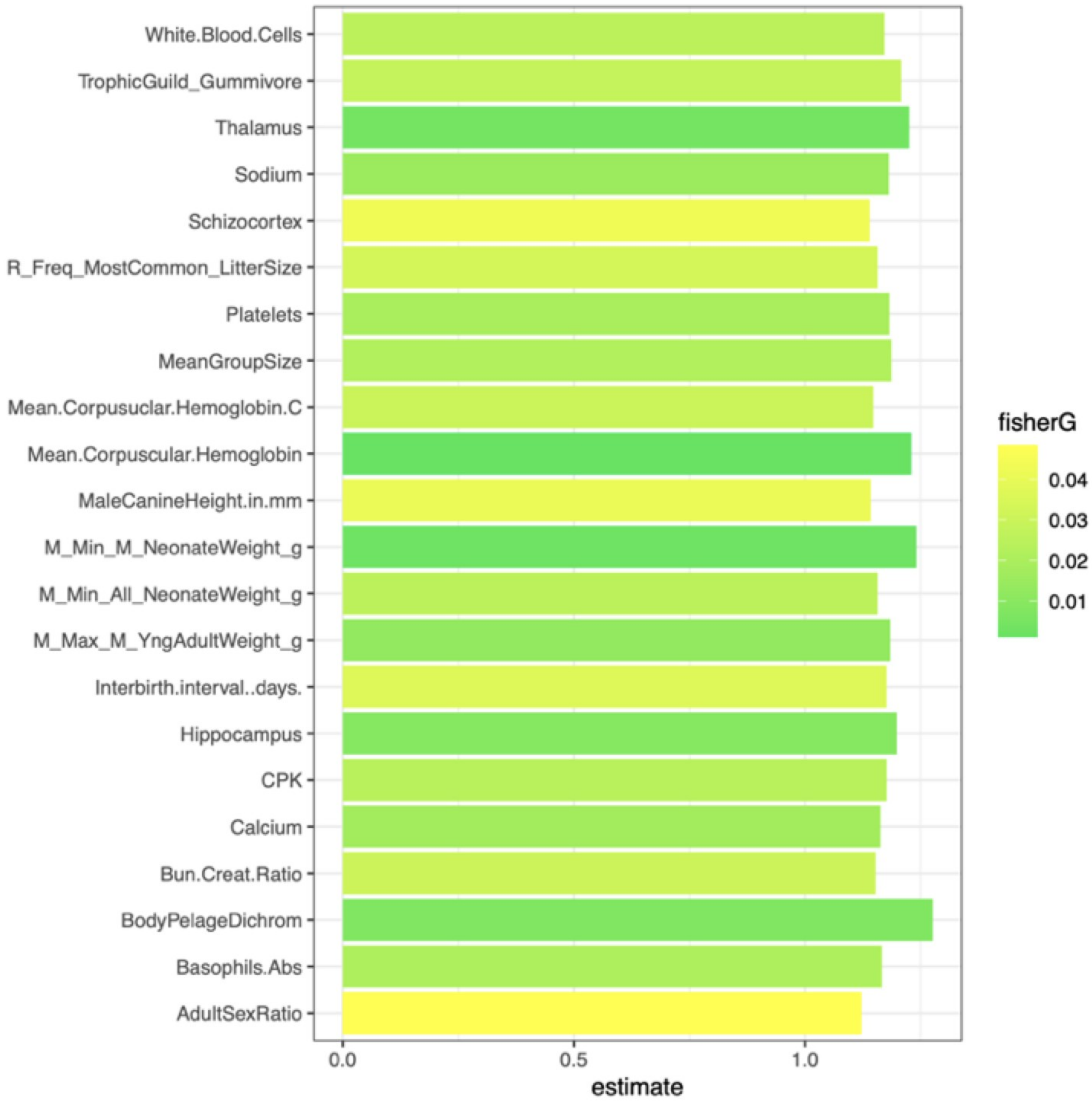


**Fig. S49.**

Results of high-impact mutation test with OMIM data for RERs gene lists, where 22 significant traits have a nominal p-value < 0.05

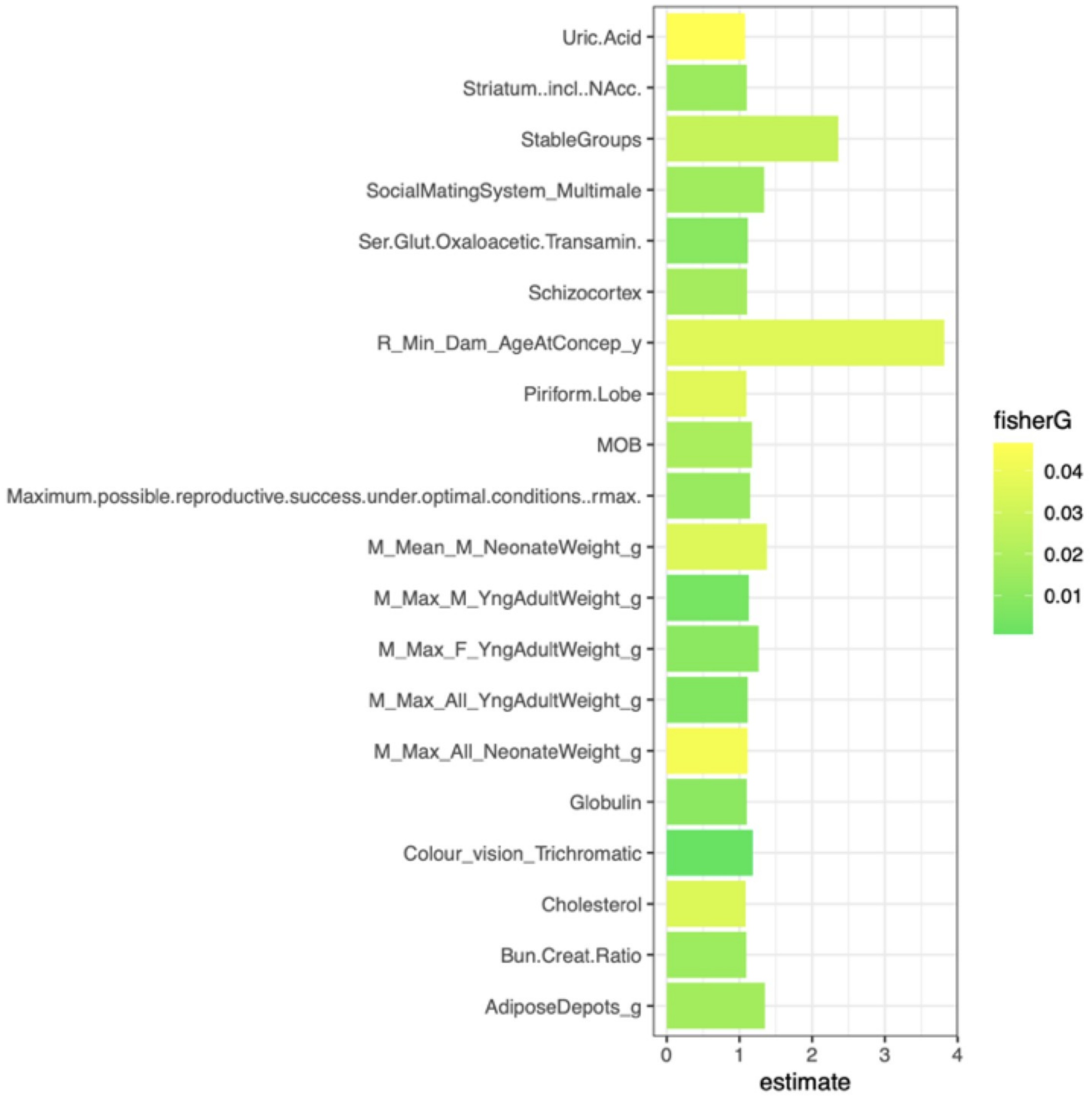


#### Fig. S50.

Results of high-impact mutation test with OMIM data for CAAS gene lists, where 23 significant traits have a nominal p-value < 0.05

#### Supplementary Tables

| Biological trait | Trait (original source nomenclature) | n°_families | n°_species |
|---|---|---|---|
| Body mass | BodyMass_kg | 16 | 214 |
| | Body_mass | 16 | 139 |
| | Body.mass..g. | 15 | 98 |
| | Body.Weight | 10 | 37 |
| | BodyMassForOrganMass_g | 6 | 19 |
| | M_Mean_All_AdultWeight_g | 6 | 24 |
| Brain size | Brain_mass | 16 | 139 |
| | BrainSizeSpeciesMean | 16 | 128 |
| | BV | 16 | 63 |
| | Endocranial.volume..cm3. | 15 | 97 |
| | BrainMass_g | 6 | 19 |
| Maximum longevity | Maximum.lifespan..y. | 15 | 130 |
| | MaxLongevity_m | 15 | 115 |
| | L_Max_All_Age_y | 6 | 24 |
| Interbirth interval period | Interbirth.interval..days. | 15 | 94 |
| | InterbirthInterval_days | 15 | 93 |
| Litter size | LitterSz | 16 | 122 |
| | Offspring.per.litter | 15 | 76 |
| | R_Mean_LitterSize | 6 | 24 |
| Gestation period length | Gestation..days. | 16 | 113 |
| | Gestation_length | 16 | 110 |
| | Gestation | 16 | 102 |
| | R_Expected_Gestation_d | 6 | 24 |

| | | | |
|---|---|---|---|
| Lactation period length | Lactation..days. | 16 | 103 |
| | WeaningAge_d | 16 | 95 |
| | Weaning_age | 16 | 93 |
| Group size | MeanGroupSize | 15 | 112 |
| | SocialGroupSize | 11 | 80 |
| Canine dimorphism | CanineDimorphism | 14 | 93 |
| | SexualCanineDimorphism | 11 | 80 |
| Mass dimorphism | MassDimorphism | 16 | 143 |
| | SexualSizeDimorphism | 11 | 80 |
| Body mass in males | BodyMassMale_kg | 16 | 197 |
| | Body.MassMaleMean | 16 | 142 |
| | meanMaleBodyMass.in.g | 11 | 80 |
| | MaleBodyMass.CanineDataset..in.g | 11 | 80 |
| | MaleBodyMass.TestesDataset..in.g | 11 | 55 |
| | MaleBodyMass.SexSzeDimorphismDataset..in.g | 11 | 79 |
| | M_Mean_M_AdultWeight_g | 6 | 24 |
| Body mass in females | BodyMassFemale_kg | 16 | 193 |
| | BodyMassFemaleMean | 16 | 142 |
| | FemaleBodyMass.SexSizeDimorphismDataset..in.g | 11 | 79 |
| | M_Mean_F_AdultWeight_g | 6 | 24 |
| Social system | Social_organization | 16 | 191 |
| | SocialSystem | 16 | 149 |
| Mating_system | Mating_system | 16 | 140 |
| | Mating.System | 16 | 110 |
| N° males per group | AdultMales | 15 | 108 |

| | | | |
|---|---|---|---|
| | MalesPerBreedingGroup | 11 | 45 |
| N° females per group | AdultFemale | 15 | 108 |
| | FemalesPerMaleInBreedingGroup | 9 | 28 |
| Neonate mass | Neonatal.mass..g. | 15 | 76 |
| | M_Mean_All_NeonateWeight_g | 6 | 23 |
| Seasonal breeding | Seasonal_breeders | 16 | 114 |
| | R_Pattern_Breeding | 6 | 24 |
| Actual evapotranspiration | AET_Mean_mm | 16 | 126 |
| | Actual.evapotranspiraton..mm. | 13 | 80 |

**Table S3.**

A set of 19 phenotypes with several traits was collected from different sources of primate phenotypic data. The selection of traits was based on both the number of species and the number of families covered by our protein-coding alignments.

| **Gene name** | **Trait** | **Rho** | **N branches** | **RERconverge p-value** | **Permulations p-value** |
|---|---|---|---|---|---|
| *ACTR8* | Segmented neutrophil absolute count | 0.6549591 | 42 | 2.525129e-06 | <0.001 |
| *ARL3* | Body mass in males | 0.2772338 | 338 | 2.220248e-07 | 0.004 |
| *CFL2* | Platelets counts | 0.8190697 | 24 | 9.884246e-07 | <0.001 |
| *CIITA* | Diencephalon relative size | -0.7248529 | 38 | 2.657240e-07 | <0.001 |
| *DEFA5* | % Fauna in diet | -0.4978560 | 89 | 6.903441e-07 | <0.001 |
| *DESI2* | Mesencephalon relative size | 0.6858511 | 42 | 5.371317e-07 | <0.001 |
| *EID2B* | Visual acuity | -0.9222696 | 12 | 1.959267e-05 | 0.014 |
| *F8* | Body mass in males | 0.2627143 | 295 | 4.791844e-06 | 0.010 |
| *FBXO34* | Dysgranular Insular cortex relative size | -0.8423690 | 24 | 2.434203e-07 | <0.001 |
| *GAL3ST1* | Brain mass | -0.3159678 | 214 | 2.399007e-06 | <0.001 |
| *HYPK* | Triglycerides levels | 0.6998466 | 37 | 1.432104e-06 | <0.001 |
| *MYL4* | Visual acuity | 0.9617807 | 10 | 8.913579e-06 | 0.002 |
| *NKX6-2* | % Time moving | 0.6168111 | 52 | 1.120037e-06 | <0.001 |
| *PTPN20* | Platelets count | 0.9528742 | 12 | 1.690916e-06 | <0.001 |
| *RARB* | Visual acuity | 0.9658973 | 10 | 5.678678e-06 | 0.002 |
| *SFTA3* | Ventral coloration | 0.5338828 | 69 | 2.311368e-06 | <0.001 |
| *SHH* | Body mass in males | 0.2558341 | 341 | 1.694375e-06 | 0.005 |
| *SOD1* | Lung relative mass | -0.9241871 | 14 | 2.324497e-06 | <0.001 |
| *SPA17* | Brain mass | -0.3193915 | 204 | 3.218927e-06 | <0.001 |
| *SSU72* | Chloride levels | -0.6875768 | 40 | 9.519273e-07 | <0.001 |
| *TAF13* | Eosinophils proportion | -0.6991929 | 38 | 1.043451e-06 | <0.001 |
| *TNNI3* | Visual acuity | 0.9764933 | 10 | 1.298493e-06 | 0.002 |
| *ZBTB46* | Adipose depots mass | -0.9476824 | 12 | 2.826437e-06 | <0.001 |
| *ZNF599* | Maximum young adult body weight in females | -0.7385527 | 30 | 3.167689e-06 | <0.001 |

**Table S4.**

Set of 24 associations between relative evolutionary rates (RERs) and quantitative traits with parametric FDR <0.05 per trait

| Trait | Nº family contrasts | Nº positions | Gene Names (Top nº: 30) |
|---|---|---|---|
| Body mass | 1 | 23 | *ADAMTSL1*, *ADCY7*, *CENPN*, *DBH*, *ICAM1*, *ICAM4*, *KIAA1549*, *LCN9*, *LIMCH*, *NDUFS7*, *NOXO1*, *OR8S1*, *RFPL4A*, *ST18*, *TPGS2*, *ZNF729*, *ZNF786* |
| Body mass in females | 2 | 2 | *CELA3B*, *ITGAX* |
| Creatine phosphokinase levels | 2 | 31 | *ADGRG5*, *ALDH1L1*, *ALMS1*, *ARHGEF28*, *BMP3*, *C16orf71*, *CARS*, *CENPU*, *CLRN3*, *COL6A6*, *CRB1*, *DEGS1*, *GCC1*, *HSD17B11*, *ICOS*, *JADE2*, *KRT74*, *KRTAP1-1*, *LAMC3*, *LILRB3*, *NAAA*, *OAS3*, *RAB11FIP1*, *RARS*, *RITA1*, *SDE2*, *SPATA13*, *SUPT3H*, *TAS1R2*, *TTC3* |
| Lactation period length | 2 | 4 | *ADGRV1*, *KIAA0586*, *MEFV*, *PUS10* |
| Lactic dehydrogenase levels | 1 | 1,518 | *ADAMTS16*, *ALPK3*, *ARHGAP23*, *CCDC73*, *CDCA2*, *CDK18*, *CYSLTR2*, *DLEC1*, *DISC1*, *DMRTB1*, *ELL3*, *EPB42*, *FAM184B*, *FBXO30*, *HADHA*, *ICAM4*, *KIAA1549*, *KIAA1755*, *LY75-CD302*, *MAMDC4*, *METTL4*, *MIER1*, *N4BP2*, *NCAPD3*, *OSMR*, *SETD4*, *SRBD1*, *TRIM66*, *UGT2B4* |
| Mesencephalon relative size | 1 | 7 | *CASP8*, *CYB5D2*, *LRRC34*, *PLAUR*, *TDRD1*, *TONSL*, *XIRP1* |
| Nº female adults per social | 2 | 5 | *ERLEC1*, *MTMR1*, *MUC13*, *TRMO*, *ZXDC* |

| | | | |
|---|---|---|---|
| group | | | |
| Neutrophils band proportion | 2 | 4 | *NPRL3*, *PTGDS*, *RTTN*, *TMEM59* |
| Neutrophils band absolute count | 2 | 4 | *NPRL3*, *PTGDS*, *RTTN*, *TMEM59* |
| Septum relative size | 2 | 261 | *AMOTL1*, *CA5B*, *CAMSAP1*, *CD276*, *CEP104*, *CNKSR1*, *CSF1R*, *CYLC1*, *DNAH1*, *EHBP1*, *GAS2L1*, *GFY*, *GOLGA4*, *IL1RL2*, *KAT6A*, *LIMS2*, *LRRN4*, *MAP3K19*, *MFSD2A*, *NLRP11*, *PEX5*, *PKD1L2*, *PTCH1*, *SBK3*, *SERHL2*, *SETX*, *SIAH3*, *TAPBPL*, *ZADH2*, *ZNF25* |
| % Time socializing | 2 | 9 | *BAHCC1*, *CD226*, *DNAJB2*, *EEF1D*, *LRRTM4*, *SLC10A6*, *TMEM18*, *UBR1*, *UCN3* |
| Trichromatic color vision | 1 | 141 | *ACIN1*, *AKAP8L*, *AMZ2*, *ARHGEF5*, *ATP13A4*, *ATP2A1*, *B3GNT5*, *CCDC105*, *DSG2*, *DUSP5*, *E2F8*, *FBXO39*, *GAS2L2*, *GRAMD4*, *GSTZ1*, *HECW1*, *HECW2*, *ICE1*, *ILF3*, *ITGA2B*, *KIAA0319L*, *LIMA1*, *LRGUK*, *MEI4*, *MGST2*, *MIS18A*, *NCKAP5*, *PHF3*, *POLD3*, *SLC17A5* |
| White blood cells count | 2 | 43 | *APOBEC3F*, *BAG6*, *BDP1*, *CACNA1A*, *CDKAL1*, *CYP7B1*, *DPP9*, *FAF1*, *FGD5*, *GGCX*, *GOLIM4*, *HSPE1*, *IZUMO1R*, *MATN2*, *MYO3B*, *MYO5C*, *NHS*, *RASIP1*, *SLC9A5*, *THY1*, *ZNF429*, *ZNF644* |

**Table S5.**

A set of 13 quantitative and qualitative traits with $p_{PGLS\text{-}FDR}< 0.05$ and $p_{phyloglm\text{-}FDR}< 0.05$, respectively, obtained from CAAS primate consistency tests, only considering >5 species in both groups and traits with at least one family contrast.

| Gene name | Trait | Position in primate alignment | Nº Top AA | Nº Bottom AA | PGLS p-value |
|---|---|---|---|---|---|
| *AGBL3* | Maximum lifespan | 13 | 15 | 76 | 2.330802e-12 |
| *AKAP11* | Maximum lifespan | 1474 | 6 | 69 | 2.265749e-05 |
| *C1orf127* | Body mass in kg | 355 | 23 | 14 | 2.253048e-04 |
| *CRAMP1* | Maximum lifespan | 479 | 6 | 23 | 1.949479e-04 |
| CX3CL1 | Maximum lifespan | 36 | 10 | 52 | 5.456237e-05 |
| FAM120B | Maximum lifespan | 224 | 8 | 30 | 7.322584e-05 |
| FAM83G | Maximum lifespan | 765 | 11 | 83 | 1.427407e-04 |
| GPA33 | Maximum lifespan | 261 | 8 | 51 | 1.273069e-04 |
| IL1R1 | Maximum lifespan | 307 | 9 | 89 | 1.887379e-15 |
| KCNH4 | Maximum lifespan | 324 | 14 | 94 | 1.835312e-09 |
| KIAA1328 | Maximum lifespan | 465 | 18 | 84 | 5.460077e-13 |
| LAPTM5 | Maximum lifespan | 54 | 37 | 57 | 3.596821E-04 |
| MAP2 | Maximum lifespan | 223 | 22 | 68 | 6.036033e-06 |
| MFSD13A | Maximum lifespan | 233 | 18 | 86 | 1.452949e-12 |
| MPDZ | Gestation period length | 948 | 40 | 72 | 1.647184e-05 |
| MPEG1 | Maximum lifespan | 670 | 22 | 63 | 1.102206e-04 |
| NELL1 | Lactation period length | 89 | 7 | 91 | 1.089895e-04 |
| NRIP2 | Body mass in kg | 114 | 21 | 95 | 4.020017e-04 |
| OR2S2 | Interbirth interval period length | 90 | 15 | 22 | 1.370469e-04 |
| ORC1 | Maximum lifespan | 647 | 10 | 96 | 6.181353e-06 |
| PPP4R1 | Maximum lifespan | 382 | 22 | 70 | 2.433866e-04 |
| PRR35 | Maximum lifespan | 254 | 21 | 78 | 1.284168e-05 |
| PTPN22 | Body mass in kg | 51 | 23 | 103 | 3.345659e-04 |
| R3HCC1L | Maximum lifespan | 418 | 6 | 80 | 2.962214e-05 |
| SMARCAL1 | Body mass in kg | 332 | 27 | 109 | 3.920861e-04 |
| SPINK5 | Maximum lifespan | 727 | 13 | 36 | 1.357845e-04 |

| | | | | | |
|---|---|---|---|---|---|
| SRBD1 | Maximum lifespan | 445 | 8 | 86 | 3.551777e-05 |
| TCF7L2 | Maximum lifespan | 420 | 16 | 86 | 2.448851e-08 |
| TIE1 | Maximum lifespan | 57 | 10 | 68 | 1.612757e-04 |
| TMEM65 | Maximum lifespan | 93 | 35 | 50 | 2.764147e-04 |
| UCP1 | Maximum lifespan | 248 | 6 | 43 | 1.446069e-05 |
| YBX2 | Maximum lifespan | 313 | 25 | 76 | 1.421607e-04 |
| ZNF707 | Lactation period length | 93 | 9 | 10 | 3.431550e-04 |

**Table S6.**

Set of 33 significant amino acid trait associations retrieved from the cross-mammalian consistency test in the external mammalian dataset (TOGA) after PGLS analysis with FDR correction per trait (8 with intra-family contrasts), only considering >5 species in both groups

| geneSet | Description | Size | Overlap | Enrichment Ratio | P-value | FDR | GO category | Trait |
|---|---|---|---|---|---|---|---|---|
| GO:0006302 | double-strand | 150 | 30 | 2.12612021857 | 5.162232272815093e | 0.005484871 | BP | Body mass in kg |

| | | | | | | | | |
|---|---|---|---|---|---|---|---|---|
| | break repair | | | 9235 | -5 | 789866036 | | |
| GO:0036297 | interstrand cross-link repair | 32 | 10 | 3.3220628415300544 | 4.8564817956875395e-4 | 0.03175391943334 1604 | BP | Body mass in kg |
| GO:0000726 | non-recombinational repair | 42 | 22 | 4.053400523457948 | 9.198575234847794e-10 | 7.791193223916082e-7 | BP | Maximum neonate body weight |
| GO:0006302 | double-strand break repair | 139 | 44 | 2.4495370069817812 | 5.703840733062293e-9 | 2.415576550451881e-6 | BP | Maximum neonate body weight |
| GO:1990391 | DNA repair complex | 25 | 11 | 3.4295470383275264 | 1.1692516226569083e-4 | 0.0138768215152 19406 | CC | Maximum neonate body weight |
| GO:0006310 | DNA recombination | 198 | 36 | 1.932836562344759 | 8.026332975319406e-5 | 0.0075804255878 01661 | BP | Body mass in kg |
| GO:0006310 | DNA recombination | 178 | 46 | 1.9997880008479967 | 2.0615282954050684e-6 | 4.365286165520232 4e-4 | BP | Maximum neonate body weight |
| GO:0032200 | telomere organization | 100 | 26 | 2.011960623461854 | 2.995323125113414e-4 | 0.0362434098138 7231 | BP | Maximum neonate body weight |
| GO:0006260 | DNA replication | 223 | 81 | 1.4208340152531833 | 2.129686473086867e-4 | 0.0362046700424 7674 | BP | Maximum young adult body weight |
| GO:0006260 | DNA replication | 223 | 74 | 1.483171701380448 | 1.1873181320032344e-4 | 0.0252305103050 68732 | BP | Maximum young adult body weight in males |
| GO:0004518 | nuclease | 165 | 52 | 1.43822 | 0.0024834 | 0.0403 | MF | Maximum young adult body weight |

| | | | | | | | | |
|---|---|---|---|---|---|---|---|---|
| | activity | | | 539116 65674 | 225824409 417 | 19096 04433 5284 | | in males |
| GO:0140097 | catalytic activity, acting on DNA | 157 | 56 | 1.42002 458375 23744 | 0.0019475 656277141 429 | 0.0408 98894 49671 458 | MF | Maximum young adult body weight in males |
| GO:0140097 | catalytic activity, acting on DNA | 157 | 52 | 1.51151 076141 70933 | 7.2039914 05548837e -4 | 0.0281 60477 01491 3237 | MF | Maximum young adult body weight in males |

**Table S7.**

Set of enriched GO terms linked to DNA repair/replication/recombination categories for *body-mass related traits* with contrasting families, showing significant hypergeometric test (pFDR<0.05)

**Data S1 (separate file). Download:**
https://docs.google.com/spreadsheets/d/1ciEoWHDYMQaGiHi2zAQXY-ASgu_LIou8S_BLglnwYh8/edit?usp=sharing

**Spreadsheet with phenomic data and metadata analysis**

**Data S2 (separate file). Download:**

https://docs.google.com/spreadsheets/d/1dwhFFPoGPVGi9rAc_5dYKCdCXA2QaNdXpKZLYZ8Y7Z0/edit?usp=sharing

**Spreadsheet with gene alignments metadata and CAAS/RERconverge lists of positions and genes**

**Data S3 (separate file)**

https://docs.google.com/spreadsheets/d/1IaXdtR02TVpIZJi5wtZeS6bHLz30NOCCm3zqMYjrtyg/edit?usp=sharing

**Spreadsheet with functional enrichment and between-domains overlap analyses**

### Technical Appendix

#### Software versions, command-line parameters, and reproducibility resources

Detailed information on the software versions, command-line parameters, and scripts used in the analyses described in the Supplementary Methods are provided below. All scripts and additional reproducibility resources are publicly available in the official project repository.

The complete set of scripts used to generate the figures, process genomic data, and run the genome–phenome analyses is available at [https://github.com/pgarchive/data](https://github.com/pgarchive/data). Within this repository, the folder supplementary_info/ contains the following:

- scripts used to generate the manuscript figures
- documentation of the software versions used in the analyses
- configuration files for the CAAS analyses
- additional metadata required to reproduce the analyses

#### TA1. Software used for genomic and phylogenetic analyses

The following software packages were used throughout the study for orthology detection, sequence alignment, phylogenetic analysis, and comparative genomic analysis.

| Software | Version | Purpose |
|---|---|---|
| BLAST+ | v2.12.0 | Orthology detection using best-bidirectional hits |
| MUSCLE | v3.8.31 | Multiple sequence alignment of protein sequences |
| trimAL | v1.2 | Alignment filtering |
| MACSE | v2.04 | Codon-aware alignment of coding sequences |
| BMGE | v1.1 | Filtering of poorly aligned codon positions |
| newick-tools | v1.5 | Pruning of phylogenetic trees |
| PAML (codeml) | v4.9 | Estimation of branch lengths in gene trees |
| HyPhy | v2.5.2 | Detection of directional selection (FADE) |
| RERconverge | v0.1.0 | Relative evolutionary rate analyses |
| webGestaltR | v0.4.4 | Functional enrichment analyses |

Additional analyses were performed in **R** using standard phylogenetic and statistical packages, including:

- phytools

- caper
- factorMineR
- Rphylip
- Phylolm
- biomaRt

### TA2. Orthology detection

Orthologous protein-coding genes were identified using BLASTP best-bidirectional-hit (BBH) searches across a set of primate reference genome seqeunces. For each human gene model, orthology clusters were constructed by iteratively identifying reciprocal best hits between the reference species and remaining primate species.

Example command:

```
blastp -query query.fasta -db reference_db -outfmt 6
```

The resulting orthology clusters were subsequently filtered to retain those with orthologous sequences present in at least 40 of the reference species.

### TA3. Multiple sequence alignments

Protein and coding sequence alignments were generated using a combination of MUSCLE, MACSE, and BMGE following a codon-aware alignment strategy.

#### Protein alignments

Initial protein alignments were generated using the following methods:

```
muscle -in input.fasta -out aligned.fasta
```

Alignment filtering was subsequently performed using **trimAl** as follows:

```
trimal -in aligned.fasta -out filtered.fasta
```

#### Codon-aware alignment of CDS

The coding sequences were aligned using MACSE, preserving the codon structure and allowing the detection of frameshifts and stop codons. Initial alignment of the reference sequences:

```
java -jar macse_v2.04.jar -prog alignSequences \
-seq reference_sequences.fasta \
-out_NT aligned_references.fasta
```

Incorporation of resequenced species:

```
java -jar macse_v2.04.jar -prog enrichAlignment \
-align aligned_references.fasta \
-seq resequenced_sequences.fasta \
-new_seq_alterable_ON \
-out_NT resequenced_aligned.fasta
```

Exporting filtered alignments:

```
java -jar macse_v2.04.jar -prog exportAlignment \
-align resequenced_aligned.fasta
```

#### TA4. Alignment filtering

Codon alignments were filtered using BMGE, which operates at the codon level to remove poorly aligned sequence positions.

Example command:

```
java -jar BMGE.jar -i input_alignment.fasta -t CODON
```

Positions containing more than **10% of gaps** were excluded.

#### TA5. Phylogenetic tree processing

Species trees were pruned to match the set of species present in each gene alignment using newick-tools.

Example command:

```
nw_prune species_tree.nw species_list.txt > pruned_tree.nw
```

Branch lengths for gene trees were estimated using **codeml** from the **PAML** software package.

#### TA6. Detection of convergent amino acid substitutions (CAAS).

CAAS analyses were performed using CAAStools (Barteri et al., 2023), following the methodological framework described in the Methods section.

The exact version of the software and configuration parameters used in the analyses are documented in the repository:

`https://github.com/pgarchive/data`

Specifically, the directory is as follows:

`supplementary_info/`

and it contains:

- the version of CAAStools used
- configuration files for CAAS analyses
- scripts used to perform trait binarization and species selection
- additional documentation describing the analysis workflow.

#### TA7. Figure generation

All figures presented in the manuscript and supplementary materials were generated using custom scripts written primarily in R. The complete set of figure-generation scripts is available in the GitHub repository:

`https://github.com/pgarchive/data`

Within the repository, the folder supplementary_data/ contains the following files:

- scripts used to generate all manuscript figures
- scripts used to process intermediate analysis outputs
- configuration files used to plot the graphs.

These scripts allow for the full reproducibility of the plots presented in the manuscript and the supplementary materials.

#### TA8. Reproducibility and data availability

All the code used in this study are available in the project repository.

`https://github.com/pgarchive/data`

The repository includes the following:

- analysis scripts
- figure-generation scripts
- software configuration files
- documentation to reproduce the analyses.

The additional datasets used in this study are provided in Data S1–S3.
